# Recognition of highly branched N-glycans of the porcine whipworm by the immune system

**DOI:** 10.1101/2023.09.21.557549

**Authors:** Barbara Eckmair, Chao Gao, Akul Y. Mehta, Zuzanna Dutkiewicz, Jorick Vanbeselaere, Richard D. Cummings, Katharina Paschinger, Iain B. H. Wilson

## Abstract

Glycans are key to host-pathogen interactions, whereby recognition by the host and immunomodulation by the pathogen can be mediated by carbohydrate binding proteins, such as lectins of the innate immune system, and their glycoconjugate ligands. Previous studies have shown that excretory-secretory products of the porcine nematode parasite *Trichuris suis* exert immunomodulatory effects in a glycan-dependent manner. To better understand the mechanisms of these interactions, we prepared N-glycans from *T. suis* and both analyzed their structures and used them to generate a natural glycan microarray. With this array we explored the interactions of glycans with C-type lectins, C-reactive protein and sera from *T. suis* infected pigs. Glycans containing LacdiNAc and phosphorylcholine-modified glycans were associated with the highest binding by most of these proteins. In-depth analysis revealed not only fucosylated LacdiNAc motifs with and without phosphorylcholine moieties, but phosphorylcholine-modified mannose and N-acetylhexosamine-substituted fucose residues, in the context of maximally tetraantennary N-glycan scaffolds. Furthermore, O-glycans also contained fucosylated motifs. In summary, the glycans of *T. suis* are recognized by both the innate and adaptive immune systems, and also exhibit species-specific features distinguishing its glycome from those of other nematodes.

## Introduction

Nematodes or roundworms represent a large group of invertebrates whose species-specific habitats range from the soil through to the mammalian gut. A large number of nematodes are parasitic and their ability to often survive years despite recognition by host immune systems is remarkable, but probably reflects co-evolution of the immunomodulatory capacity of the nematode and the response of the host. The ability of parasites to shift the balance of the host immune system is due to a range of molecules, some of which are glycoproteins ^(*1*)^. The dichotomy between recognition as ‘foreign’ and a lack of full expulsion or eradication suggests that an expected immune response is misdirected or aborted. One example of this is the role of phosphorylcholine on nematode glycoconjugates; this zwitterionic moiety is well known as a component of phosphatidylcholine, a typical class of eukaryotic phospholipids and of platelet-activating factor, a related signalling molecule ^(*2*)^, but is also a widespread modification of glycoconjugates in non-vertebrate species, including lipopolysaccharides, N-glycans and glycolipids ^(*3,4*)^. In the case of bacterial infections in mammals, recognition of phosphorylcholine-modified glycoconjugates via C-reactive protein or anti-phosphorylcholine antibodies is associated with complement-mediated killing of the pathogen ^(*5*)^. In contrast, N-glycans carrying phosphorylcholine, exemplified by those on the ES-62 excretory-secretory protein of the rodent parasite *Acanthocheilonema viteae*, may be conformationally more flexible than bacterial polysaccharides; thus, despite binding of an ES-62/C-reactive protein complex to C1q, the full complement cascade response is not activated ^(*6*)^. On the other hand, ES-62 is just one example of a nematode protein with immunomodulatory effects ^(*7*)^; indeed many helminth parasites alter the balance between various T cell responses, so aiding parasite survival, but also affecting other aspects of the immune system ^(*8*)^.

Other than phosphorylcholine, N-glycans from nematodes are often fucosylated either on the core region or the antennae. One such motif, the so-called LDNF epitope (fucosylated LacdiNAc; GalNAcβ1,4[Fucα1,3]GlcNAc), is present in some nematodes ^(*9*)^ as well as schistosomes, which are trematode parasites ^(*10*)^; this motif is immunogenic in mammals ^(*11,12*)^, but is a mimic of the Lewis X epitope present on a range of mammalian glycans, including sialylated and sulphated forms recognized by selectins with roles in the immune system. In nematodes and lepidopteran insects ^(*13*)^, LDNF on N-glycans can also occur in a phosphorylcholine-modified form, as exemplified by structures we previously found in *Trichuris suis* ^(*14*)^, a porcine parasite having a negative effect on pig productivity and relative of the human parasite *T. trichiura*. Further modifications of nematode glycans include glucuronylation, fucosylation of the second (distal) core *N*-acetylglucosamine residue and galactosylation of core fucose residues ^(*9,15*)^. However, all these variations do not exist in a single organism and the glycan modifications can be differently combined resulting in a high degree of interspecies glycomic variability.

Considering the observed immunomodulatory effects, links have been made between reduced nematode infections and increased occurrence of allergies and autoimmunity in humans living within more developed societies ^(*16*)^. As a parasite that generally does not have a productive life-cycle in humans, *T. suis* has attracted attention as a potential unorthodox therapy for such diseases and its eggs have been administered to patients with Crohn’s disease and autoimmune rhinitis as well as to animals with experimental autoimmune encephalomyelitis ^(*17–19*)^. While the effects and safety of such therapies are controversial ^(*20*)^, it is clear that *T. suis* excretory-secretory products do exert typical immunomodulatory effects, which are partly glycan-dependent ^(*21*)^. With this background and to investigate interactions of *T. suis* glycans with components of mammalian immune systems, we have established the first microarray of natural glycans of this species, accompanied by an in-depth study of its N-glycomic capacity. Thereby, we show that especially the larger N-glycans bind mammalian lectins as well as antibodies present in the sera of infected pigs.

## Results

### A natural T. suis N-glycan array

As a means for understanding glycan-dependent immunomodulatory effects of *T. suis* excretory-secretory products, we undertook the construction of a natural N-glycan array. Enymatically released glycans were non-reductively labelled with FMAPA, a fluorescent methoxyamino-based linker developed in the Cummings’ laboratory, and then separated by HPLC and analysed by MALDI-TOF MS/MS (***Figure 1*** and ***Supplementary Figures 1-4***); prior to printing the glycans are treated with piperidine, which removes the fluorescent Fmoc moiety and reveals a free amino group which can be printed onto NHS-modified glass slides ^(*22*)^. In total twenty-seven *T. suis* glycan fractions, five control compounds and a ‘buffer only’ set of spots were printed in each field.

**Figure 1.**
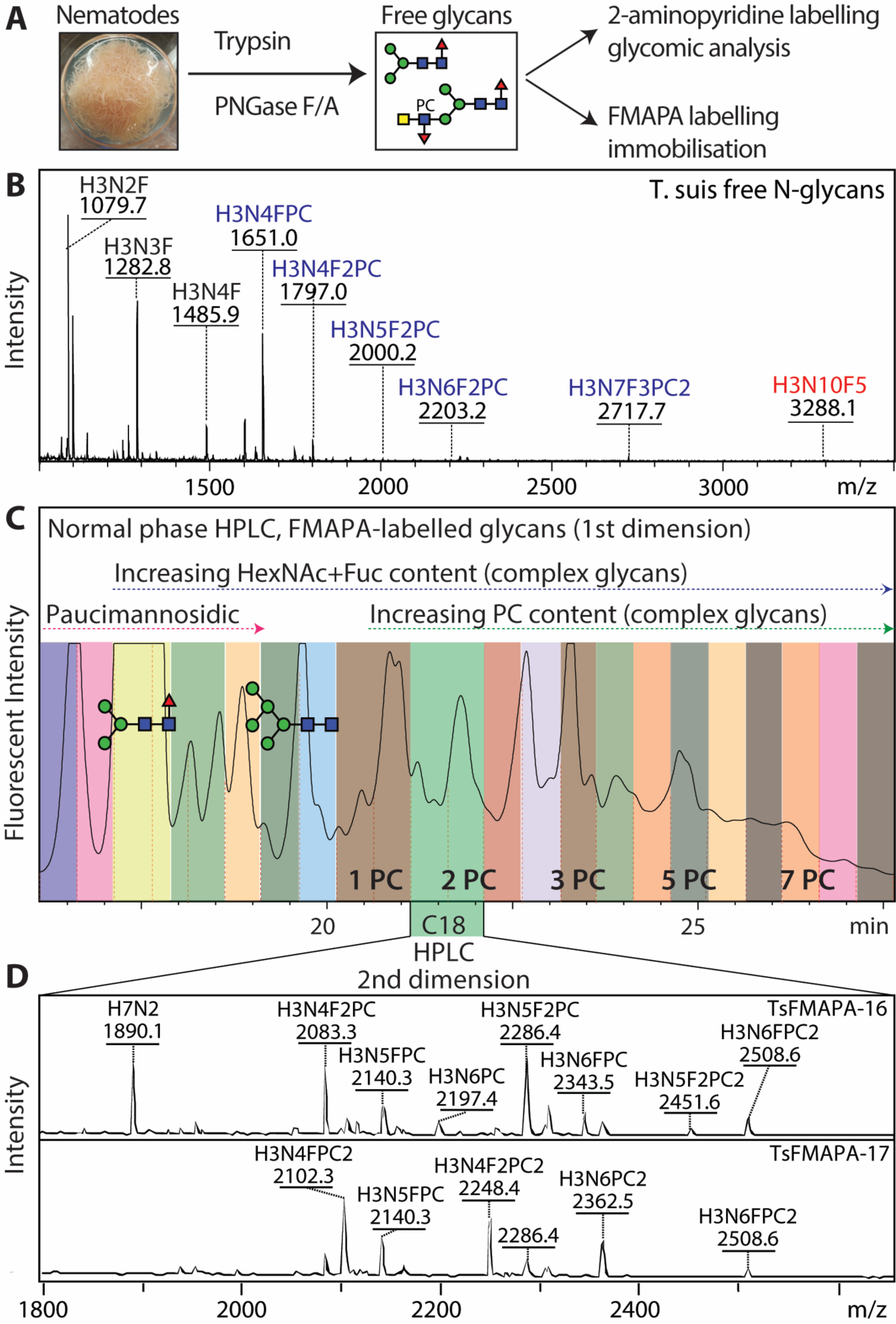
Glycomic analysis of *T. suis* N-glycans. (A and B) N-glycans were released from tryptic peptides and analysed by MALDI-TOF MS; 90% were labelled with FMAPA for preparation of glycan arrays, the remaining 10% with 2-aminopyridine for in-depth glycomic analysis (see ***Figures 3-7*** and ***Supplementary Figures 7-17***). (C) The FMAPA-labelled glycans (1600 nmol) were fractionated by semi-preparative normal phase HPLC (Luna NH2), annotations of a pauci- and an oligomannosidic structure are on the basis of MALDI-TOF MS data; the elution positions for complex glycans with increasing numbers of phosphorylcholine residues are also indicated as PC1 – PC7. Selected pools were subject to a second dimension on a reversed-phase column. (D) The resulting 27 pools, each enriched in a subset of glycans (see mass spectra for examples indicating that each printed fraction contained a number of structures with phosphorylcholine-modified glycans detected as [M+H]^+^ and other glycans as [M+Na]^+^), were treated with piperidine prior to array printing (see ***Figure 2*** and ***Supplementary Figure 5*** for results). For further mass spectra of FMAPA-labelled glycans refer to ***Supplementary Figures 1-4***.

### Standard lectins

an initial validation of the array was performed using a number of widely-used lectins binding a range of glycan motifs (***Figure 2A***; see also ***Supplementary Figure 5*** for data). A majority of *T. suis* glycan fractions were bound by the ‘fucose-specific’ *Aleuria aurantia* lectin (AAL) and the ‘mannose- and simple complex-specific’ Concanavalin A (ConA; most fractions 1-20); many of the fractions recognized by the latter were also bound by the ‘mannose-specific’ *Galanthus nivalis lectin* (GNA; fractions 1-11, except 10). On the other hand, only a few fractions, concluded to contain glycans with unsubstituted LacdiNAc motifs (B fragments of *m/z* 407, i.e., HexNAc_2_), were recognized by wheat germ or *Wisteria floribunda* agglutinins (WGA and WFA; e.g., 9 and 14-16). Binding by *Lotus tetragonolobus* lectin (LTL) is most obvious for the largest glycans displaying fucosylated antennae (fractions 23-26), whereas recognition by *Ricinus communis* (RCA) or *Sambucus nigra* (SNA) lectins to *T. suis* glycans was relatively weak, which correlates with the glycomic analyses (see below) indicating a lack of antennal galactose or sialic acid.

**Figure 2.**
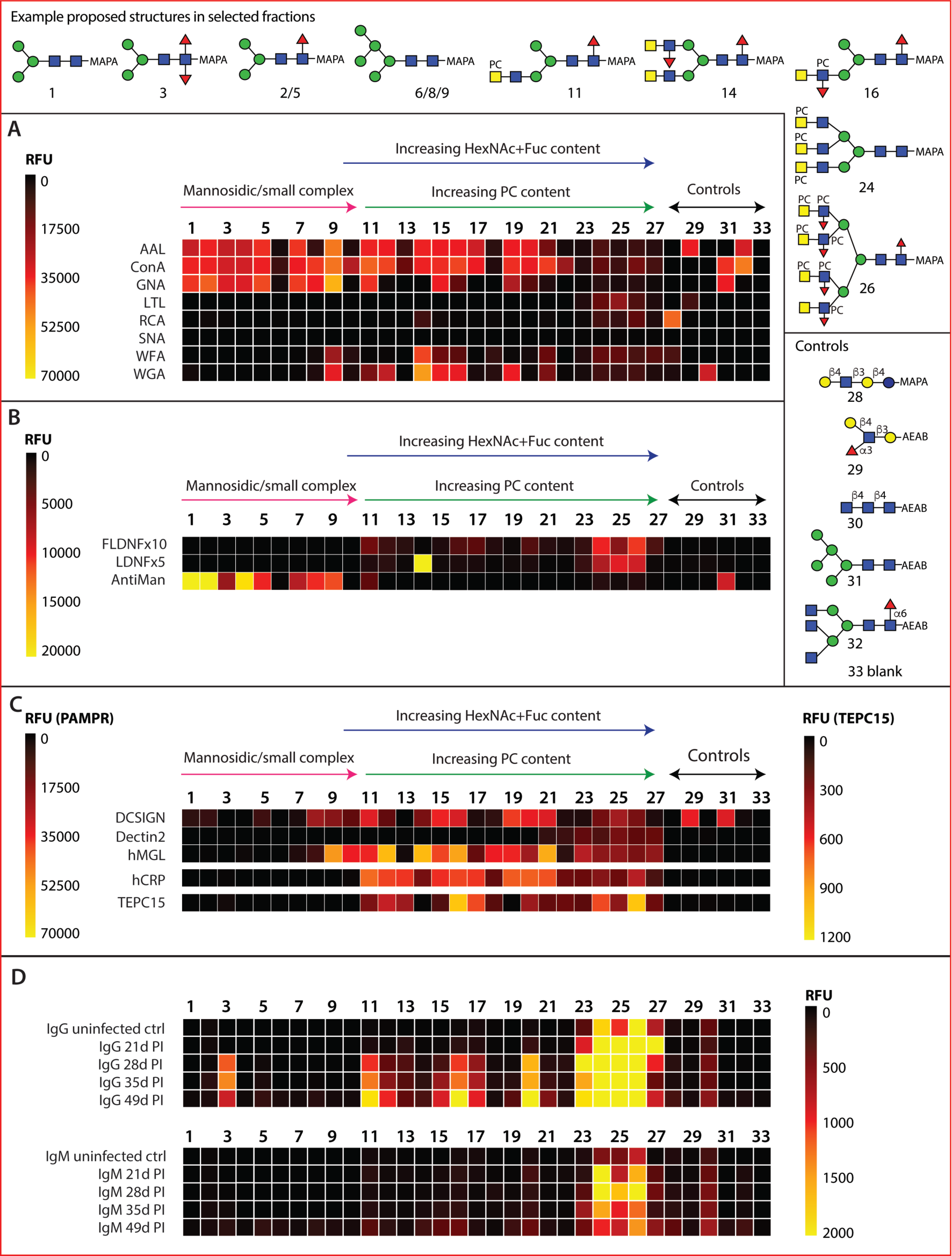
Array analysis of *T. suis* N-glycans. Twenty-seven HPLC fractions of F-MAPA-labelled natural N-glycans derived from *T. suis* (1–27) in addition to five controls (28–32) and a buffer control (33) were printed and incubated with a variety of (A) commercial lectins, (B) anti-glycan antibodies, (C) pathogen-associated molecular pattern receptors and (D) sera from infected pigs. The data are shown as heatmaps representing the relative fluorescence units (RFUs) following the key in each panel; for using a comparable scale for the anti-glycan antibodies, the values for anti-FLDNF (F2D2) and anti-LDNF (L6B8) are multiplied by ten- or five-times. Example proposed structures for the major components in selected fractions and the pure control glycans are shown according to the Symbol Nomenclature for Glycans (refer also to ***Supplementary Figures 1-4*** for relevant HPLC and MS data and ***Supplementary Figure 5*** for bar chart representations).

### Anti-glycan antibodies

Three monoclonal antibodies originally raised against schistosomal antigens, with reported binding to mono/difucosylated LacdiNAc or paucimannosidic structures, were used to probe the array (***Figure 2B***). The LDNF antibody bound especially to fraction 14, which contains glycans with probable fucosylated LacdiNAc motifs (B fragments of *m/z* 553, i.e., HexNAc_2_Fuc_1_); a lower degree of binding was also observed for the LDNF and FLDNF antibodies to higher molecular weight fractions 24-26 containing glycans with putative LacdiNAc motifs decorated with fucose and phosphorylcholine, as judged by B fragments of *m/z* 515 and 718 (HexNAc_1-2_Fuc_1_PC_1_). The anti-mannose monoclonal had a specificity rather similar to that of GNA, binding best to fractions containing Man_3-5_GlcNAc_2_Fuc_0-1_ as well as the Man_5_GlcNAc_2_ control.

### Innate immune lectins

Three human lectins, including two known to bind *T. suis* soluble products ^(*21*)^, were used to probe the array (***Figure 2C***). DC-SIGN appeared to bind especially fractions containing oligomannosidic glycans (Man_5-9_GlcNAc_2_), overlapping with those bound by ConA. Those fractions recognized by macrophage galactose lectin (MGL) partly overlapped with those of WGA and C-reactive protein (CRP), suggesting that LacdiNAc modified glycans (with or without phosphorylcholine, defined on the basis of *m/z* 407 and 572 B fragments) were ligands. Dectin-2 displayed relatively low binding rather to the fractions containing the largest N-glycans.

### Phosphorylcholine binding proteins

CRP and the TEPC-15 IgA monoclonal are well established as being specific for phosphorylcholine, a typical modification of nematode glycans. Here, this specificity is verified (***Figure 2C***), as fractions rich in glycans with *m/z* 369 (HexNAc_1_PC_1_) and related B fragments were recognised, but there are subtle differences. CRP bound a range of fractions containing glycans predicted to contain one or two phosphorylcholine residues, but relatively less well to those with multiple phosphorylcholine-modified antennae. TEPC-15 binding to various fractions overlapped, but was lower in terms of absolute fluorescence intensity.

### Pig infection sera

IgG and IgM from pigs infected with *T. suis* bound especially to glycans present in a limited number of fractions (***Figure 2D***), especially those with the highest molecular weight glycans (2500-5000 Da) carrying multiple phosphorylcholine residues as well as to fraction 3 containing a core difucosylated N-glycan. Thereby, the trend is that the antibodies, only partly present prior to infection, bind to those glycan fractions with lower binding to DC-SIGN, MGL and CRP; on the other hand, the pig IgG and IgM bind most to fractions 23-26 which are also recognized by the anti-FLDNF, anti-LDNF and anti-PC monoclonal antibodies.

### Analysis of an extended range of T. suis N-glycans

MALDI-TOF MS of the F-MAPA-labelled N-glycans indicated the presence of highly complex structures, but their separation was suboptimal despite two rounds of HPLC; this meant their fragmentation was partly ambiguous due to co-eluting structures. Thus, we undertook a more detailed glycomic analysis of *T. suis*. Previously, we analysed its N-glycome using one adult individual and observed, in addition to a plentiful paucimannosidic Man_3_GlcNAc_2_Fuc_1_ structure, a number of minor glycans carrying one or two antennae modified with LacdiNAc, fucose and/or phosphorylcholine residues ^(*14*)^. In the present study, we first employed MALDI-TOF MS of permethylated N-glycans; this did indicate the occurrence of structures with multiple fucosylated LacdiNAc motifs (***Supplementary Figure 6***), but phosphorylcholine-modified glycans cannot be analysed by this method as they are lost during derivatization. Therefore, we labelled the glycans with 2-aminopyridine and performed off-line 2D-HPLC-MALDI-TOF MS as in a previous study on *Dirofilaria* ^(*15*)^; the resulting analyses showed an immense glycomic variety, including many structures predicted to contain multiple phosphorylcholine residues. Due to the ambiguities when predicting structure from mass spectral data, 2D-HPLC (first hydrophilic interaction, then reversed-phase; ***Supplementary Figure 7*** and ***Figure 3***) was performed for many glycan fractions prior to MALDI-TOF MS/MS complemented by chemical and enzymatic treatments. The typical paucimannosidic and oligomannosidic glycans again dominated the glycome ^(*14*)^ with about 60% being Man_3_GlcNAc_2_Fuc_1_ as judged by fluorescence; these glycans were analysed, but the data are only summarised as they have been found in many other species (***Supplementary Figures 7 and 8***). However, compared to our previous study with limited material, we could now detect larger structures of up to 3500 Da and above, including tri- and tetra-antennary glycans with various categories of antennal motif (***Figure 3*** and ***Supplementary Figure 7***).

**Figure 3.**
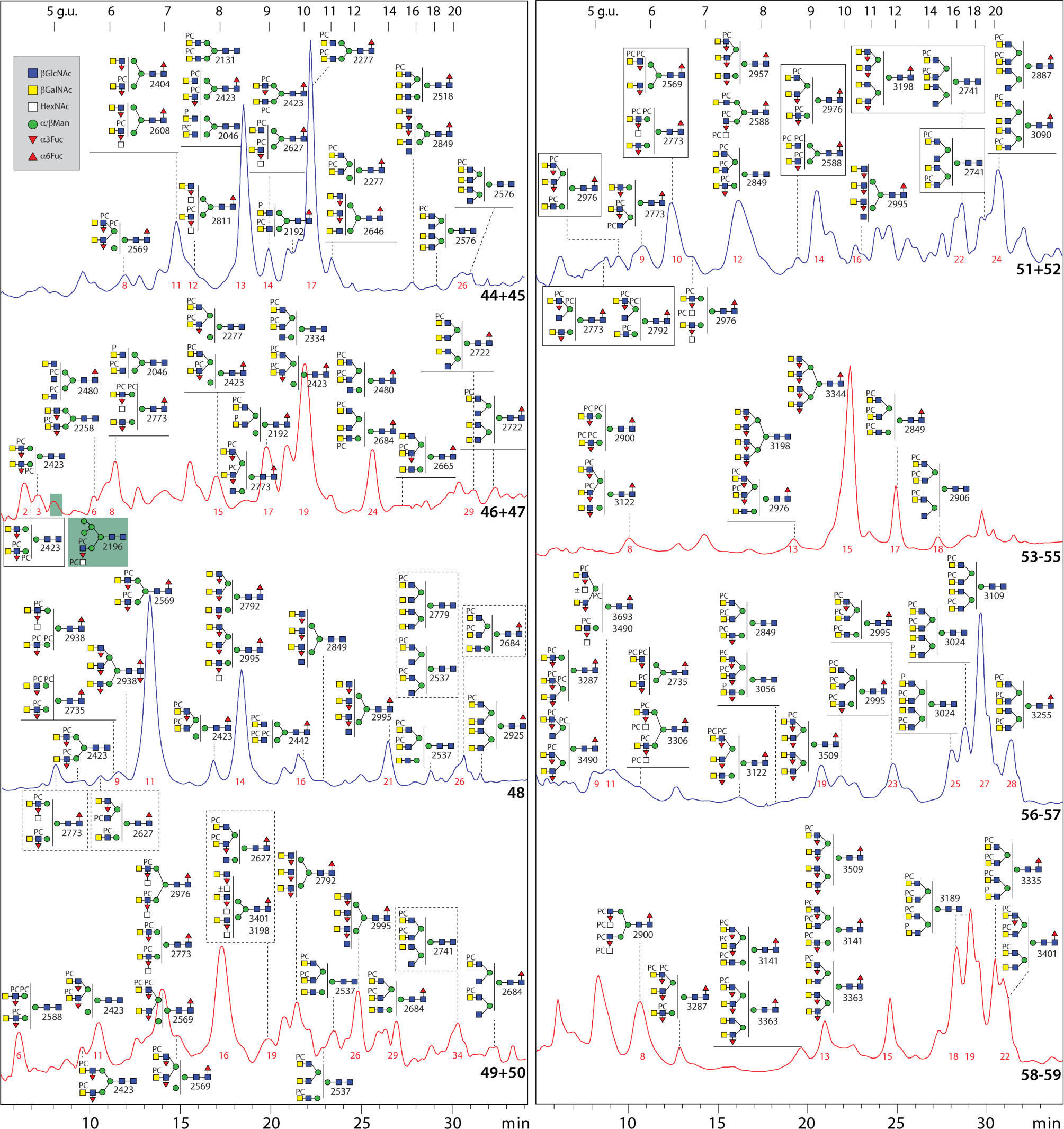
2D-HPLC analysis of *T. suis* N-glycans. Pyridylamino-labelled N-glycans derived from *T. suis* were fractionated by hydrophilic interaction chromatography (***Supplementary Figure 7***); selected fractions were pooled and subject to RP-amide HPLC prior to MALDI-TOF MS/MS before and after chemical and enzymatic treatments (for examples, see ***Figures 4-7*** and ***Supplementary Figures 8-17***). The RP-amide column was calibrated in terms of glucose units (5-20 g.u.). Proposed structures are shown according to the Symbol Nomenclature for Glycans (see also key in grey box) and are listed in ***Supplementary Table 2***; P, phosphate; PC, phosphorylcholine. The ability of RP-HPLC to resolve isomers of pyridylaminated N-glycans, as demonstrated by their different fragmentation patterns, has also been shown in previous studies on other nematode glycomes ^(*15,37*)^. Selected fraction numbers are shown in red.

### LacdiNAc and fucosylated LacdiNAc

MS/MS of various glycans yielded B-fragments of *m/z* 407 and 553 (HexNAc_2_Fuc_0-1_; ***Figure 4***), which are commonly observed in various invertebrates; the simplest examples of glycans with these motifs (*m/z* 1395, 1541, 1687 and 1744; Hex_3_HexNAc_4-5_Fuc_0-2_) were previously reported ^(*14*)^. Such motifs can be based on LacdiNAc (GalNAcβ1,4GlcNAc) or chitobiose (GlcNAcβ1,4GlcNAc). Fortunately, the GalNAc-specific *C. elegans* HEX-4 hexosaminidase can distinguish the two motifs, while *S. plicatus* chitinase does not ^(*15*)^. Based on use of these two exoglycosidases, it could be determined that the HexNAc_2_ motifs in *T. suis* are solely LacdiNAc (for example digests, see ***Supplementary Figure 9 A and B***), as compared to the chitobiose or chitotriose motifs found in a variety of other nematodes ^(*15,23,24*)^. The fucose residue was sensitive to hydrofluoric acid or almond α1,3/4-fucosidase ^(*25,26*)^, whereby it is assumed that the antennal fucose residues are α1,3-linked (see examples for *m/z* 1687; ***Supplementary Figure 9B***), as previously concluded ^(*14*)^.

**Figure 4:**
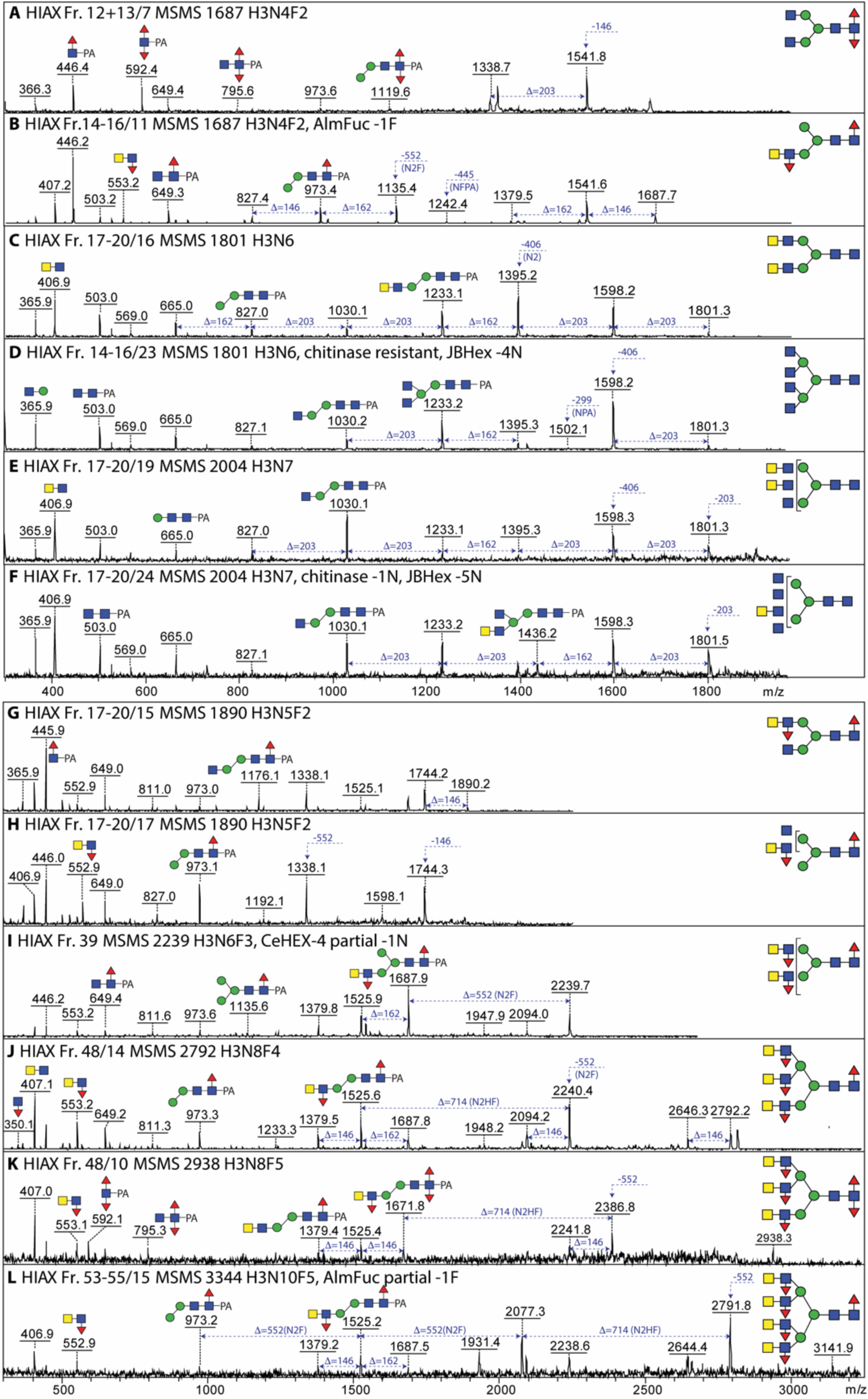
Isomeric variations of LacdiNAc-modified and fucosylated N-glycans. MALDI-TOF MS/MS of selected N-glycans with LacdiNAc and fucose modifications from different 2D (HIAX followed by RP-amide) or 1D (HIAX only) fractions. Shown are isomers of *m/z* 1687 (A, B), 1801 (C, D), 2004 (E, F) and 1890 (G, H) as well as single structures of between *m/z* 2239 and 3344 (I-L). Core difucosylated structures display the signature *m/z* 592 Y1 ion (A and K), whereas LacdiNAc and antennal fucosylated LacdiNAc are characterized respectively by *m/z* 407 and 553 B2 ions. Fractions are named with the HIAX pool number followed, if relevant, by the RP-amide second dimension fraction number (see ***Figure 3*** or ***Supplementary Figures 7 and 8***), the *m/z* and the abbreviated composition of the form H3NxFy for Hex_3_HexNAc_4-10_Fuc_0-5_. Some fractions were also incubated with exoglycosidases with the noted resistance or sensitivity.

### Difucosylated core GlcNAc

We have previously observed that *T. suis* has a simple α1,3/6-difucosylated N-glycan identical to one found in a large range of invertebrates, specifically Man_3_GlcNAc_2_Fuc_2_ yielding a characteristic *m/z* 592 Y-fragment upon MS/MS ^(*14*)^ as also observed for one isomer of *m/z* 1687 (***Figure 4A***); however, also a limited number of larger glycans displayed this modification, including a triantennary glycan of *m/z* 2938, featuring both *m/z* 553 (B-ion) and *m/z* 592 (Y-ion) fragments characteristic of having both fucosylated LacdiNAc and core difucosylation (***Figure 4K***).

### Phosphorylcholine- and phosphate modified LacdiNAc

Glycans in many nematodes, as well as example cestode and lepidopteran species, are modified with the zwitterion phosphorylcholine 6-linked via phosphodiesters to *N*-acetylhexosamine residues ^(*13,27–29*)^. We have previously reported simple examples from *T. suis* ^(*14*)^, but here we find a very large range of phosphorylcholine-modified structures with the signature *m/z* 369 (PC_1_HexNAc_1_) B-ion fragment as well as examples with *m/z* 515, 572 and 718 (PC_1_HexNAc_1-2_Fuc_0-1_; ***Figure 5*** and ***Supplementary Figures 9C, 10-17***). The pattern of MS/MS fragments of PC-modified LacdiNAc and fucosylated LacdiNAc was indicative of two positions for the PC moieties. Often the *m/z* 369 fragment or the loss of 368 Da was very dominant, shown also by terminal loss of a PC_1_HexNAc_1_ unit; in other cases, *m/z* 572 was more intense. In the case of fucosylated antennae, the occurrence of *m/z* 369, 515 (PC_1_HexNAc_1_Fuc_1_) and 718 (PC_1_HexNAc_2_Fuc_1_) indicated either that the terminal HexNAc was PC-modified and the subterminal HexNAc substituted with fucose (see, e.g., the *m/z* 1852, 2055 and 2423 glycans in ***Figure 5 A, C and F, Supplementary Figure 10E and 13A***) or that the subterminal HexNAc was substituted with both phosphorylcholine and fucose (see, e.g., the *m/z* 1852 and 2423 glycans in ***Figure 5 B, D and E*** and ***Supplementary Figure 10A***). The antennal phosphorylcholine and fucose residues were, in either case, sensitive to hydrofluoric acid treatment, known to cleave phosphodiester bonds ^(*30*)^ as well as α1,3-fucose; the underlying LacdiNAc motifs were digested with either chitinase or *C. elegans* HEX-4 (***Supplementary Figures 10 and 13***).

**Figure 5:**
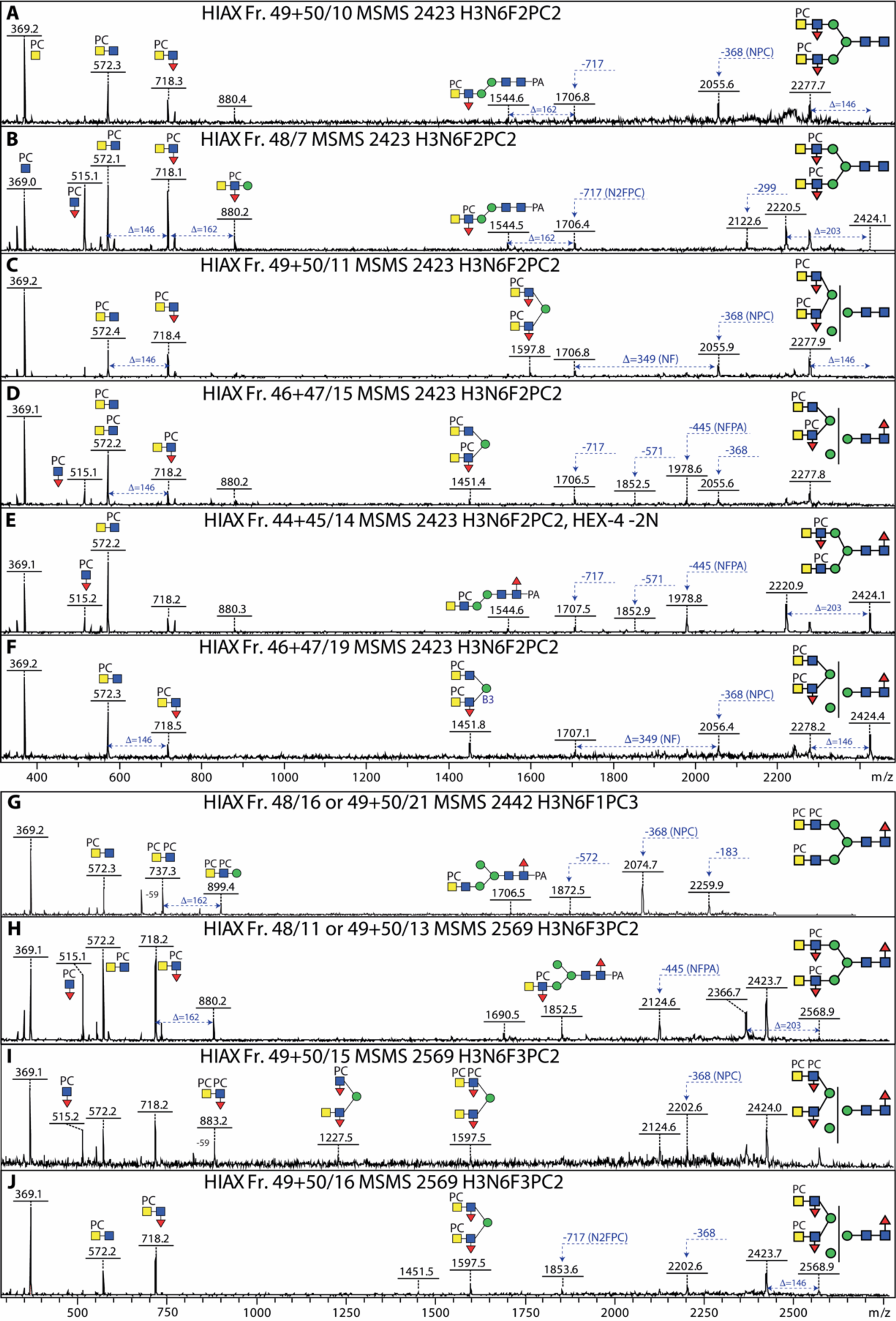
Isomeric variations of phosphorylcholine-modified N-glycans. MALDI-TOF MS/MS of selected N-glycans with phosphorylcholine modifications from different 2D fractions (HIAX followed by RP-amide; see ***Figure 3***). Shown are isomers of *m/z* 2423 (A-F) and 2569 (H-J), as well as one structure of *m/z* 2442 (G). The pattern of B1 and B2 ions (ratios of *m/z* 369 and 572 or *m/z* 515 and 718) as well as the loss of either 203 or 368 Da is indicative for the position of the phosphorylcholine on LacdiNAc and fucosylated LacdiNAc (either on the terminal GalNAc or subterminal GlcNAc) or whether both zwitterionic residues are on the same antenna (*m/z* 737 and 883). For further examples of phosphorylcholine-modified N-glycans refer to ***Supplementary Figures 9-17***.

Due to the 2D-HPLC fractionation, many isomeric forms could be detected; for instance, hybrid or pseudohybrid structures with two antennae on the same α-mannose (e.g., with key B3 fragments of *m/z* 1140, 1305, 1451 and 1597; Hex_1_HexNAc_4_Fuc_0-2_PC_1-2_) or bi- and triantennary structures with one or two antennae on each of the core α-mannose residues were detected (***Figure 5*** and ***Supplementary Figures 14, 16 and 17***) as well as the aforementioned tetra-antennary glycans (***Supplementary Figure 11***). Some structures carrying phosphorylcholine were isobaric with heavily fucosylated ones, e.g., glycans of *m/z* 2995 have compositions of either Hex_3_HexNAc_8_Fuc_2_PC_3_ and Hex_3_HexNAc_9_Fuc_4_ as distinguished by retention time and fragmentation (***Supplementary Figure 14 G-J***); for other compositions up to ten isomers were detected, e.g., Hex_3_HexNAc_6_Fuc_2_PC_2_ and Hex_3_HexNAc_7_Fuc_3_PC_2_ (*m/z* 2423 and 2773; ***Figure 5 A-F*** and ***Supplementary Figure 14 D-F***).

In a few cases, *m/z* 737 or 883 fragments were observed, which would be indicative of a LacdiNAc motif, with or without fucose, modified with two phosphorylcholine residues (see *m/z* 2442 and one isomer of *m/z* 2569 in ***Figure 5G and I***). In addition, some glycans had an 80 Da modification, which was also observed in negative mode and was sensitive to hydrofluoric acid: thus, it was concluded that these glycans carry a phosphate residue rather than a phosphorylcholine (***Supplementary Figure 10E*** and ***Supplementary Figure 17 A and B***); in some cases, the glycans had one antenna modified with phosphate and others with phosphorylcholine (***Supplementary Figure 17 D, F, H and J***). This combination has not been previously observed in nematodes.

### Substituted antennal fucose residues

In a number of nematodes, such as *Dirofilaria*, HexNAc_3_Fuc_1_PC_0-1_ motifs can be detected based on the relevant *m/z* 756 and 921 Y fragments, which upon hydrofluoric acid treatment is replaced by one at *m/z* 610 (HexNAc_3_). Puzzlingly, in the case of *T. suis*, no HexNAc_3_ motifs were observed upon this chemical defucosylation treatment – rather only *m/z* 407 (HexNAc_2_) motifs, even in the case of large glycans with eight or more HexNAc residues and multiple fucose and phosphorylcholine modifications; furthermore, these digests showed that a HexNAc and a fucose were removed at once (***Supplementary Figures 9A and 13E***). Thus, as depicted in ***Figure 6*** and ***Supplementary Figures 14-16***, we assume that the antennal fucose residues can be substituted by an *N*-acetylhexosamine, which is in turn occasionally modified by phosphorylcholine (*m/z* 921 and 1087 HexNAc_3_Fuc_1_PC_1-2_ B-ions). Whereas many examples of the substituted fucose are in the context of LacdiNAc units, others lack the GalNAc of the LacdiNAc; as judged by the lack of *m/z* 572 and 718 B-ions, but the presence of *m/z* 515 and 883 (HexNAc_1-2_Fuc_1_PC_1_) fragments, they are concluded to carry PC-modified HexNAc directly linked to the antennal fucose (***Figure 6 G and H***). To date, these specific combinations of modifications have not been observed in other organisms.

**Figure 6:**
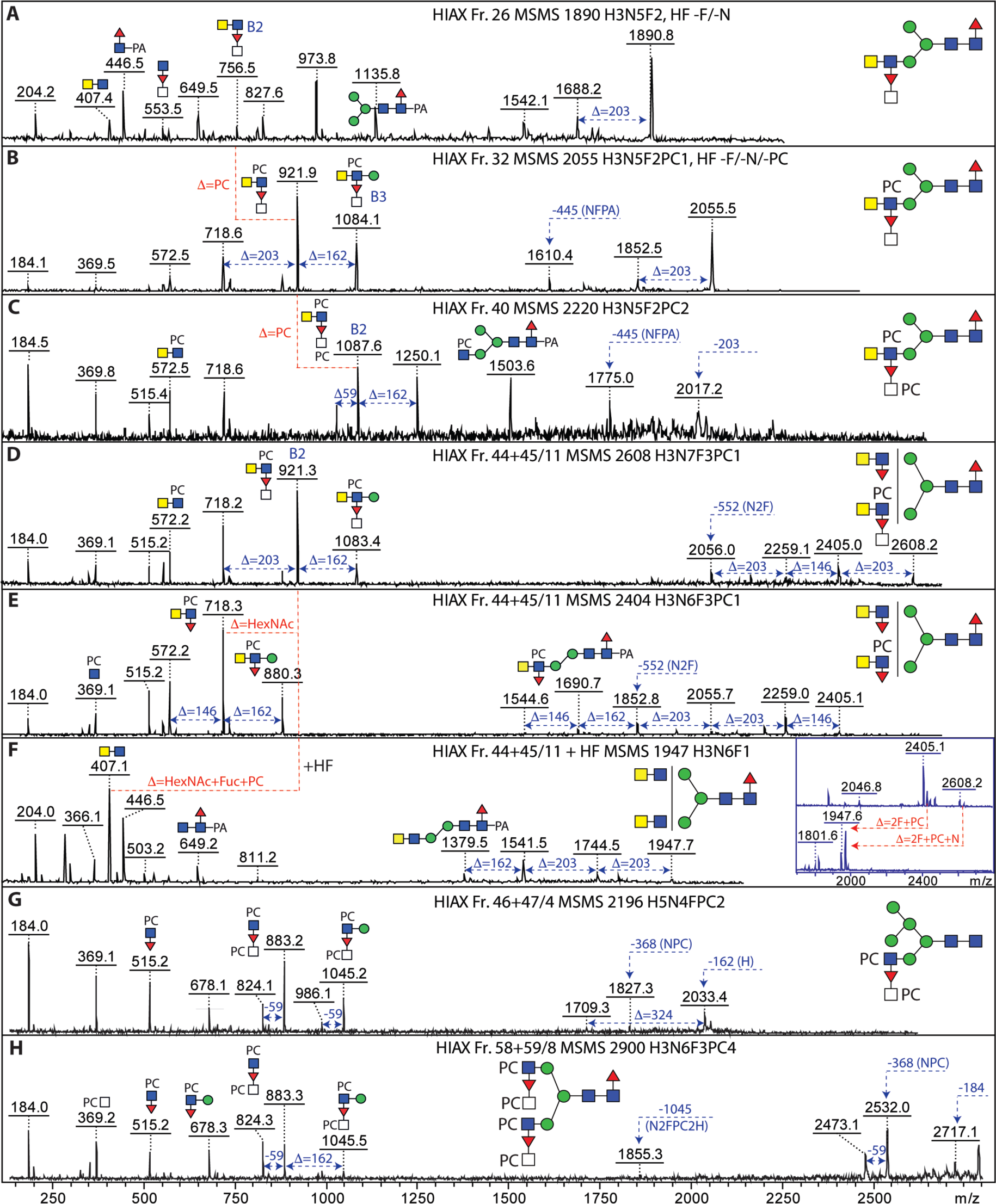
N-glycans with antennal N-acetylhexosamine-substituted fucose residues. MALDI-TOF MS/MS of selected N-glycans in different 1D- or 2D-HPLC fractions (see ***Supplementary Figure 7*** and ***Figure 3***) with fucosylated LacdiNAc modifications carrying further N-acetylhexosamine and phosphorylcholine residues. The simplest hybrid structures with substituted fucose residues (A-C) show differences in the numbers of phosphorylcholine residues as indicated by the B2 fragments at *m/z* 756, 921 and 1087; HF treatment of the *m/z* 1890 and 2055 glycans (***see Supplementary Figures 9 and 13***) resulted in loss of the FucHexNAc unit. The co-eluting biantennary structures (D-F) of *m/z* 2404 and 2608 were both digested down to *m/z* 1947 with HF (see inset) accompanied by loss of the B2 ions at *m/z* 718 and 921, indicative of the loss of a FucHexNAc unit from the latter. (G and H) PCHexNAc substitution of antennal fucose in the context of hybrid and biantennary glycans; for further examples, see also ***Figure 7*** and ***Supplementary Figures 14-16***.

### Phosphorylcholine-modified core mannose

Amongst the various glycans modified with phosphorylcholine, some yielded MS/MS fragments indicative of modification of the core trimannosylchitobiosyl region (*m/z* 1154 and 1300, i.e., 165 Da in addition to 989 or 1135; see ***Figure 7***). Also, mass differences of 327 Da (Hex_1_PC_1_) rather than the usual 368 Da (HexNAc_1_PC_1_) in addition to the corresponding putative B1-ions at *m/z* 328 and 369 were observed; furthermore, B3-ions at *m/z* 1045 or 1597 (HexNAc_2_Hex_1_Fuc_1_PC_2_ or HexNAc_4_Hex_1_Fuc_2_PC_2_; ***Figure 7*** and ***Supplementary Figure 16 F, H and I***) suggested that there were phosphorylcholine modifications of core α-mannose residues to which PC-modified fucosylated LacdiNAc units were attached. HexNAc-substituted antennal fucose was also present on some glycans with PC-modified mannose (***Figure 7 F and G***).

**Figure 7:**
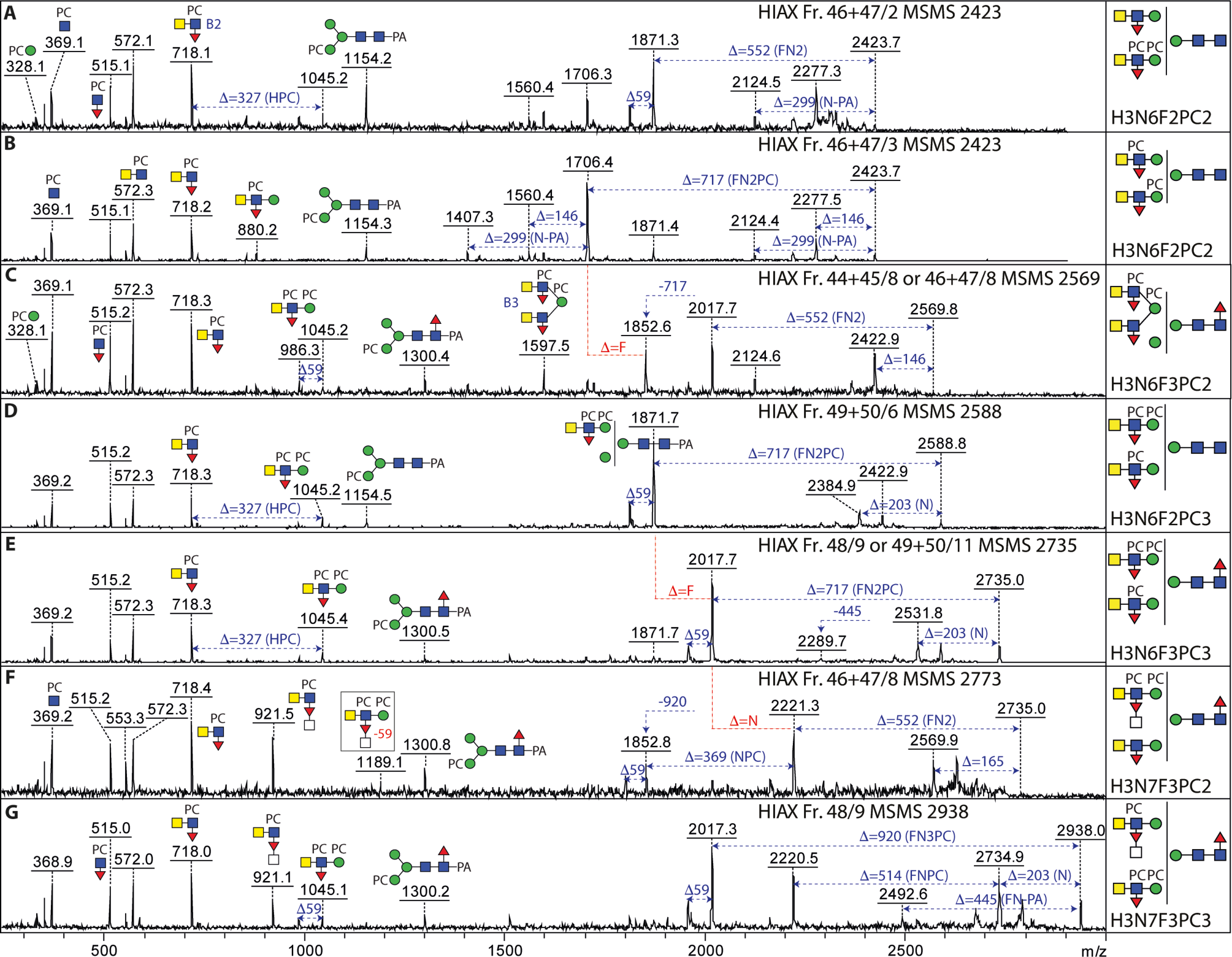
N-glycans with phosphorylcholine modifications of mannose and fucosylated LacdiNAc motifs. MALDI-TOF MS/MS of selected N-glycans in different 2D-HPLC fractions (see ***Figure 3***) with multiple phosphorylcholine modifications. (A-G) While the typical PC-modified fucosylated LacdiNAc motif (B2; *m/z* 718), also in substituted form (*m/z* 921), shows the occurrence of antennal phosphorylcholine, putative Y fragments at m/z 1154 and 1300 are compatible with a PC-modification of a core α-mannose residue. Furthermore, B fragments at *m/z* 986, 1045 or 1189 as well as *m/z* 328 are also indicative of PCMan motifs; the occurrence of two phosphorylcholine residues on juxtaposed residues results in the formation of ions 59 Da smaller than the true B ion. The *m/z* 2938 glycan (panel G) is isobaric to the triantennary pentafucosylated glycan shown in ***Figure 4K*** and further isomers of m/z 2423, 2569 and 2773 (A-C and F) are shown in ***Figure 5*** and ***Supplementary Figure 14*.**

### O-glycome

β-Elimination followed by permethylation was employed to examine the O-glycans of *T. suis*. Whereas some structures were reminiscent of the hexose-modified structures from *C. elegans* (e.g., *m/z* 942; Hex_4_HexNAc_1_), others were based on multiple HexNAc residues and could also be fucosylated (***Figure 8***); potentially, the composition could correspond to a fucosylated LacdiNAc-type structure as proposed for some O-glycans from *Haemonchus contortus* ^(*31*)^. As the glycans were permethylated, it is not possible to determine whether they were modified with phosphorylcholine, as recently found for some O-glycans from *C. elegans*.

**Figure 8:**
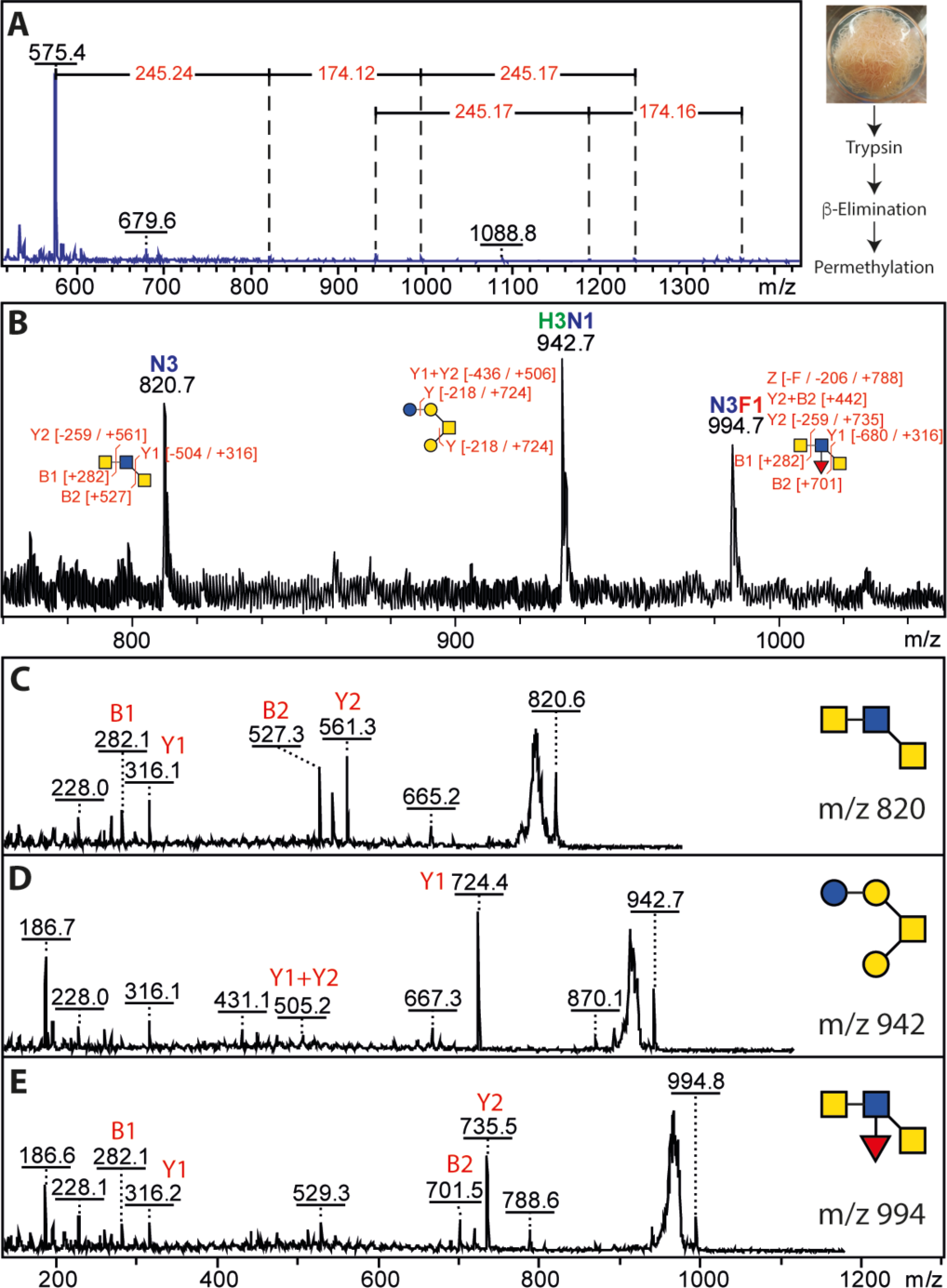
MALDI-TOF MS analysis of permethylated O-glycans. **(A)** Glycans released by reductive β-elimination were permethylated and analysed by MALDI-TOF MS showed a potential series of up to *m/z* 1362 can be interpreted as being based on HexNAc and Fuc modifications of HexNAc_2_ and Hex_3_HexNAc_1_ structures. **(B-E)** The *m/z* range 800-1000 showed three glycans with interpretable MS/MS.

### Phylogenetic analyses

In order to understand the N-glycan branching patterns, the existing *T. suis* genomic data were searched for homologues of *N*-acetylglucosaminyltransferases involved in eukaryotic N-glycan biosynthesis. Unlike *C. elegans* which possesses three GlcNAc-TI, one GlcNAc-TII and one GlcNAc-TV homologues, respectively GLY-12, −13, −20 and −2 ^(*32*)^, and *Trichinella spiralis* which is predicted to have one isoform each of GlcNAc-TI, GlcNAc-TII, GlcNAc-TIV and GlcNAc-TV, *T. suis* is predicted to lack any GlcNAc-TV (***Supplementary Figure 18***), which may account for the different isomeric form of the tri- and tetra-antennary N-glycans as compared to other nematodes (***Supplementary Figure 8***).

### Glycoprotein analyses

Lectin blotting of *T. suis* lysates indicated binding to ConA, AAL, DC-SIGN, MGL and Dectin-2, which was generally reduced upon pre-incubation with PNGase F (***Figure 9***), a finding compatible with the array data. A rather dominant band at 46 kDa was observed and excised from the corresponding Coomassie-stained gels prior to tryptic peptide mapping and glycan analysis. The best match for the proteomic data was to an “uncharacterized” *T. suis* protein related to a secreted Poly-Cysteine and Histidine-Tailed Metalloprotein from *Trichinella spiralis*, which is also N-glycosylated ^(*33*)^. Homologous proteins, also known as p43, have also been identified in *Trichinella pseudospiralis* adult secretome ^(*34*)^, *Trichuris muris* excretory-secretory products ^(*35*)^ and *Trichuris trichuria* egg and female extracts ^(*36*)^. In terms of N-glycomics of the protein band, a typical range of pauci- and oligomannosidic glycans as well as structures carrying LacdiNAc residues with or without phosphorylcholine were revealed upon MALDI-TOF MS/MS of the HPLC-fractionated N-glycans, thereby representing a subset of the overall N-glycome.

**Figure 9:**
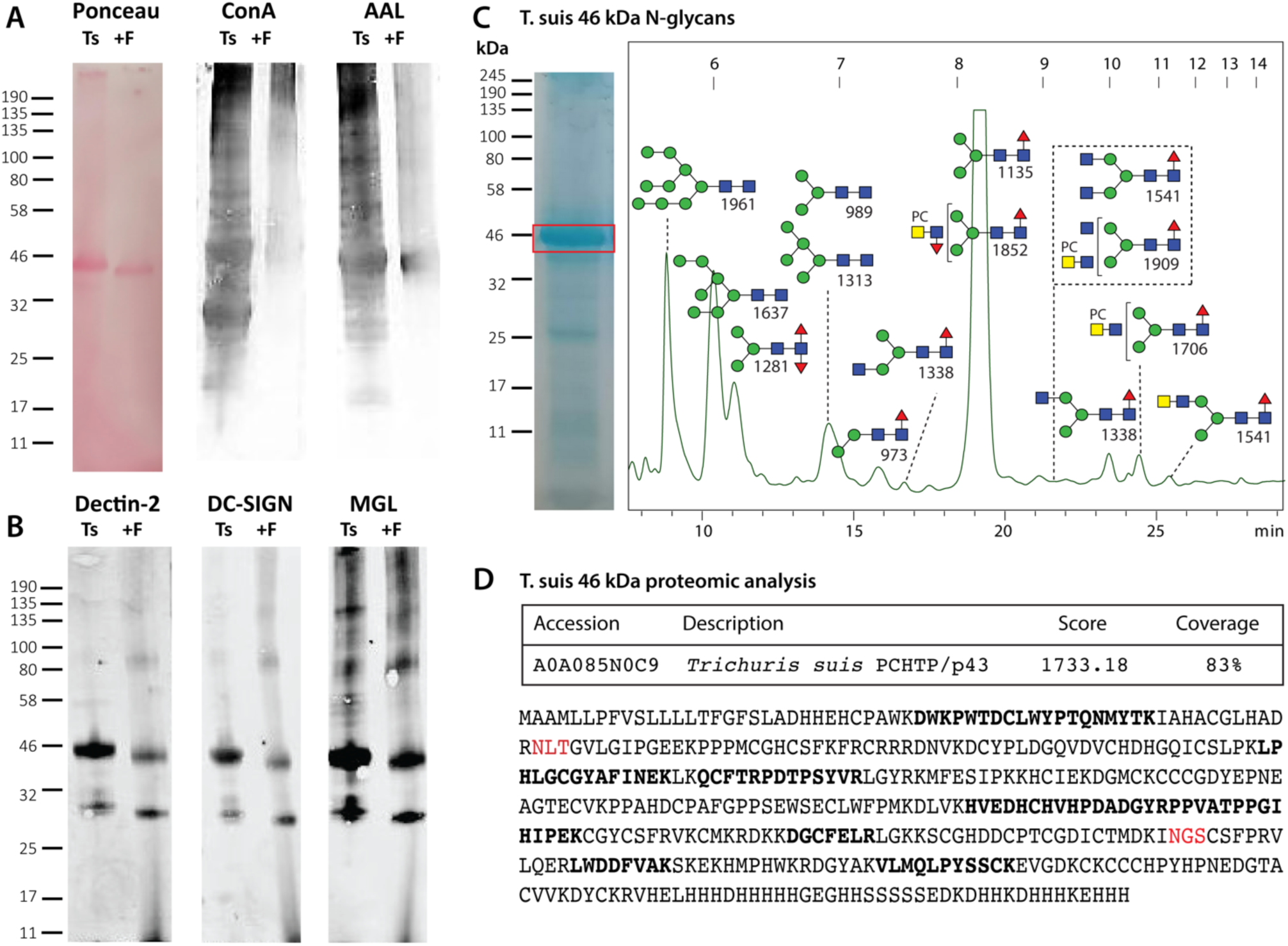
Western blotting and glycomic analysis of the major *T. suis* glycoprotein. **(A)** *T. suis* extract was subject to SDS-PAGE followed by Western blotting with Concanavalin A and *Aleuria aurantia* lectin before and after PNGase F treatment; Ponceau S staining indicates a shift in the molecular weight of the major protein upon PNGase F digestion. **(B)** Western blotting with three innate immune system proteins, Dectin-2, DC-SIGN and MGL showing partial loss of staining after PNGase F treatment. **(C)** Coomassie blue staining of SDS-PAGE indicating the excised band of 46 kDa, which was subject to trypsin and PNGase A digestion, prior to labelling of the released N-glycans with 2-aminopyridine and off-line RP-HPLC/MALDI-TOF MS analysis; the MS/MS-verified structures are annotated. **(D)** Online LC-MS analysis of the tryptic peptides of the 46 kDa band indicate the highest score to a homologue of Poly-Cysteine and Histidine-Tailed Metalloproteins (PCHTP, otherwise known as p43; theoretical mass including signal peptide of 46.7 kDa) found in other Trichinellid species with an overall coverage of 83%; peptides with between 10 and 120 peptide spectrum matches are indicated in bold and potential N-glycosylation sites are in red.

## Discussion

Nematodes are amazing in terms of their glycomic diversity and, although it is a relatively well-explored species, *T. suis* is no exception. By combining glycan array and glycomic data, we have sought to close the gap between structure and immunological function of the N-glycans of this parasite. Here, we have significantly expanded the knowledge of the range of N-glycans in *T. suis*, showing the occurrence of tetra-antennary N-glycans with variations on the theme of LacdiNAc and have confidently defined some 200 N-glycan structures, including isomeric forms, as compared to the less than 40 previously described ^(*14*)^. This has been accomplished by using far more starting material, but also by fractionating by 2D-HPLC prior to MALDI-TOF MS/MS, whereby two labelling methodologies were employed: FMAPA for preparing glycans for printing, but not so suitable for isomeric glycan analysis and PA for the in-depth structural determination, but not compatible with printing. As many further masses over 3500 Da probably correspond to various isomers of highly substituted glycans, but MS/MS in the higher mass range did not result in full structural assignments, the actual number of N-glycan isomers will significantly exceed 200.

On the antennae of the higher molecular weight N-glycans, fucosylated versions of LacdiNAc as well as multiple phosphorylcholine modifications were observed (for a summary of glycan epitopes in *T. suis*, see ***Figure 10***). As previously found in *T. suis* and *D. immitis* ^(*14*)^, where there was a single modification of a non-fucosylated LacdiNAc, the phosphorylcholine tended to be on the terminal GalNAc rather than on the subterminal GlcNAc, which is in contrast to *C. elegans* ^(*37*)^; in the case of fucosylated LacdiNAc, on the other hand, the zwitterionic moiety was often on the GlcNAc, but HexNAc_2_Fuc_1_PC_2_ antennae as well as phosphorylcholine modification of the HexNAc substitution of fucose and of the underlying mannose are present. The largest detected FMAPA-glycan (***Supplementary Figures 1 and 4***; *m/z* 5059) would be predicted to contain nine zwitterionic moieties on a Hex_3_HexNAc_10_Fuc_5_ scaffold, but a higher degree of substitution cannot be ruled out. Certainly, our data indicate multiple positions for phosphorylcholination on *T. suis* tetra-antennary N-glycans (two on each of the maximally four antennae, in addition to one substitution of mannose), whereby the MS/MS fragmentation for the largest glycans is primarily limited to B-fragments. The theme of LacdiNAc, modified by phosphorylcholine or fucose, in the context of tetra-antennary glycans is shared with the related *Trichinella spiralis* ^(*38*)^; however, due to technical limitations in early studies, the degree of phosphorylcholinylation directly observed in *T. spiralis* was lower than that detected here in *T. suis*. Zwitterionic modification of mannose is rarer, but has been recently reported for one other nematode, *Brugia malayi*, and in filamentous fungi ^(*39,40*)^.

**Figure 10:**
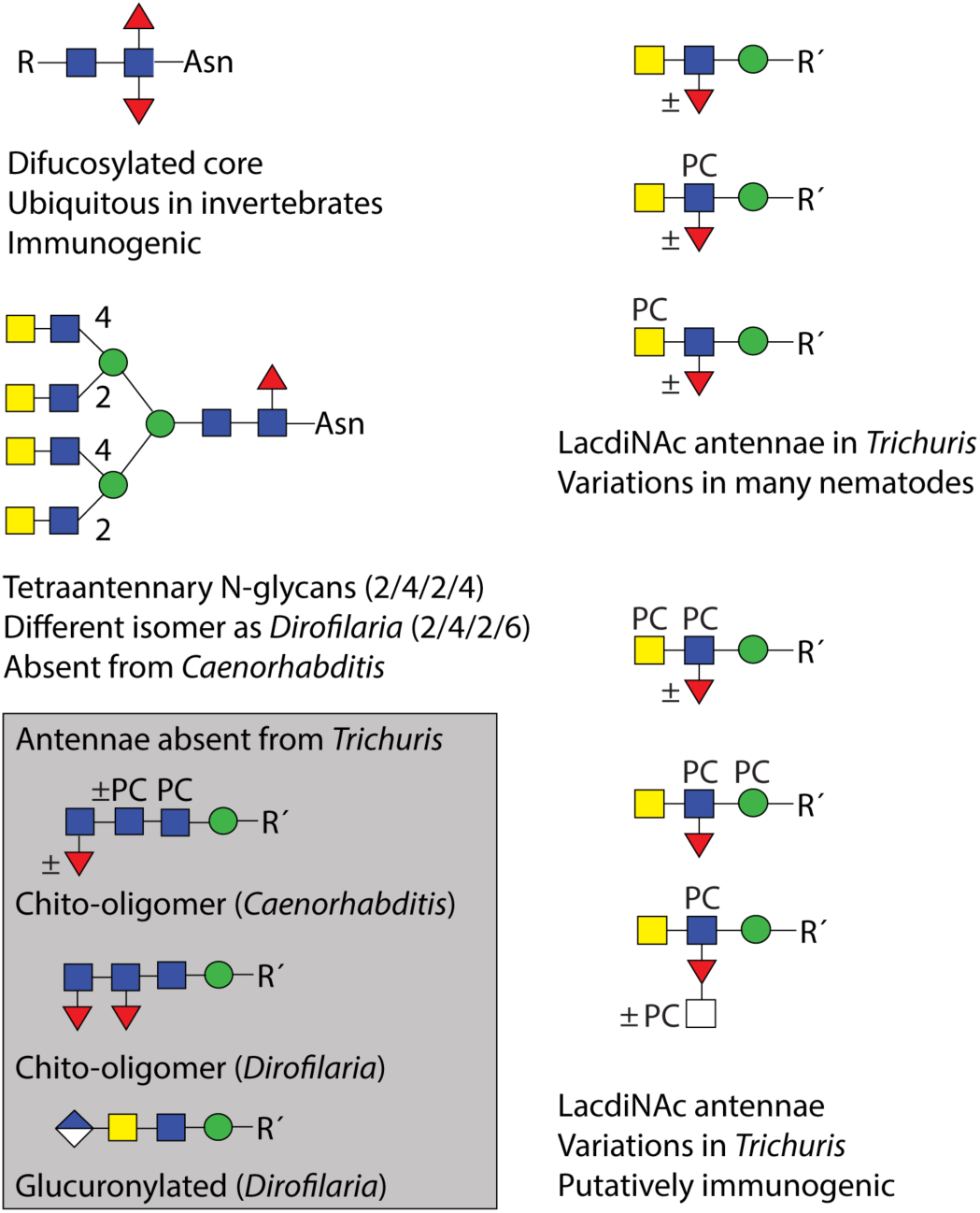
Summary of N-glycan motifs in *Trichuris* and selected other nematodes. Based on the N-glycomic data, the basic elements found or absent from *Trichuris suis* as compared to *Caenorhabditis elegans* and *Dirofilaria immitis* are shown. Glucuronylation, phosphorylcholine-modification of mannose and fucosylated LacdiNAc have also been reported in *Brugia malayi* or *Trichinella spiralis*. The presence of specific epitopes in other nematodes cannot be excluded and more variations may remain to be detected.

The one modification in *T. suis* that can be considered ‘species-specific’ is the modification of the fucose residue of the fucosylated LacdiNAc motif with a further HexNAc, often in combination with a phosphorylcholine modification. Although there are many instances of substituted fucose on N-glycans from nematodes ^(*31,37*)^ and molluscs with galactose or disubstitution with hexuronic acid and another monosaccharide ^(*41,42*)^, this is the only instance of an *N*-acetylhexosamine substitution of fucose in nematodes. On the other hand, unlike *C. elegans*, *O. dentatum*, *H. contortus* or filarial nematodes, there is no evidence for extended chito-oligomer chains ^(*23,24,31,43*)^, but akin to *D. immitis*, the N-glycan core region in *T. suis* is relatively unmodified (primarily α1,6-fucosylation and a trace of α1,3-fucosylation) and lacking the galactosylated fucose or the methylated epitopes found in *O. dentatum* or especially the highly modified cores in *C. elegans* ^(*23,37*)^. Also, other than some phosphorylated structures, no anionic N-glycans such as the glucuronylated ones found in *D. immitis* or *B. malayi* were detected ^(*15,39*)^. Due to the phosphate-modified glycans having the same ‘backbone’ structure as those carrying phosphorylcholine, we assume either that there is an artefact due to the initial sample preparation (reduction and alkylation prior to trypsin digestion) or that a cholinesterase has removed the choline moiety, leaving a phosphate residue; we have not observed such a remodelling in other species, but cholinesterases and phosphorylcholine hydrolases are known from other nematode species ^(*44*)^.

The glycan array is one of a few based on natural N-glycans from a nematode, others being focused on *C. elegans*, *D. immitis* and *B. malayi* ^(*15,39,45*)^. Beyond using the typical commercially-available plant and fungal lectins, we probed the arrays with antibodies recognizing glycan modifications and with proteins of the innate immune system. The typical lectins ConA, GNA and AAL recognised a variety of glycans as expected, whereas fewer fractions were bound by WGA and even fewer by WFA; the latter is a trend previously seen for binding to *T. suis* soluble products ^(*21*)^. Considering the abundance of antennal fucose, LTL bound fewer fractions than may be expected, but the high degree of modification of fucosylated LacdiNAc motifs in *T. suis* may interfere with binding. On the other hand, RCA predominantly bound a galactosylated control on the array and recognition by SNA was relatively low as compared to other standard lectins, compatible with a lack of sialic acid (***Figure 2*** and ***Supplementary Figure 5A***). Overall, the results with these lectins are in keeping with our own previous dot-blot based studies ^(*46*)^ as well as a recent large-scale reanalysis of glycan array data ^(*47*)^.

For the antibodies binding mono- and difucosylated LacdiNAc (LDNF and FLDNF) motifs, it was perhaps no surprise that the anti-LDNF antibody ^(*48*)^ bound a number of fractions containing LacdiNAc-modified glycans, including those with phosphorylcholine-modified forms of this epitope, but especially to fraction 14 containing a Hex_3_HexNAc_6_Fuc_2_ structure (***Supplementary Figures 3 and 5B***); the case of anti-FLDNF ^(*49*)^ is puzzling, as we detected no difucosylated LacdiNAc motif, but perhaps the multiple fucosylated antennae can also generate an epitope. Amongst the lower molecular weight N-glycans, the anti-mannose 100-4G11-A antibody ^(*50*)^ bound, as expected, fractions containing Man_3-5_GlcNAc_2_ and Man_3_GlcNAc_2_Fuc_1-2_. The TEPC-15 IgA antibody and the pentraxin C-reactive protein bound almost all of the fractions judged to contain phosphorylcholine-modified N-glycans in keeping with their known specificity ^(*51*)^.

Of the lectins of the innate immune system examined, DC-SIGN recognises a rather wide range of oligomannosidic and fucosylated ligands ^(*52–54*)^; in the present study, its binding correlated generally with the occurrence of Man_5-9_GlcNAc_2_ in the relevant fractions (***Figure 2***); this does not rule out binding to fucosylated LacdiNAc motifs as there was also some recognition of some of the largest molecular weight glycans, but it has been shown that glycans with a terminal GlcNAc on the α1,3-mannose with or without a LacdiNAc antenna on the α1,6-mannose are better recognized than those carrying LacdiNAc on the α1,3-mannose ^(*54*)^. Dectin-2 binding was lower on the array relative to other innate immune system lectins (***Figure 2***), despite reports that it binds larger oligomannosidic structures ^(*55*)^, although it bound well to the major 46 kDa glycoprotein (***Figure 9***) and in absolute terms recognized the larger molecular weight glycans (***Supplementary Figure 5B***). MGL, which is known to interact with LacdiNAc ^(*56*)^, bound the majority of fractions, regardless of the presence of phosphorylcholine-modifications of the putative LacdiNAc-containing ligands (***Figure 2***); the Western blotting results with MGL showing recognition of a 46 kDa band also correlated with the ‘band-specific’ glycomic data showing presence of LacdiNAc motifs (***Figure 9***). In terms of immune modulation by *T. suis* soluble products, it was concluded that their DC-SIGN-mediated effect on dendritic cells is indeed rather via the oligomannosidic glycans, whereas MGL may have a role in interactions of these soluble products with other cell types ^(*21*)^. Our data show an obvious increase in antibodies recognizing phosphorylcholine- and fucose-modified N-glycans of 2500-5000 Da during *T. suis* infection (***Figure 2*** and ***Supplementary Figure 5C***); binding to PC-modified glycans is in line with a number of studies on identified cestode and nematode antigens ^(*57–60*)^, although in some cases phosphorylcholine appears only to have a minor effect on IgG reactivity ^(*61*)^. Additionally, we observed binding to the widespread invertebrate core difucosylated motif, which is a known epitope for IgE in *Haemonchus*-infected sheep ^(*62*)^ and for anti-horseradish peroxidase ^(*63*)^. In terms of glycan array data for other helminths, the best studied example is the trematode *Schistosoma mansoni*, with which *T. suis* shares the LDNF and core α1,3-fucose motifs. Data from shotgun and defined arrays also show binding of IgG of *Schistosoma-*infected animals to various fucosylated antennal motifs ^(*49,64,65*)^, while an immune response to core α1,3-fucose ^(*66*)^ may be restricted to the IgG4 subclass ^(*65*)^.

The major protein band was shown to contain a homologue of Poly-Cysteine and Histidine-Tailed Metalloproteins (***Figure 9***), otherwise known as p43. This protein binds glycosaminoglycans and interleukin-13 ^(*35*)^, thus it represents a potential immunomodulator; in its native form, p43 is also a potent inducer of protective immunity, whereas the insect-derived recombinant form is not ^(*67*)^, but is a cryptic antigen in terms of natural infection ^(*35*)^. As noted above, the band-specific glycomic analysis indicates that p43 may well carry LacdiNAc-containing motifs with and without phosphorylcholine (***Figure 9C***), which is indeed a type of glycan modification found to a low level in insect cells ^(*13*)^; thus, reengineering of these cells to increase the proportion of nematode-like glycans would in theory generate p43 in a more native form and so increase the protective effect. Overall, the N-glycome of *T. suis* encompasses both simple and complex structures – with many modifications being variations of the LacdiNAc unit found in many invertebrates. Multiple antennal fucose (maximally four) and phosphorylcholine residues (at least up to seven) in addition to core fucosylation are possible; the combinations found are, to date, novel in *T. suis*, even if most of the modification types are represented in other species (***Figure 10***). This leads to the question as to which glycan-modifying enzymes are encoded by its genome. In the closely related *T. muris*, there are obvious homologues to three branching N-acetylglucosaminyltransferases (***Supplementary Figure 18***) indicating a different branching pattern as compared to *D. immitis*, as well as possibly ten α1,3-fucosyltransferases and one putative BRE-4-type LacdiNAc-forming enzyme; however, no nematode phosphorylcholinyltransferase has been identified to date and a fucose-modifying HexNAc-transferase would be novel. Such enzymes are of especial interest as they are responsible for biosynthesis of the glycans which are recognized by pentraxins or antibodies in the sera of infected animals, but which are also associated with immunomodulation; on the other hand, chemical syntheses of phosphodiester-modified glycans are far from routine. Thus, the availability of larger quantities of defined PC-glycan conjugates is limited until a chemoenzymatic approach becomes realisable. The combination of glycomics and glycan arrays presented here is a further step in the definition of complex nematode N-glycans and their interactions with host immune systems, while reinforcing the need for new tools to determine the role of these key post-translational modifications in the evolutionary success of this large group of often parasitic organisms.

## Experimental Procedures

### Biological material

Adult *Trichuris suis* worms were kindly provided by Dr. Irma van Die (Amsterdam University Medical Center). The parasites were isolated from pigs three months post infection; one part of the harvested parasites was maintained in media before further treatment. In total up to 29 g (wet weight) of worms were used for preparation. *T. suis* pooled uninfected and infected pig sera from different time points of infection (21 days, 28 days, 35 days and 49 days post infections) for glycan array analysis were kindly provided by Dr. Andrew Williams (University of Copenhagen).

### Enzymatic release and N-glycan purification for glycomic and array analyses

The worms were thoroughly washed with 0.98% NaCl solution using a 0.22 µm filter device (Millipore) prior to lyophilization and grinding in liquid nitrogen. The dry powder was dissolved in 20 ml lysis buffer (25 mM Tris, 150 mM NaCl, 5 mM EDTA and 0.5% (w/v) CHAPS, pH 7.4) followed by multiple rounds of sonification. The insoluble fraction was removed by centrifugation and the supernatant was dialysed prior to lyophilization. The dried glycoproteins were reduced using 1,4-dithiothreitol (Sigma; 0.18 M) followed by carboxymethylation using iodacetamide (Sigma; 0.18 M). The reduced and carboxymethylated *T. suis* protein homogenates were proteolysed using trypsin (Sigma-Aldrich, Trypsin Sequencing Grade from bovine pancreas, pH 8.4; 1 mg/ml), prior to a solid-phase extraction on 2 g C18. The glycopeptides were eluted from the C18 material with increasing 1-propanol concentration (20%, 40% and finally 100%). The eluates were combined and lyophilized overnight. Thereafter, the N-glycans were released using peptide:N-glycosidase F (NEB, in 50 mM ammonium bicarbonate, pH 8.4; twice 250 U/ml overnight) followed by peptide:N-glycosidase A (NEB, in 50 mM ammonium acetate, pH 4.0; 60 U/ml). The free glycans were separated from residual O-glycopeptides on a C18 solid-phase extraction. The glycans were eluted using 5% acetic acid and the residual peptides were released using increasing 1-propanol concentration (20%, 40% and finally 100%). The free N-glycans were analysed by MALDI-TOF MS (UltraFlex II MALDI/TOF Mass Spectrometer, Bruker equipped with a Smartbeam II laser).

### Immobilization of N-glycans

90% of the free N-glycans were conjugated to the fluorescent Fmoc-(3-(methoxyamino)propylamine) linker (abbreviated as FMAPA) as described by Wei et al ^(*22*)^. Briefly, the free N-glycans were incubated with 0.35 M Fmoc-MAPA linker, 0.5 M sodium acetate and 0.1 M 2-amino-5-methoxybenzoic acid (2-AM) at 65°C for 4h. Afterwards, a ten-fold excess of ethylacetate was added and the mixture incubated at −20°C for at least 20 minutes to precipitate the neoglycoconjugates. The pellet was washed 3 times prior to solid phase extraction on C18 (Waters) material. The successful labelling of the *T. suis* N-glycans was determined by MALDI-TOF MS. The Fmoc part was released by incubating the labelled N-glycans with 5% -methyl-piperidine for 30 minutes at room temperature under constant shaking. The chemical was removed by chloroform precipitation prior C18-SPE purification. The MAPA-labelled glycans were analysed by MALDI-TOF MS prior to printing on NHS-slides. The Fmoc-MAPA labelled N-glycans were fractionated by a semi-preparative normal phase HPLC (Luna, Phenomenex, LC Column, 250 × 10 mm, 5 µm, 100 Å) on a Shimadzu HPLC CBM-20A system, equipped with a UV detector (SPD-20AV) and a fluorescence detector (RF-20A). The following gradient was applied using a three-buffer system (Buffer A - 250 mM ammonium acetate, pH 4; buffer B – dH_2_O HPLC grade water; buffer C – 99% (v/v) acetonitrile) at a flow rate of 4 ml/min: 0-5 min, 16% B and 4% C; 5-50 min, 16-40% B and 4-50% C; 60 – 60.1 min back to starting conditions and hold for another 10 min. Selected peaks were re-injected on a second dimension using a reversed phase analytical C18 – HPLC column (Agilent Zorbax Eclipse Plus 95Å C18, 150 x 2.1 mm, 5 µm, analytical). The following gradient of buffer D (water) and buffer E (0.1% (v/v) TFA in acetonitrile) was applied at a flow rate of 1 ml/min: 0-4 min, 5-50% E; 40-45 min, hold 50% E; 45-46 min, 50-60% E; 46-51 min, hold 60% E; 51-52 min, back to starting conditions and hold for another 8 min. The detector was set to Ex/Em 265/315 nm. The column was calibrated daily in terms of in house prepared standard Lacto-*N*-neotetraose (LnNT) FMAPA glycans. The MAPA-glycans were diluted to 100 µM in spotting buffer (100 mM Sodium phosphate, pH 8.5) and printed using a sciFLEXARRAYER S11 (Scienion, Germany) on NHS activated glass slides (SCHOTT Nexterion, Slide H) by non-contact piezo printing. For each probe, 4 spots were printed per array. After 16 h of hybridization, slides were blocked (50 mM ethanolamine in 100 mM sodium borate, pH 9.0) and subsequently rinsed with TSM-WB (1xTSM, 0.05% Tween-20) and water. The slides were sealed and stored dry at −20°C until use. The slides were rehydrated in TSM buffer (20 mM Tris-Cl, 150 mM sodium chloride, 2 mM calcium chloride and 2 mM magnesium chloride) before incubation with (i) 10 µg/ml biotinylated lectins ConA, AAL, LTL, GNA, WGA, WFA, RCA and SNA (VectorLabs in 1xTSM, 0.05% Tween-20 and 1% BSA, *i.e.* TSMBB) followed by 2 µg/ml anti-biotin antibody AF 488 conjugated (Invitrogen) in TSMBB; (ii) 2 µg/ml anti-LDNF (L6B8 in TSMBB)^(*48*)^ and anti-FLDNF (F2D2 in TSMBB)^(*49*)^ followed by anti-mouse IgG AF 647 conjugated (1:1000, Invitrogen) in TSMBB or 5 µg/ml anti-mannose antibody (100-4G11-A) ^(*50*)^ followed by 2 µg/ml anti-mouse IgM AF 647 (Invitrogen); (iii) 5 µg/ml human CRP (biotechne) in TSMBB with additional 2 mM Ca^2+^ followed serially by anti-CRP from mouse (biotechne) in TSMBB and 2 µg/ml anti-mouse IgG AF 647 in TSMBB; (iv) 10 µg/ml TEPC-15 (Sigma-Aldrich) followed by 2 µg/ml anti-mouse IgA FITC in TSMBB; (v) 5 µg/ml human DC-SIGN-Fc (R&D System), 5 µg/ml human MGL-Fc (R&D System) and 1 µg/ml human Dectin-2-Fc (Sino Biological) in TSMBB followed by 5 µg/ml anti-mouse IgG AF 488 (Invitrogen); (vi) 1:250 dilution of uninfected and infected pig sera (provided by Dr. Andrew Williams, University of Copenhagen) followed by anti-pig IgG or IgM antibody (1:500 diluted in TSMBB). Binding was detected by 2 µg/ml anti-mouse AF 647 (Invitrogen). After each incubation step, slides were serially washed with TSM-WB, TSM and water. The dried slides were scanned with a GenePix 4300A Scanner (Molecular Devices) equipped with multiples lasers with a laser power from 10-100% and PMT gain from 450-600, and the images processed using the GenePix Pro 7.2 software (Molecular Devices). The raw fluorescent values were automatically normalized and the background was automatically substracted from each individual spot. The generated data files (gpr file) were further analyzed by Excel (Microsoft). The average relative fluorescence units (RFU) were calculated as averaged of all fluorescence values of each probe (i.e., 4 spots) and the standard deviations were calculated and plotted. Probes with generally less than 500 RFU in the assays and/or poor MS data were excluded from the presented data; heat maps were generated using the GLAD tool ^(*68*)^.

### In-depth N-glycan analysis

The remaining 10% of the free N-glycans (see above) was subject to solid-phase extraction on non-porous graphitised carbon (SupelClean ENVICarb, Sigma-Aldrich) and eluted with 40% acetonitrile. N-glycans were labelled via reductive amination using 2-aminopyridine (PA). For 1D-HPLC, 10% of the pyridylaminated N-glycome was fractionated by HIAX-HPLC (IonPac AS11 column, Dionex, 4 × 250 mm, combined with a 4 × 50 mm guard column) was used with a two-solvent gradient (buffer A, 0.8 M ammonium acetate (pH 3.85), and buffer B, 80% acetonitrile, LC-MS grade) at a flow rate of 1 ml/min as folllows: 0–5 min, 99% B; 5–50 min, 90% B; 50–65 min, 80% B; 65–85 min, 75% B. The HIAX-HPLC was calibrated using oligomannosidic PA-labelled bean glycans. All manually collected HPLC glycan fractions were analyzed after lyophilization by MALDI-TOF MS and MS/MS. Selected fractions were combined and subject to a 2D-HPLC analysis by reversed-phase HPLC (Ascentis Express RP-amide from Sigma-Aldrich; 150 × 4.6 mm, 2.7 µm) and a gradient of 30% (v/v) methanol (buffer B) in 100 mM ammonium acetate, pH 4 (buffer A), was applied at a flow rate of 0.8 ml/min (Shimadzu LC-30 AD pumps) as follows: 0–4 min, 0% B; 4–14 min, 0–5% B; 14–24 min, 5–15% B; 24–34 min, 15–35% B; 34–35 min, return to starting conditions. The RP-amide HPLC column was calibrated daily in terms of glucose units using a pyridylaminated dextran hydrolysate and the degree of polymerisation of single standards was verified by MALDI-TOF MS. Monoisotopic MALDI-TOF MS was performed using an Autoflex Speed (Bruker Daltonics) instrument in either positive or negative reflection mode with 6-aza-2-thiothymine or 2,5-dihydroxybenzoic acid as matrix. In general, MS/MS was performed by laser-induced dissociation of [M+H]^+^ or [M-H]^−^ ions; typically 2000 shots were summed for MS (reflector voltage, lens voltage, and gain of 27 kV, 9 kV, and 2059 V, respectively) and 4000 for MS/MS (reflector voltage, lift voltage, and gain of 27 kV, 19 kV, and 2246 V, respectively). Spectra were processed with the manufacturer’s software (Bruker Flexanalysis 3.3.80) using the SNAP algorithm with a signal/noise threshold of 6 for MS (unsmoothed) and 3 for MS/MS (smoothed four times). Glycan spectra were manually interpreted on the basis of the masses of the predicted component monosaccharides; differences of mass in glycan series, comparison with coeluting structures from other insects, marine organisms or other nematodes, and fragmentation patterns before and after chemical treatment or exoglycosidase digestion. A list of theoretical *m/z* values for each glycan composition is presented in Supplementary Tables 1 and 2.

### Enzymatic and chemical treatments

Glycans were treated, prior to re-analysis by MALDI-TOF–MS, with α-fucosidase (bovine kidney from Sigma-Aldrich or almond α1,3/4-specific from Prozyme), α-mannosidase (jack bean from Sigma), or β-*N*-acetylhexosaminidases (jack bean from Sigma-Aldrich, *Streptomyces plicatus* chitinase from New England Biolabs or in-house-produced recombinant *Caenorhabditis elegans* HEX-4 specific for β1,4-GalNAc-linked residues) in 50 mM ammonium acetate, pH 5, at 37 °C overnight (except for pH 6.5 in the case of HEX-4). Hydrofluoric acid was used for removal of core or antennal α1,3-fucose, phosphorylcholine or phosphate. As appropriate, treated glycans were re-chromatographed by RP-amide HPLC to ascertain retention time shifts prior to MALDI-TOF-MS.

### O-linked glycan analysis by β-elimination

55 mg sodium borohydride (NaBH_4_, Sigma-Aldrich) were dissolved in 1 ml 0.1 M NaOH solution. 400 µl were added to a fraction of the completely dried residual *T. suis* glycopeptides after PNGase F/A release and incubated over night at 45°C. The reaction was stopped by adding pure acetic acid. The mixture was initially purified by a cation exchange material (Dowex AG50 H^+^ form, Bio Rad). The free O-glycans were released by 5% acetic acid, lyophilized prior to co-evaporation using 10% acetic acid in methanol. This step was repeated at least twice to remove the remaining salts. Afterwards, the sample was further cleaned using a C18-SPE. Briefly, the sample was resuspended in 50% methanol and loaded on the column. The O-glycans were released using 5% acetic acid and further permethylated prior to analysis by MALDI-TOF MS.

### Glycan permethylation

For permethylation, *T. suis* N- and O-glycans were dried in a glass tube and resuspended in 1 ml of a slurry of grinded NaOH pellets (Sigma-Aldrich) in DMSO followed by adding 500 µl iodomethane. The mixture was incubated for 20 min at room temperature under constant shaking and then the reaction was quenched using 200 µl dH_2_O. Subsequently, permethylated N- and O-glycans were extracted in chloroform by constant washing with dH_2_O and then applied to a pre-equilibrated C18 solid-phase extraction column. Salts and contaminants were removed using 15% (v/v) acetonitrile prior to elution of the permethylated glycans using a 50% (v/v) acetonitrile solution. MALDI-TOF MS was performed using an Autoflex Speed or a Rapiflex (Bruker Daltonics) instrument in positive reflection mode with 2,5-dihydroxybenzoic acid as matrix.

### Western blotting

15 µg worm proteins were denatured at 95°C in 5 x SDS loading buffer (5% β-mercaptoethanol, 0.02% bromophenol blue, 30% glycerol, 10% SDS and 250 mM Tris-Cl pH 6.8) for 10 minutes prior to SDS-PAGE separation (Bio-Rad, 10% acrylamide, 150 V) for subsequent analysis by either Coomassie staining or blotting on nitrocellulose. For western blot analysis the nitrocellulose blot was blocked in 3% bovine serum albumin in 0.5% in PBST or 0.5% BSA in TBST (blocking buffer) at 4°C. Afterwards, the membrane was incubated with either 10 µg/ml biotinylated lectins (Vector Laboratories), Dectin-2, DC-SIGN or hMGL (R&D Systems) or 1:200 dilution of murine IgA TEPC 15 (Sigma-Aldrich), 1:200 dilution of human C-reactive protein (MPBIO) including additional 5 mM CaCl_2_ or 1:1000 anti-HRP from rabbit (Sigma) in blocking buffer. After washing with PBST (TBST), the Dectin-2, DC-SIGN or hMGL were incubated an anti-human IgG Alexa Fluor 690 (Jackson ImmunoResearch) and after extensive washing in PBST the dried membrane was scanned with an imaging system (GE Healthcare Amersham Typhoon). For lectins, the binding was detected by 10 µg/ml streptavidin coupled with alkaline phosphatase; the TEPC-15 binding was detected by an anti-mouse IgA conjugated with alkaline phosphatase. For CRP, the membrane was first incubated with a 1:1000 dilution of a murine anti-CRP antibody (biotechne) prior to detection with a 1:2000 dilution coupled anti-mouse IgG coupled with alkaline phosphatase. The binding of the alkaline phosphatase-conjugated antibodies was monitored with SigmaFAST BCIP/NBT (Sigma).

### Proteomic analysis

Selected bands were cut from the SDS-PAGE gel (∼15 µg/lane). After washing and destaining proteins were fixed in the gel and reduced with dithiothreitol and alkylated with iodoacetamide. In-gel digestion was performed with trypsin (Trypsin Gold, Mass Spectrometry Grade, Promega, Madison, WI; Trypsin-ultra^TM^, MS grade NEB) with a final trypsin concentration of 20 ng/µl in 50 mM aqueous ammonium bicarbonate and 5 mM CaCl_2_. Digest proceeded for 8 hours or overnight at 37°C. Afterwards, peptides were extracted thrice with 50 µL of 5% trifluoroacetic acid in 50% aqueous acetonitrile supported by ultrasonication for 10 min. Extracted peptides were dried down in a vacuum concentrator (Eppendorf, Hamburg, Germany) and resuspended in 0.1% TFA for LC-MS/MS analysis. Peptides were separated on a nano-HPLC Ultimate 3000 RSLC system (Dionex). Sample pre-concentration and desalting was accomplished with a 5 mm Acclaim PepMap μ-Precolumn (300 µm inner diameter, 5 µm, 100 Å; Dionex). For sample loading and desalting 2% acetonitrile in ultra-pure H_2_O with 0.05% TFA was used as a mobile phase with a flow rate of 5 µl/min. Separation of peptides was performed on a 25 cm Acclaim PepMap C18 column (75 µm inner diameter, 2 µm, 100 Å) with a flow rate of 300 nl/min. The gradient started with 4% B (80% acetonitrile with 0.08% formic acid) for 7 min, increased to 31% in 30 min and to 44% in additional 5 min. It was followed by a washing step with 95% B. Mobile Phase A consisted of ultra-pure H_2_O with 0.1% formic acid. For mass spectrometric analysis the LC was directly coupled to a high-resolution Q Exactive HF Orbitrap mass spectrometer. MS full scans were performed in the ultrahigh-field Orbitrap mass analyzer in ranges *m/z* 350−2000 with a resolution of 60 000, the maximum injection time (MIT) was 50 ms and the automatic gain control (AGC) was set to 3e^6. The top 10 intense ions were subjected to Orbitrap for further fragmentation via high energy collision dissociation (HCD) activation over a mass range between *m/z* 200 and 2000 at a resolution of 15 000 with the intensity threshold at 4e^3. Ions with charge state +1, +7, +8 and >+8 were excluded. Normalized collision energy (NCE) was set at 28. For each scan, the AGC was set at 5e^4 and the MIT was 50 ms. Dynamic exclusion of precursor ion masses over a time window of 30s was used to suppress repeated peak fragmentation.

### Protein glycan analysis

For protein-specific glycan analysis, the glycopeptides were heat treated to inactivate the trypsin and the sample was subject to treatment with PNGase F (250U, Sigma; pH 8) followed by PNGase A (NEB; pH 6). The released N-glycans were purified using initially a cation exchange material (Dowex AG50 H^+^ form, Bio Rad) and the glycans were eluted using 2% acetic acid, whereas the residual peptides were eluted using 0.1 M ammonium acetate, pH 6.0. N-glycans were purified using a column packed respectively nonporous graphitized carbon/Lichroprep C18. The column was pre-washed with 100% acetonitrile and equilibrated with water. The N-glycans were then eluted with 40% acetonitrile containing 0.1% trifluoroacetic acid. After drying, the glycans were fluorescently labeled by reductive amination using 2-aminopyridine and then analyzed with MALDI-TOF MS and RP-HPLC as described above.

### Phylogenetic analyses

Defined N-acetylglucosaminyltransferase protein sequences from *Homo sapiens* as annotated by NCBI (MGAT1, P26572; MGAT2, Q10469; MGAT3, Q09327; MGAT4A, Q9UM21; MGAT4B, Q9UQ53; MGAT4C, Q9UBM8; MGAT5A, Q09328) and *Caenorhabditis elegans* as annotated in Wormbase (MGAT1, GLY-12, GLY-13 and GLY-14; MGAT2, GLY-20; MGAT5, GLY-2) were used as a query to deep searching homologous sequence of the whole nematode proteome (NCBI 11.01.2023) using hidden Markov Models algorithm from phmmer (http://hmmer.org/). All found sequences were used to built alignment with MAFFT ^(*69*)^ and after the final approximately-maximum-likelihood phylogenetic tree was build with FastTree tool ^(*70*)^. The resulting phylogeny tree was visualized with iTOL ^(*71*)^.

## Acknowledgements

This work was funded in part by the Austrian Science Fund (FWF) by grants to K.P. [P 32572], I.B.H.W. [P 33453] and J.V. [P 35516]. B.E. and Z.D. were funded by the FWF-funded BioTOP doctoral programme [W 1224]. C.G., A.Y.M. and R.D.C. were funded through the National Center for Functional Glycomics (NIH grants P41GM103694 and R24GM137763 to R.D.C.). The authors thank Dr. Irma van Die for the *Trichuris* worms, Dr. Andrew Williams for the porcine sera and Angelika Paschinger for preparation of Supplementary Figure 5. This research was supported using resources of the VetCore Facility (Proteomics) of the Veterinärmedizinische Universität Wien and the Core Facility Mass Spectrometry of the Universität für Bodenkultur.

## Data Availability

The glycomics data is available via Glycopost ^(*72*)^ (GPST000360). The mass spectrometry proteomics data have been deposited to the ProteomeXchange Consortium via the PRIDE partner repository with the dataset identifier PXD045512 and 10.6019/PXD045512.

## Supplemental Data

The Supplement contains further information on the glycomics and glycan array experiments as well as Supplementary Figures 1-18 and Supplementary Tables 1 and 2.

## Author contributions

B.E. performed glycomics and glycan array experiments and prepared some figures and tables; C.G., A.Y.M. and R.D.C. contributed novel reagents, glycan array printing and supervised the glycan array experiments; Z.D. performed bioinformatic analyses; J.V. performed and analysed permethylation analyses; K.P. supervised the glycomics experiments, interpreted data and prepared figures; I.B.H.W wrote the text, prepared figures and performed some mass spectrometry experiments. All authors contributed to reviewing the manuscript.

