## Supplementary Information for "Recognition of highly branched N-glycans of the porcine whipworm by the immune system"

### Further information regarding the glycomic analyses

#### *Definition of the level of the glycan structural analysis:*

The goal was the N-glycomic analysis of *Trichuris suis* adults. Thereby, individual glycan-containing HPLC fractions of glycans labelled with PA [2-aminopyridine] were subject to MALDI-TOF MS and MS/MS, in combination with a range of chemical and enzymatic treatments. In addition, glycans labelled with FMAPA [Fmoc-(3-(methoxyamino)propylamine)] were also fractionated by HPLC, analysed by MALDI-TOF MS and immobilized for use in glycan array experiments.

#### *Search parameters and acceptance criteria*

a. **Peak lists:** As stated in the methods section: typically 2000 shots were summed for MS and 4000 for MS/MS. Spectra were processed with the manufacturer's software (Bruker Flexanalysis 3.3.80) using the SNAP algorithm with a signal/noise threshold of 6 for MS (unsmoothed) and 3 for MS/MS (four-times smoothed).
b. **Search engine, database and fixed modifications:** All glycan data were manually interpreted and no search engine or database was employed; for the pyridylaminylated glycans, the fixed modification is the pyridylamine label at the reducing end ($GlcNAc_1$-PA fragments of *m/z* 300), whereas the FMAPA-labelled glycans generally show an $^{O,2}A$ fragment, resulting from loss of 409 Da from the parent upon MS/MS.
c. **Exclusion of known contaminants and threshold:** All glycan data were manually interpreted; only peaks with an MS/MS consistent with a pyridylaminated chitobiose core were included – the 'threshold' for inclusion was an interpretable MS/MS spectrum (at least in terms of composition).
d. **Enzyme specificity:** A description of the release methods (PNGase F followed by PNGase A) is given in the methods section. Enzymes used during the analysis (glycosyl hydrolases) are defined in the methods by species name and supplier. Citations for in-house purified recombinant enzymes are also given. As previous experience with normalizing glycosidase amounts based on units of activity towards *p*-nitrophenyl sugars reduced digestion efficiency towards native oligosaccharides, aliquots of glycans (equivalent to 5 – 50 mV in terms of fluorescence) were incubated with 0.2 µl of the various enzyme preparations (whether commercial, desalted commercial or in-house produced) overnight. These conditions result in no obvious unspecific removal of residues as defined by shifts in mass, MS/MS or retention times, although steric hindrance in some glycans leads to a requirement for longer incubation times (48 hours). Ammonium acetate buffers were used (supplemented with $CaCl_2$ where required) as suppliers' buffers interfere with MALDI-TOF MS analysis; generally one-quarter of any glycosidase digest was applied directly to the target plate prior to drying and addition of 6-aza-2-thiothymine as matrix. Hydroflouric acid treatment (3 µl of 48% HF added to the dried glycan) was 24 or 48 hours on ice in the cold room prior to drying under vacuum; expected release of α1,3-fucose and phosphodiester residues, but not of other sugars, was observed under these conditions. Incubating the N-glycans with β-galactosidases or α-*N*-acetylhexosaminidases did not affect any mass spectra.

   ***Fucosidases:*** bovine α-fucosidase (Sigma, native) removes core α1,6-fucose, but not core α1,3-fucose; almond α1,3/4-fucosidase (Prozyme, native) removes antennal α1,3-fucose from the subterminal GlcNAc of LacdiNAc, but apparently neither defucosylates chitobiose antennae nor a terminal GalNAc of a LacdiNAc motif and proved inefficient with larger structures; thus, hydrofluoric acid was more effective for the fucose removal in this study. α1,2-Fucosidases were not employed in this study.

***Hexosaminidases:*** Jack bean β-*N*-acetylhexosaminidase (Sigma, native) is a general-purpose enzyme unspecifically removing β-linked GlcNAc and GalNAc residues, but as enzyme lots age they lose the ability to remove GalNAc from substituted LacdiNAc motifs; *Streptomyces* β1,3/4-*N*-acetylhexosaminidase (NEB, 'chitinase' recombinantly-expressed in *E. coli*) removes β1,4-linked GalNAc from LacdiNAc and β1,4-linked GlcNAc from chitobiose motifs; *C. elegans* HEX-4 is an in-house prepared enzyme (His-tag purified recombinant form expressed in *Pichia*) which demonstrably removes β1,4-linked GalNAc residues, but in this and previous studies is not observed to remove any other HexNAc; insect FDL is also prepared in-house (His-tag purified recombinant form expressed in *Pichia*) and under the stated conditions (3 hour incubation) only removes the β1,2-linked GlcNAc attached directly to α1,3-mannose in Man3-based hybrid and biantennary glycans, in keeping with its proven role in the *in vivo* generation of paucimannosidic glycans.

***Mannosidases:*** Jack bean α-mannosidase (Sigma or Prozyme, native) removes all α-mannose residues, but steric hindrance slows its action, e.g., digestion of core α1,6-mannose is reduced if there is a 'lower' arm modification on the core α1,3-mannose; more specific α1,2-, α1,2/3- or α1,6-mannosidases were not used in this study.

e. **Isobaric/isomeric assignments:** For isobaric/isomeric species, 2D-HPLC elution, differences in MS/MS and/or digestion data were used for the assignment (as described in the text).

#### *Glycan or glycoconjugate identification*

a. **Precursor charge and mass/charge (*m/z*):** All glycans detected were singly-charged. For the positive mode, the *m/z* values for pyridylaminated glycans or FMAPA-labelled phosphorylcholine-modified glycans are for protonated forms, whereas in negative mode the ions are $[M\text{-}H]^-$ and most FMAPA-labelled glycans and all permethylated glycans are detected in the sodiated form. Depending on the glycan amount or presence of buffers in exoglycosidase preparations, the relative amounts of the $H^+$ and $Na^+$ adducts varied. Maximally two decimal places used for the *m/z* annotations consistent with the accuracy of MALDI-TOF MS; in the figures and due to space limitations, only one decimal place is presented. Previous data indicate an average +0.03 Da (+ 22 ppm) deviation between the measured and the calculated *m/z* values on the instrument used.

b. **MALDI-TOF MS settings (positive and negative modes):** Ion Source 1 and 2 were 19.00 and 16.75 kV; Lens, 9.00 kV (7.95 kV for over 3000 Da or for negative mode); Reflector 1 and 2, 21.05 and 9.65 kV; Pulsed Ion Extraction, 160 ns (120 ns in negative mode); Matrix Suppression typically up to 700 Da; Detector Gain, typically 2059 V.

c. **MALDI-TOF MS/MS settings (positive and negative modes):** Ion Source 1 and 2 were 6.00 and 5.35 kV; Lens, 2.90 kV; Reflector 1 and 2, 27.00 and 11.75 kV; Lift 1 and 2, 19.00 and 4.00 kV; Pulsed Ion Extraction, 140 ns; Detector Gain, typically 2246 V when fragmenting; Laser Power Boost typically 50%; not in CID mode; PCIS typically 0.65%.

d. **All assignments:** For the glycans present in each pool, see the HIAX and RP-amide-HPLC chromatograms annotated with structures shown according to the Standard Nomenclature for Glycans. Downwardly- and upwardly-drawn core fucose and mannose residues are respectively α1,3- and α1,6-linked (see **Figure 3**, inset, in the main text).

e. **Modifications observed:** Listed are the *m/z* values for glycans carrying a reducing terminal pyridylamine group as judged by presence of an *m/z* 300 $GlcNAc_1$-PA fragment. As the glycans are otherwise chemically unmodified, $\Delta m/z$ of 80, 146, 162, 165 and 203 correspond to phosphate, deoxyhexose (presumed to be fucose), hexose, phosphorylcholine or *N*-acetylhexosamine. No natural methylation and no glucuronic acid residues were observed.

In the case of permethylated samples, phosphorylcholine residues were removed due to the experimental procedure.

f. **Number of assigned masses:** Glycan assignments were not just based on measured mass only, but on at least MS/MS, in most cases corroborated by digest and elution data. The linear mass spectra of higher molecular weight glycans are shown to indicate their presence but not their structure.
g. **Spectra:** Representative annotated spectra (MS and MS/MS) defining structural elements are given in various figures. In total, MS and/or MS/MS data for over 75 of the approximately 200 structures are shown in the figures, representative of all glycostructural motifs; nearly 200 mzxml files have also been uploaded to Glycopost. The overall data is based on some 2500 MS and 3500 MS MS/MS spectra.
h. **Structural assignments:** As noted in the results section, the typical oligomannosidic structures are assigned based on elution time and fragmentation pattern; it is otherwise assumed that the glycans contain a trimannosyl core consistent with typical eukaryotic N-glycan biosynthesis. The presence of an GlcNAcTIV (MGAT4) homologues in *T. suis* is compatible with the proposed tri- and tetra-antennary glycans, whose fragmentation patterns show preferential loss of the 'heaviest' antenna.
The assignments of antennal and core fucose residues are based on RP-HPLC retention time, fragmentation pattern and/or susceptibility to digestions. Other antennal modifications (phosphorylcholine and *N*-acetylgalactosamine; including anomericity of the glycosidic linkage) are defined based on digestions and fragmentation patterns with rechromatography after digestion in some cases. Phosphorylcholine is assumed to be 6-linked via phosphodiesters to HexNAc or Man residues; antennal fucose and *N*-acetylgalactosamine are assumed to be α1,3-linked and β1,4-linked respectively, while core fucose is α1,6-linked, except in the case of core α1,3/α1,6-difucosylation. There is no evidence of in-source fragmentation of either neutral terminal monosaccharides (including Lewis-type fucosylation) or phosphorylcholine (as evidenced by this and previous publications).

***Summary of glycosidases used in this study***

| | |
|---|---|
| PNGases | *Flavobacterium* PNGase F (recombinant, *E. coli*)<br>*Oryza sativa* PNGase A (recombinant, *Pichia*) |
| Fucosidases | Bovine α-fucosidase (native)<br>Almond α1,3/4-fucosidase (native) |
| Hexosaminidases | Jack bean β-*N*-acetylhexosaminidase (native)<br>*Streptomyces* β1,3/4-*N*-acetylhexosaminidase/chitinase (recombinant, *E. coli*)<br>*Caenorhabditis* HEX-4 β1,2-*N*-acetylgalactosaminidase (recombinant, *Pichia*)<br>Honeybee FDL β1,2-*N*-acetylglucosaminidase (recombinant, *Pichia*) |
| Mannosidases | Jack bean α-mannosidase (native) |

### Further information regarding the glycan array analyses

| **1. Glycan Binding Samples** | |
|---|---|
| Description of Sample | *Fungal/plant lectins:* Concanavalin A (ConA), wheat germ agglutinin (WGA); *Aleuria aurantia* Lectin (AAL); *Ricinus communis* agglutinin 1 (RCA-I); *Sambucus nigra* agglutinin (SNA); *Lotus tetragonolobus* lectin (LTL); *Galanthus nivalis* agglutinin (GNA); *Wisteria floribunda* agglutinin (WFA); (Vector Laboratories); used at 10 µg/ml.<br>*Pathogen-associated molecular pattern receptors:* Macrophage galactose lectin (hMGL, R&D Systems); DC-SIGN (R&D Systems); Dectin-2 (Sino Biological); human C-reactive protein (biotechne); used at 1 or 5 µg/ml.<br>*Anti-glycan antibodies:* TEPC-15 (Sigma, M1421); diluted 1:200, anti-LDNF (clone F6B8); diluted 1:900, anti-FLDNF (anti-F2D2); diluted 1:50, anti-mannose AB, used at 1:400.<br>Control serum was from the pigs prior to infection; the sera of pigs infected with *T. suis* were collected at time points 21, 28, 35 and 49 days post T. suis infection; sera were diluted 1:250. |
| Assay protocol | The slides were incubated (all dilutions in TSMBB) with either:<br>(i) Biotinylated forms of recombinant or commercial lectins (10 µg/ml) followed by anti-biotin antibody AF488 conjugated (Invitrogen, 2 µg/ml).<br>(ii) DC-SIGN-Fc (5 µg/ml), Dectin-2-Fc (1 µg/ml) and hMGL2-Fc (5 µg/ml) followed by anti-mouse IgG AF 488 (Invitrogen, 2 µg/ml).<br>(iii) Human C-reactive protein (biotechne, 5µg/ml) followed by anti-CRP IgG from mouse (biotechne; diluted 1:1000 and finally anti-mouse IgG, AF 647 (Invitrogen, 2 µg/ml))<br>(iv) TEPC-15 (Sigma, C1740; 1:200, i.e., 10 µg/ml) followed by anti-mouse IgA FITC (Invitrogen, 2 µg/ml).<br>(v) Anti-LDNF antibody (clone F6B8, diluted 1:900), anti-FLDNF antibody (clone F2D2, diluted 1:50) and anti-mannose antibody (100-4G11-A) followed by anti-mouse IgG or IgM AF 647 (Invitrogen, 2 µg/ml).<br>(vi) Pig sera (1: 250) followed by mouse anti-pig IgM or anti-pig IgG (Sigma, 1:500) and finally anti-mouse IgG AF647 conjugate (Invitrogen,2 µg/ml). |
| Incubation and washing | Each incubation step was one hour; washing was by dipping ten times in TSMWB, then ten times in TSM and then water, followed by drying. |
| Sample modifications | Fluorescent antibodies (conjugated with either AlexaFluor-647/488 or FITC) were used which detect the relevant primary antibodies or which were specific for biotin in order to detect binding of biotinylated lectins. |
| **2. Glycan Library** | |
| Glycan description for defined glycans | Refer to Figure 2 for SNFG-style structures of the positive controls and selected defined natural glycans. |
| Glycan description for undefined glycans | N-glycans were prepared by PNGaseF/A release of glycopeptides derived by tryptic proteolysis of *T. suis* homogenates; see the experimental procedures for further details regarding isolation, FMAPA-labelling and HPLC as well as Supplementary Figures 1-4 for MALDI-TOF and HPLC profiles. A detailed analysis of pyridylaminated *T. suis* N-glycans is also presented. Normalisation was on the basis of fluorescence intensity of HPLC peaks. |

| | |
|---|---|
| Glycan modifications | Glycan pools were derivatised with FMAPA (closed ring) and the pools were HPLC purified to remove residual linker prior to re-pooling; the glycans were treated with piperidine prior to printing. |
| **4. Arrayer (Printer)** | |
| Description of Arrayer | Scienion Flexarrayer S11 |
| Dispensing mechanism | Non-contact printing with the supplier's nozzles (Type 3) and a pulse of 45 µs, ca. 103 V. |
| Glycan deposition | Four replicates of 0.8 nl each, except for fractions. For FMAPA-labelled natural glycan pools, an estimated 2 nmol per spot was printed as estimated by fluorescence area of HPLC peaks in comparison to a standard. |
| Printing conditions | Derivatised glycans and oligosaccharides were mixed 1:1 with spotting buffer (100 mM sodium phosphate pH 8.5); printing was at room temperature; arrays were left to hybridise overnight prior to blocking (50 mM ethanolamine in 100 mM sodium borate, pH 9.0) for 1 h at RT, washing (in TSM + Tween, followed by TSM alone, and finally $H_2O$) and drying. |
| **5. Glycan Microarray** | |
| Array layout | 8 subarrays per slide with 4 replicates per glycan sample. |
| Glycan identification and quality control | Glycans on the array were either pools of natural glycans or chemically/chemoenzymatically synthesized oligosaccharides. All glycans were HPLC purified and verified by MALDI-TOF MS/MS.<br>Positive controls for lectin binding were included. Spotting buffer alone was used as one of the negative controls. |
| **6. Detector and Data Processing** | |
| Scanning hardware | GenePix 4300A Scanner |
| Scanner settings | Multiple photomultiplier tube (PMT) gain values from 450-600 and laser power from 10-100% |
| Image analysis software | GenePix Pro 7 |
| Data processing | Raw .tif image files were imported to GenePix 7.2.22; feature diameter was set to 100 µm and fluorescent intensities of spots in each sub-array were analysed. Resulting data (F525 or F635 mean values) were exported into Excel raw fluorescent values were automatically normalized and the background was automatically removed from each individual spot without subtraction; significance calculations and t-tests (parametric, unpaired, two-tailed, confidence level 95%) were performed in Excel. |
| **7. Glycan Microarray Data Presentation and Interpretation** | |
| Data presentation | Heat maps summarizing the data are shown in Figure 2. Bar charts with error bars are shown in Supplementary Figure 5. No software used for interpretation |

**Supplementary Figure 1. MALDI-TOF MS and HPLC fractionation of FMAPA-labelled *T. suis* N-glycans.** N-glycans were labelled with FMAPA and analysed by MALDI-TOF MS in positive mode prior to normal phase HPLC using an NH2 Luna column. Fractions collected were again analysed by MALDI-TOF MS and pooled or refractionated on an RP-HPLC column (see ***Supplementary Figures 2-4*** for MS and example MS/MS spectra). Elution regions for the fractions **1-27** corresponding to those shown for the array data in ***Figure 2*** are indicated in addition to glycan categories. Compositions of pauci- and oligomannosidic glycans are written in green (detected as $[M+Na]^+$), simple fucosylated glycans in black (detected as $[M+Na]^+$) and PC-modified glycans in blue (detected as $[M+H]^+$). For a list of *m/z* for FMAPA-labelled glycans refer to ***Supplementary Table 1***; *m/z* differences of 146, 162, 165 and 203 correspond to Fuc, Hex, PC or HexNAc. Of the original 39 fractions, twelve with generally low lectin/antibody binding and/or significant impurities as judged by MS were excluded from the presented data.

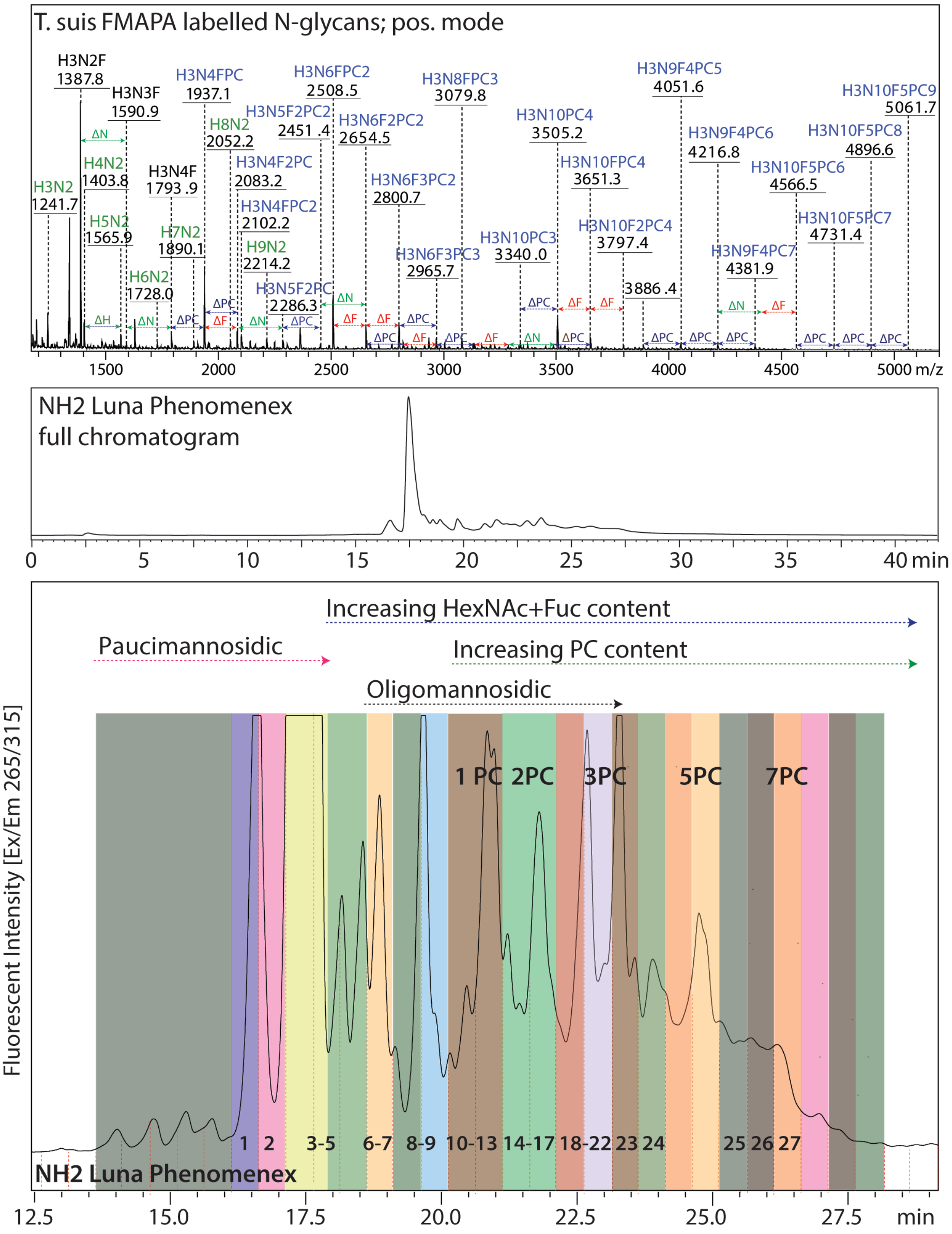

**Supplementary Figure 2. MALDI-TOF MS of FMAPA-labelled *T. suis* N-glycans (fractions 1-9).** NP-HPLC fractions (up to 20 min elution time; see ***Figure 1*** and ***Supplementary Figure 1***), partly refractionated by RP-HPLC, used in the glycan array experiments (***Figure 2***) were analysed by MALDI-TOF MS and MS/MS. For fractions 10-27 refer to ***Supplementary Figures 3 and 4***. While MS/MS of $[M+Na]^+$ parent ions yielded intense $^{0,2}$A fragments (loss of 409 Da), fragmentation of $[M+H]^+$ parent ions was biased to B fragments. Major reactivities for each fraction are summarized. Glycans are depicted according to the Symbol Nomenclature for Glycans. Theoretical *m/z* values are given in ***Supplementary Table 1***.

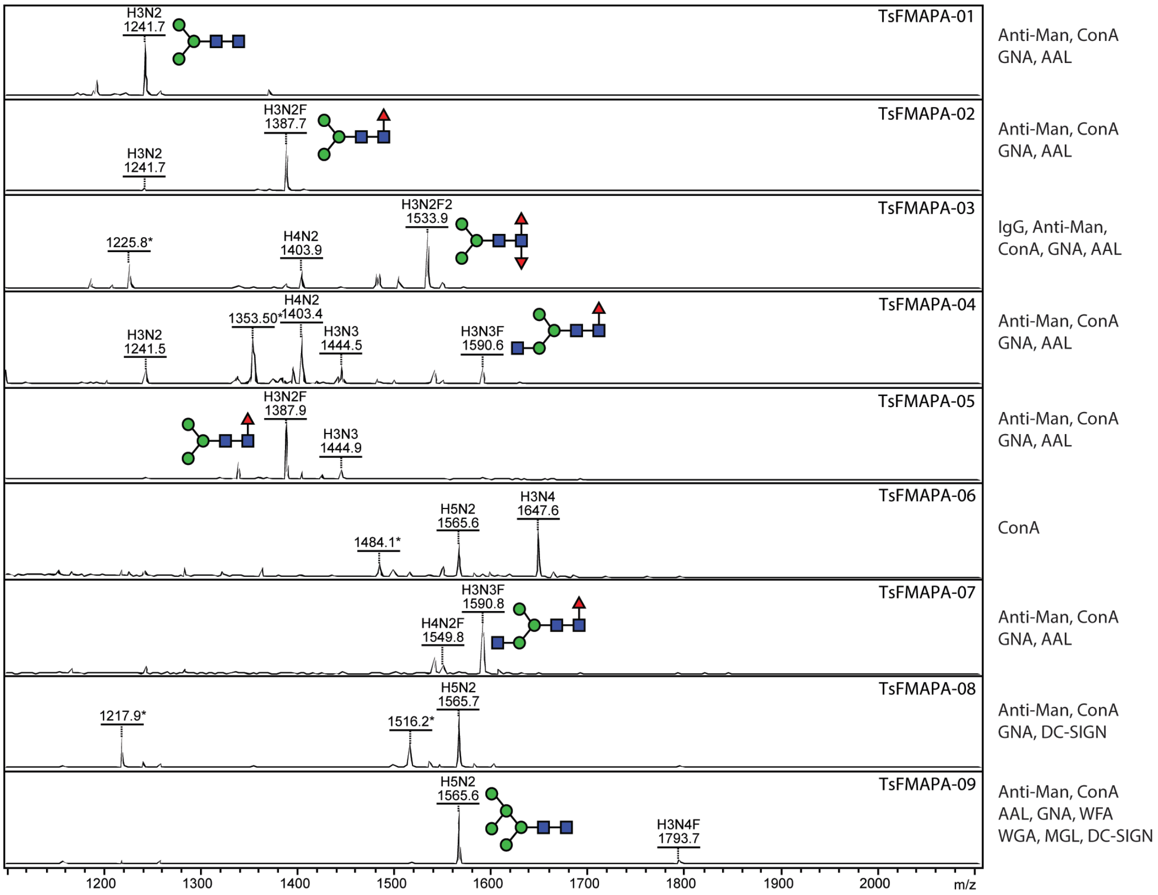


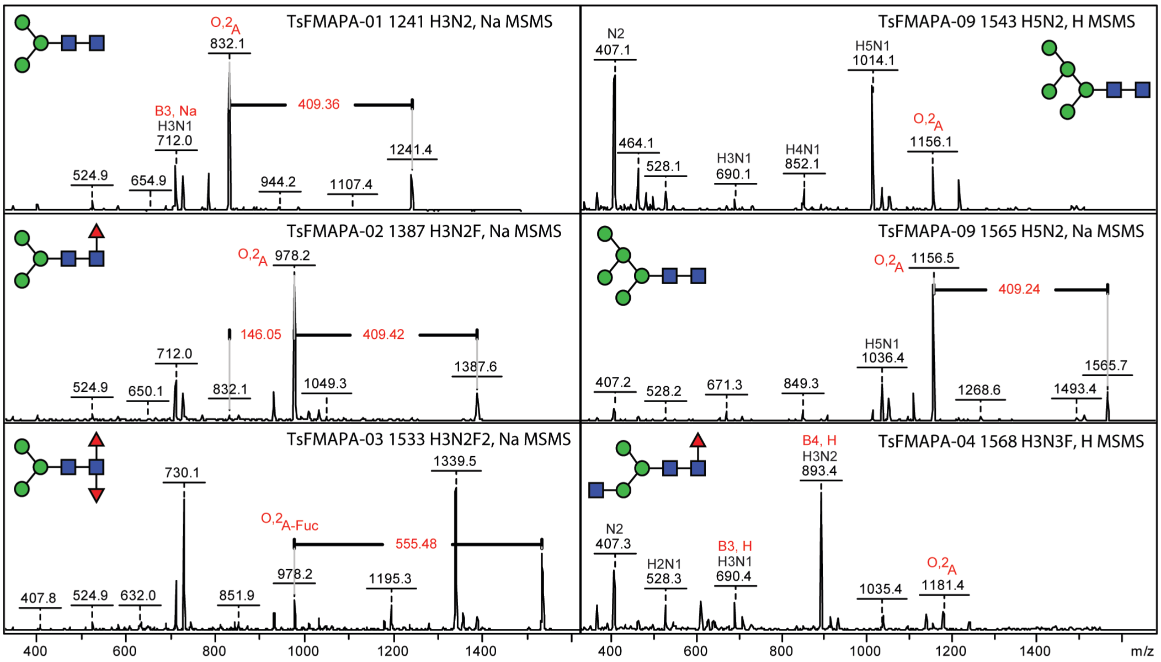

**Supplementary Figure 3. MALDI-TOF MS of FMAPA-labelled *T. suis* N-glycans (fractions 10-18).** NP-HPLC fractions (20-22 min elution time, see ***Figure 1*** and ***Supplementary Figure 1***, refractionated by RP-HPLC) were enriched in $Man_{6-7}GlcNAc_2$ and glycans modified with 1-2 phosphorylcholine residues. Interpretation of MS/MS was aided by comparison to the corresponding PA-labelled glycans, whereby the intense PC-containing B fragment ions are informative regarding the antennal motifs (see ***Figures 5-7*** in the main text as well as ***Supplementary Figures 10-17***).

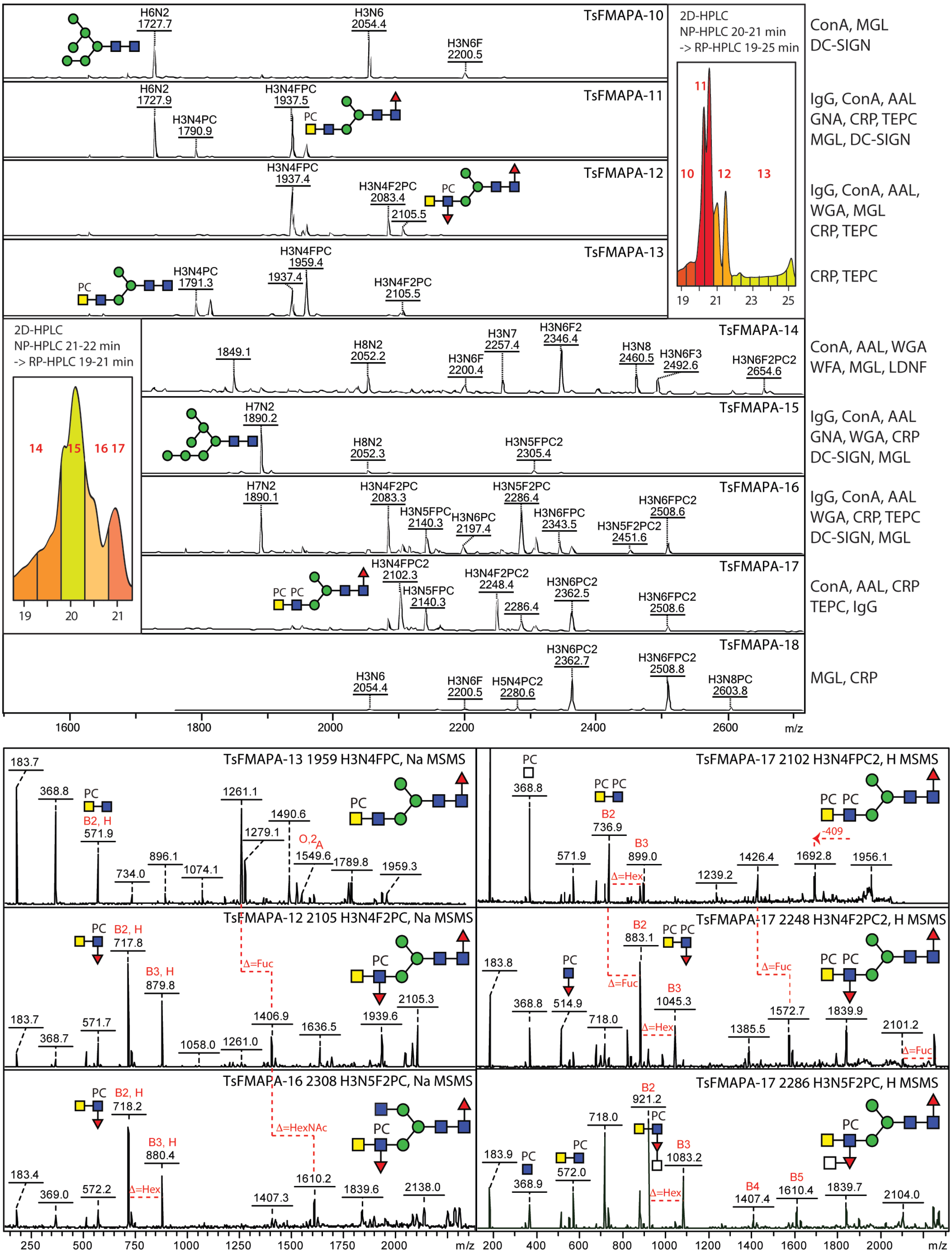

**Supplementary Figure 4. MALDI-TOF MS of FMAPA-labelled *T. suis* N-glycans (fractions 19-27).** NP-HPLC fractions (up to 22-26 min elution time, partly refractionated by RP-HPLC) were enriched in $Man_{8-9}GlcNAc_2$, $Glc_1Man_9GlcNAc_2$ and glycans modified with two or more phosphorylcholine residues. Interpretation of MS/MS was aided by comparison to similar PA-labelled glycans (see, e.g., ***Figures 5 and 7***), whereby the B fragment ions and the neutral losses from parent mass are informative regarding the antennal motifs. The isomeric annotations are suggested, not definitive. The calibration of the largest glycans was performed with ACTH peptides (*m/z* 2093, 2932, 3658 and 4539).

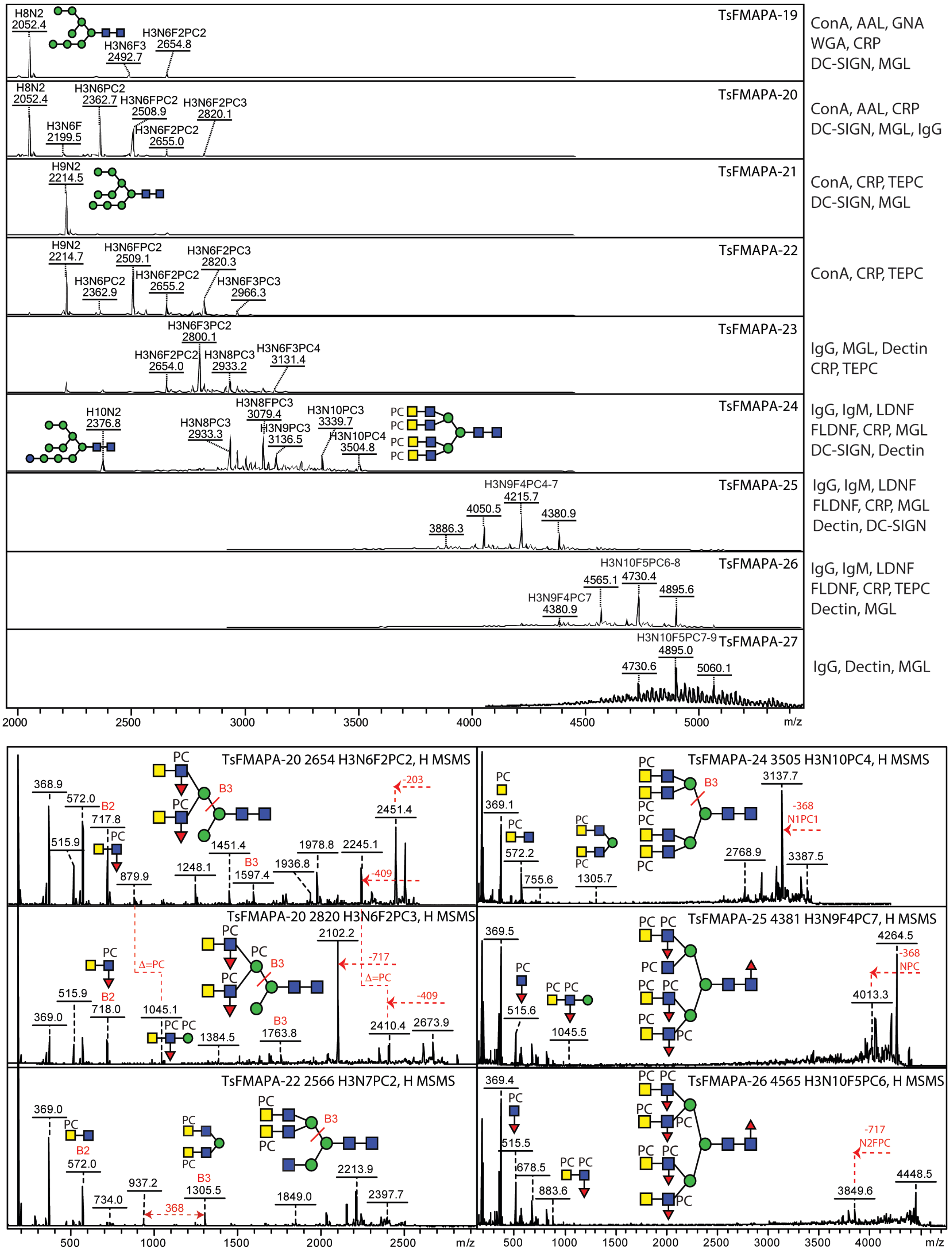

**Supplementary Figure 5A.** Raw glycan array data for biotinylated plant and fungal lectins; for the heat map summary and MALDI-TOF MS for the individual glycan fractions 1-27, refer to ***Figure 2*** and ***Supplementary Figures 2-4***. Standard glycans 28-32 and the buffer blank 33 are controls. Note that the maximal relative fluorescence units for SNA are 2000 as compared to 30000-70000 for the other standard lectins, reflecting a lack of sialylated glycans on the array, whereas RCA binds well to the lacto-N-tetraose control 28, but relatively poorly to any *T. suis* glycans. For details regarding the procedures, refer to the "further information regarding the glycan array analyses" section. A table with the individual data points is available as a supplementary Excel file.

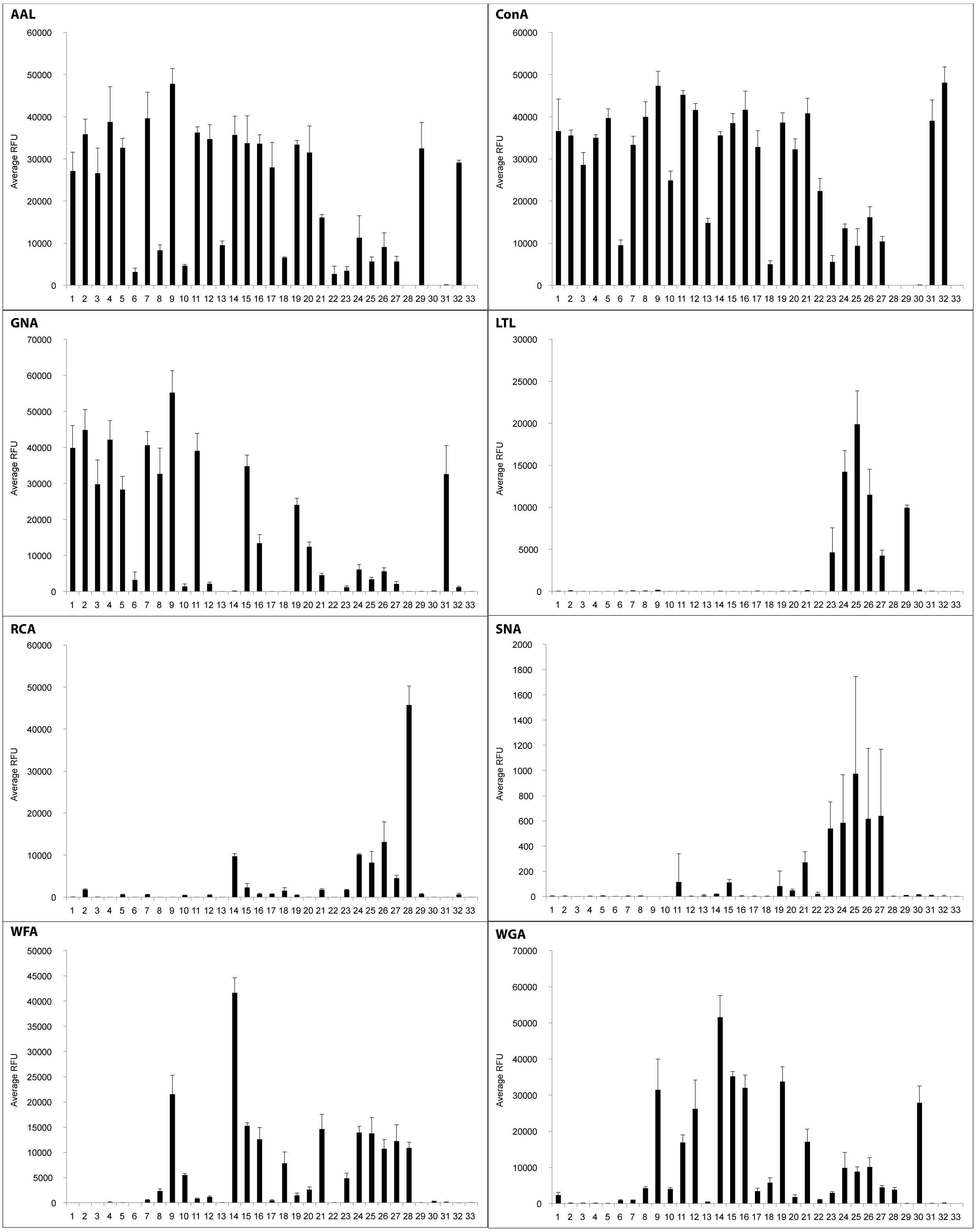

**Supplementary Figure 5B.** Raw glycan array data for monoclonal antibodies and mammalian innate immunity lectins. Monoclonal antibodies anti-FLDNF (F2D2), anti-LDNF (L6B8), anti-Man (100-4G11-A) and TEPC-15 (anti-phosphorylcholine) as well as mammalian lectins DC-SIGN, Dectin-2 and hMGL2 and the pentraxin C-reactive protein were employed. For the heat map summary, refer to ***Figure 2***, whereby the data for anti-FLDNF and anti-LDNF are adjusted 10- or 5-fold in the heat map comparison with anti-Man, due to the low absolute relative fluorescent units (RFU).

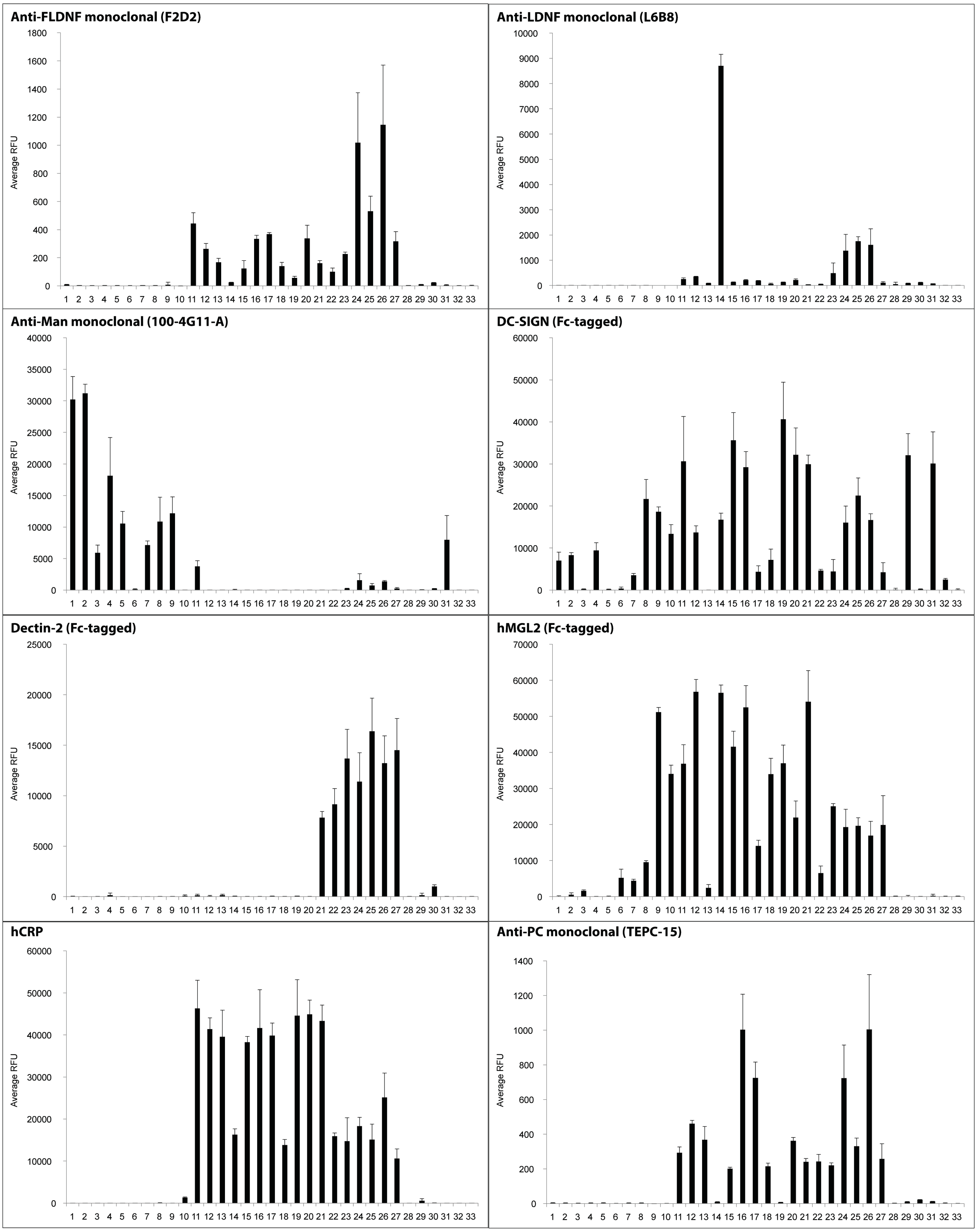

**Supplementary Figure 5C.** Raw glycan array data for pooled sera of uninfected and infected pigs. For the heat map summary, refer to ***Figure 2***. In absolute RFU, the highest binding of IgG and IgM is for 21-28 days post-infection and is predominantly towards the tetra-antennary N-glycans in fractions 24-26, but IgG reactivity towards a glycan with a difucosylated core (fraction 3) is observed for 28-49 days. The largest molecular weight glycans in fractions 24-26 are recognized by IgG and IgM; the IgG signal for the difucosylated glycan in fraction 3 is ca. 1000 RFU for days 28-49.

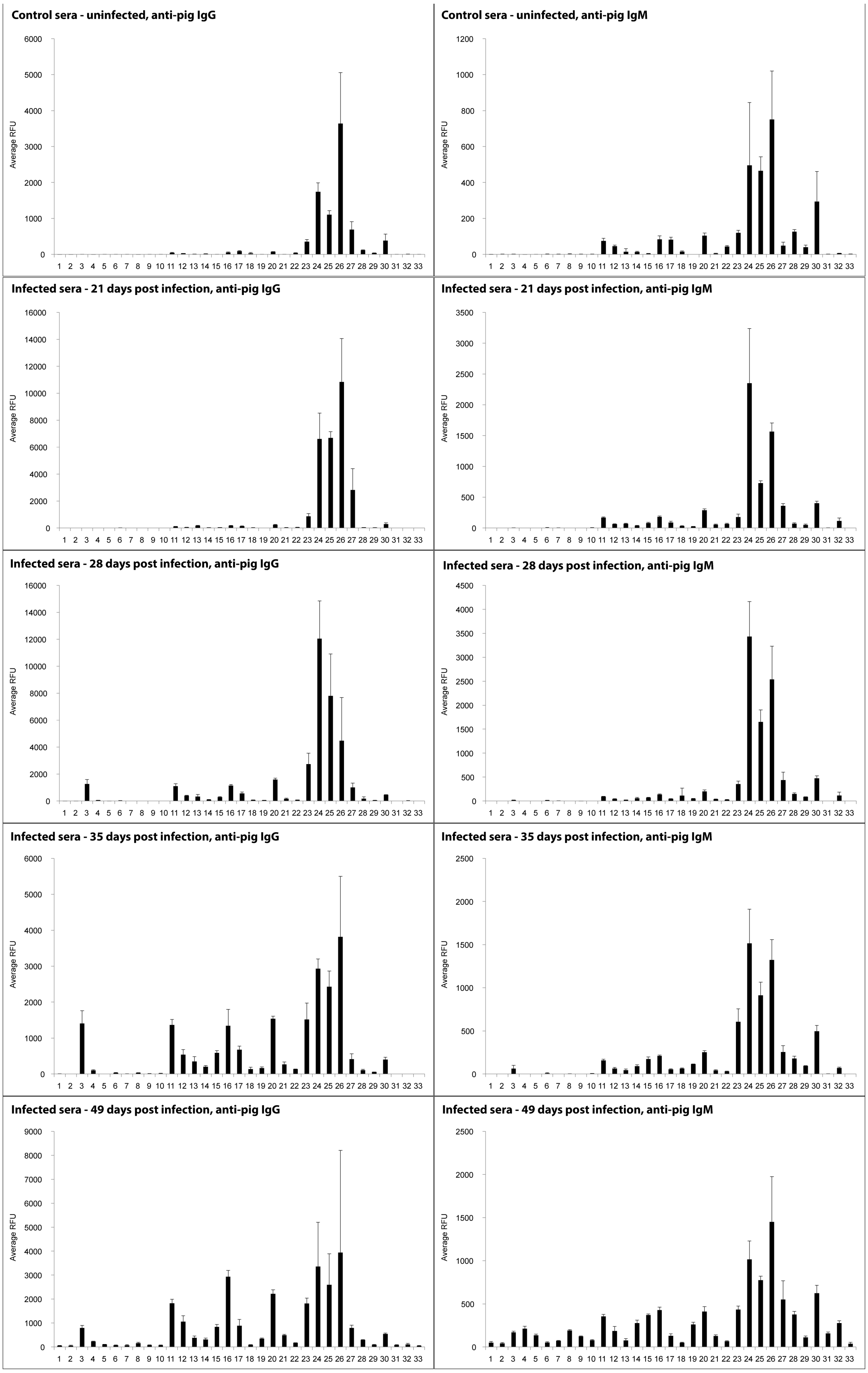

**Supplementary Figure 6. Permethylation of *T. suis* N-glycans. (A and B)** PNGase F and A-released N-glycans analysed by MALDI-TOF MS before and after permethylation. **(C)** MS/MS of a free phosphorylcholine-modified N-glycan. **(D-G)** MS/MS of four permethylated N-glycans with fucosylated LacdiNAc motifs, including a HexNAc-substituted fucose as also observed with the PA-labelled N-glycans (see ***Figure 6*** and ***Supplementary Figures 14-16***). Note that phosphorylcholine is alkali labile and is not observed after permethylation. The annotated *Δm/z* of 190, 259 and 419 are interpreted as corresponding to disubstituted Hex, terminal HexNAc and a HexNAcFuc motif, while low *m/z* fragments at m/z 474, 701 and 946 are interpreted to respectively correspond to reducing terminal GlcNAcFuc, the fucosylated LacdiNAc and the HexNAc-substituted fucosylated LacdiNAc.

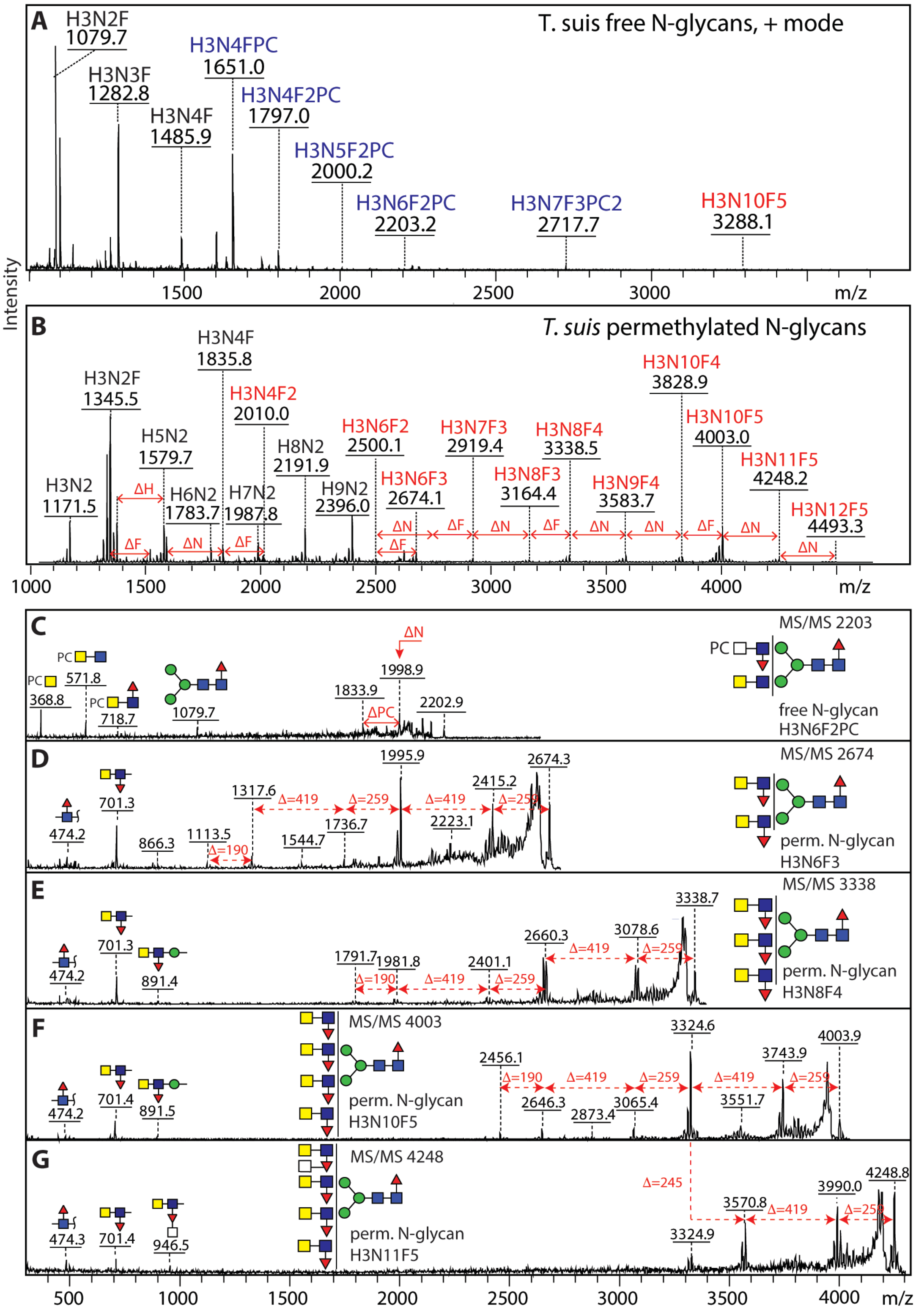

**Supplementary Figure 7. Size-fractionation of *T. suis* N-glycans.** HIAX (hydrophilic interaction/anion exchange; AS11 column) HPLC was performed twice on the pool of PNGase F/A-released N-glycans. The order of elution is approximately according to size, but oligomannosidic structures elute rather late as compared to other glycans of similar mass, while HexNAc-substitution of antennal fucose reduces retention and phosphorylcholine or phosphate modifications increase retention. Selected fractions were pooled on the basis of MALDI-TOF MS data and rechromatographed on an RP-amide column (see ***Figure 3*** of the main text for 2D-HPLC of fractions containing N-glycans of 2000-3500 Da, ***Supplementary Figure 8*** for 2D-HPLC of N-glycans of 1100-2000 Da and ***Supplementary Figure 11*** for 2D-HPLC N-glycans of around 3500 Da). Theoretical *m/z* values, proposed structures and fraction numbers, as well as Glytoucan accessions for glycans up to 2000 Da, are listed in ***Supplementary Tables 1 and 2***. Fraction numbers are shown in red as are masses of core difucosylated glycans. The elution positions for a set of oligomannosidic standards are also indicated (Man5, Man6, Man7, Man8 and Man9).

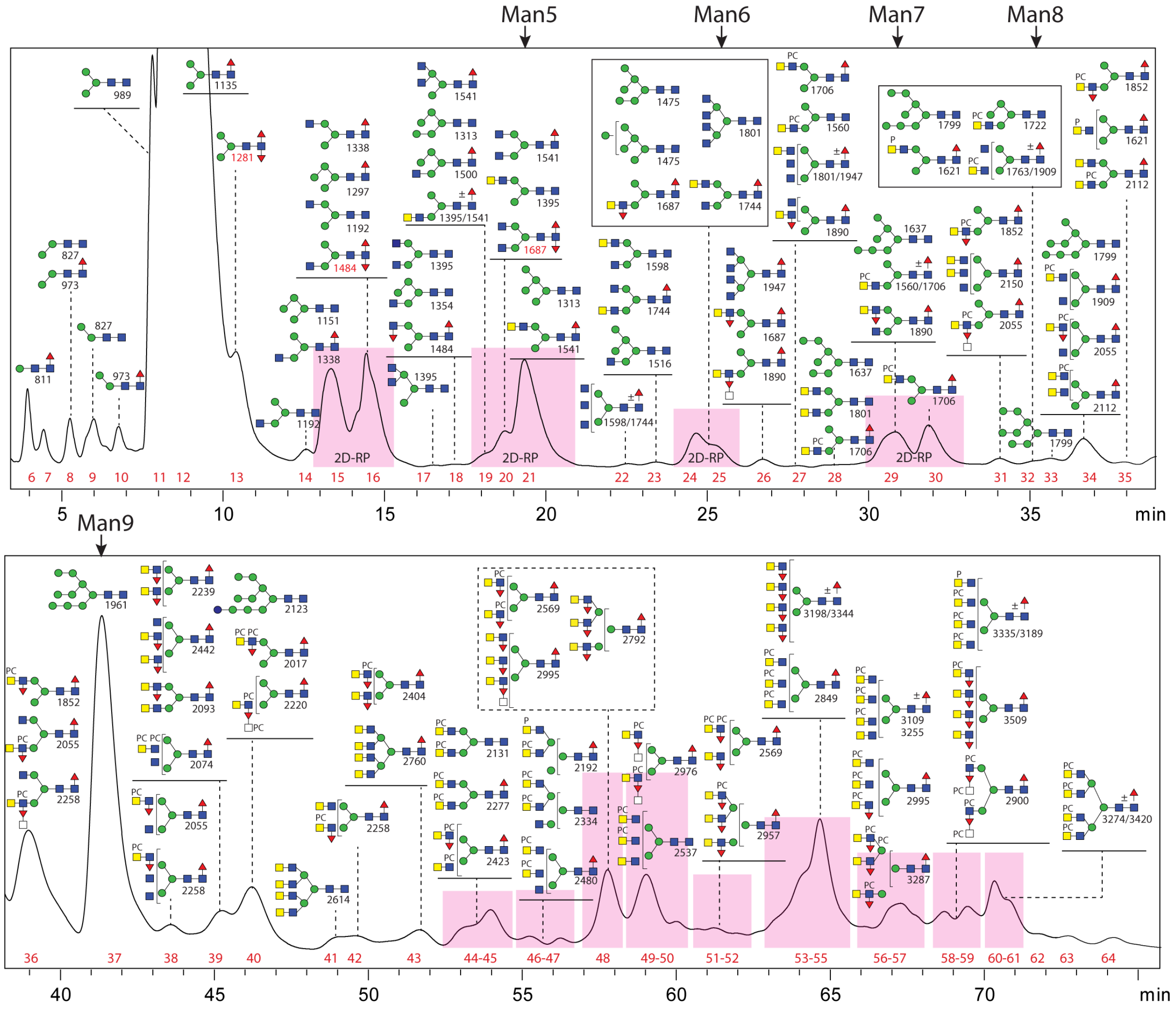

**Supplementary Figure 8. 2D-HPLC of lower molecular weight *T. suis* N-glycans.** Four HIAX glycan pools containing glycans of between 1100 and 2000 Da were subject to a second dimension of HPLC on an RP-amide column (fractions 9-20 from a second HIAX run, corresponding to fractions 14-30 from the first HIAX as shown in ***Supplementary Figure 7***); for the simple structures, annotations are based on MS/MS and comparisons to previous elution times for the same column (e.g., the two isomers of *m/z* 1338 eluting at 8.5 and 11.5 g.u. elute similarly to those in *Dirofilaria*; Martini *et al.*, 2019, *Nat. Commun*. 10, 75). However, some bi- and tri-antennary and all tetra-antennary structures elute late on RP-amide (18-20 g.u.), suggesting a different isomeric structure as compared to *Dirofilaria*. The 11.5 g.u. *m/z* 1744 structure is also found in insects; defucosylation of these insect glycans co-elute with a desialylated/degalactosylated glycan derived from bovine fetuin, indicative of the lower β1,4-antenna (Kurz *et al.*, 2015, *J. Proteomics*, 126, 172). See ***Figure 4*** and ***Supplementary Figures 9*** for example MS and MS/MS data on glycans in these fractions.

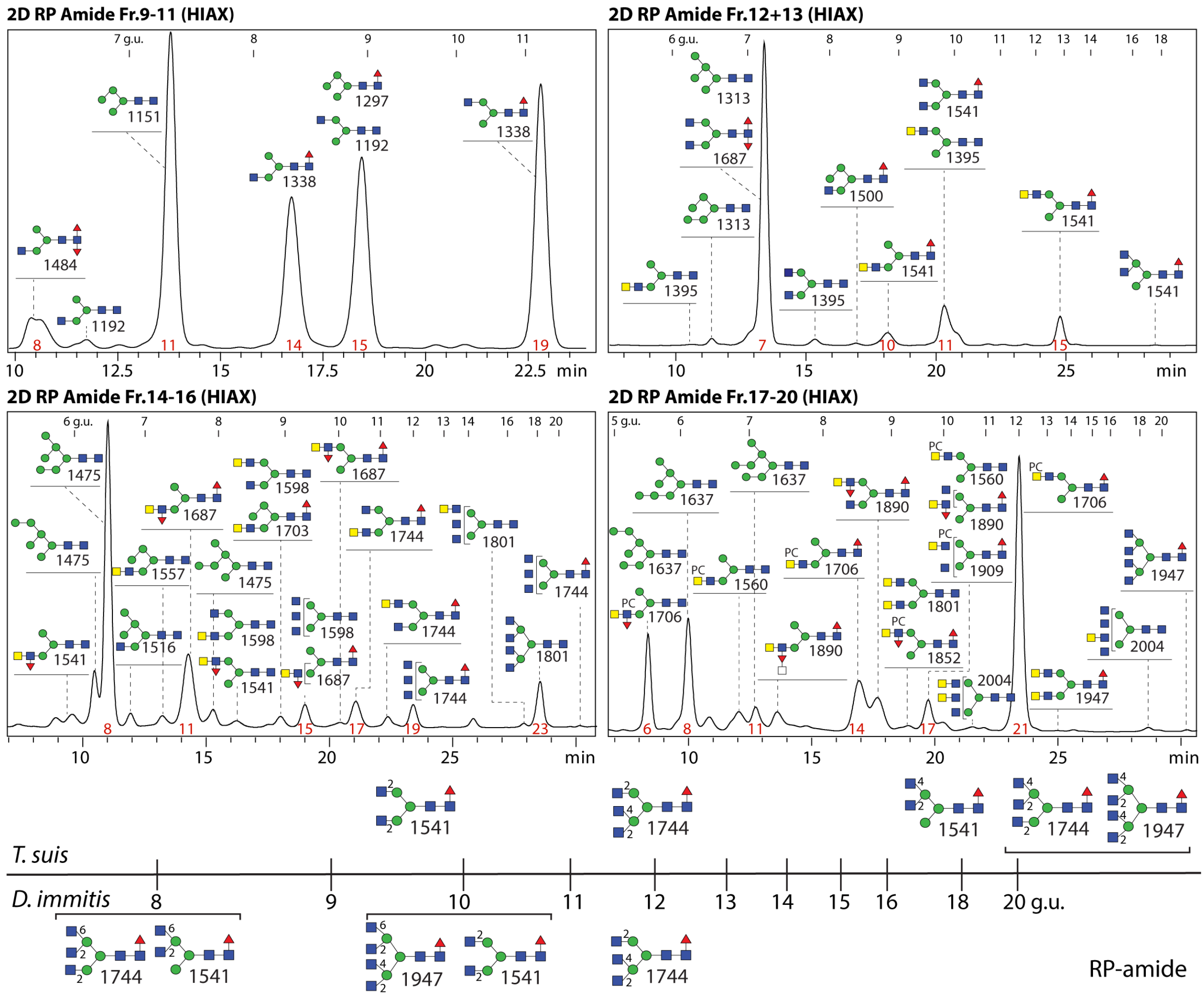


Analogous to our RP-amide data, studies on classical C18 RP-HPLC columns with pyridylaminated glycans showed that the contrasting earlier-eluting *m/z* 1744 isomers (2/2/6 and 2/4/2) eluted at 10.3 and 14.8 g.u., as compared to the typical *m/z* 1947 isomer (2/4/2/6) at 12.4 g.u.; the 'unit contribution' of lower and upper arm β1,4-GlcNAc was calculated as +2.1 and +1.3 g.u., whereas that for upper arm β1,6-GlcNAc is -2.1 g.u (Tomiya *et al.*, 1988, *Anal. Biochem*. 171, 73 and Tomiya & Takahashi, 1998, *Anal. Biochem*. 264, 204). Therefore, we hypothesise that the 'upper arm' GlcNAc residues are β1,2- and β1,4-linked and that no β1,6-linked GlcNAc is found in the *T. suis* N-glycome, compatible with the presence of an MGAT4 β1,4-*N*-acetylglucosaminyltransferase gene and apparent lack of one encoding an MGAT5 β1,6-*N*-acetylglucosaminyltransferase.

**Supplementary Figure 9. MALDI-TOF MS and MS/MS of lower molecular weight *T. suis* N-glycans.** (A and B) MALDI-TOF MS and MS/MS of two *m/z* 1687 isomers as well as respective enzymatic or chemical treatments to demonstrate the isomeric structures; the antennal fucose residues of the *m/z* 1687 isomers are sensitive to HF or almond α1,3/4-fucosidase; the underlying LacdiNAc motif is sensitive to chitinase or *C. elegans* GalNAc-specific HEX-4. The MS/MS comparison of the *m/z* 1687 glycans shows the difference in the ratio of the *m/z* 973 and 1135 Y-fragments depending on whether the fucosylated LacdiNAc (LDNF) motif is on the upper or lower arm. The MS/MS spectrum of the HF-sensitive *m/z* 1890 glycan is shown in ***Figure 6A***; for a comparison with the *m/z* 1687 glycan with a difucosylated core, refer to ***Figure 4A***. (C) MALDI-TOF MS and MS/MS of three 1706 isomers showing variations in the detected B fragments; the α-mannosidase sensitivity of one *m/z* 1706 structure suggests that the free terminal mannose is the α1,3-linked residue. Refer to ***Supplementary Figures 7 and 8*** for the elution positions of the HIAX fractions 26/27 and the 2D fractions 14-16/11, 17-20/6 and 17-20/21.

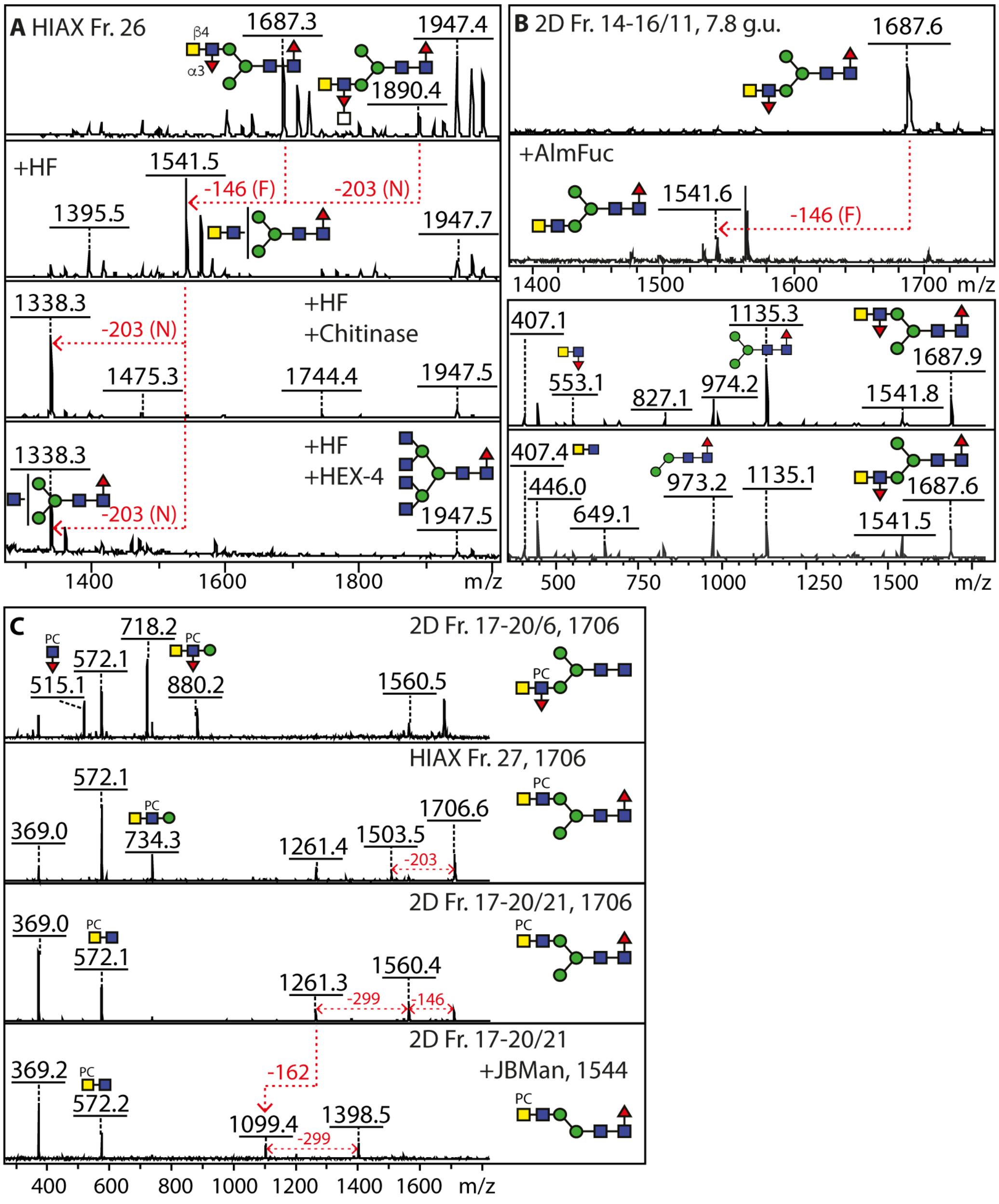

**Supplementary Figure 10. Enzymatic and chemical treatments of two isomeric glycans.** Two different HIAX fractions containing isomeric glycans of *m/z* 1852 were subject to different combinations of HF and hexosaminidase treatments. (A-D) The *m/z* 1852 glycan in HIAX fraction 31 was sensitive to HEX-4, HF and serial HF-HEX-4 treatment; together with the MS/MS data showing an *m/z* 515 fragment ($HexNAc_1Fuc_1PC_1$) and the sensitivity of the original glycan to HEX-4, specific for β1,4-*N*-acetylgalactosamine (Paschinger et al., 2023, *J. Biol. Chem*., 299, 103053), it is concluded that the PC residue is on the same GlcNAc as the antennal fucose and not on the terminal GalNAc. . (E-H) The *m/z* 1852 glycan in HIAX fraction 36 was sensitive to HF and serial HF-HEX-4 treatment, but not to HEX-4 alone; together with the MS/MS data showing the absence of *m/z* 515 fragment ($HexNAc_1Fuc_1PC_1$), it is concluded that the PC residue is on the terminal GalNAc and not on the sub-terminal fucose-substituted GlcNAc. A co-eluting phosphate-modified glycan of *m/z* 1621 displays a similar sensitivity and it is concluded that the phosphate residue substitutes the terminal GalNAc. Note that while PC-modified glycans tend to ionize as $[M+H]^+$, but that the digested glycans were partly detected as $[M+Na]^+$ and fragmented as $[M+H]^+$. The phosphorylcholine residue is assumed to be 6-linked considering previous data on nematode glycolipids (Gerdt *et al.*, 2003, *Biochem. J.* 369, 89-102), as well as considering the antennal GlcNAc residue in the Fraction 31 *m/z* 1852 carries 2-*N*-acetyl-, 3-fucosyl and 4-*N*-acetylgalactosaminyl-substitutions, leaving only the C6 available for the phosphodiester modification. Furthermore, previous Q-TOF CAD-MS/MS and FAB-MS data have indicated that phosphorylcholine is 6-linked to HexNAc residues of perdeuteroacetylated *C. elegans* N-glycans (Haslam *et al.*, 2002, *Biochem. Soc. Symp*. 69, 117-134).

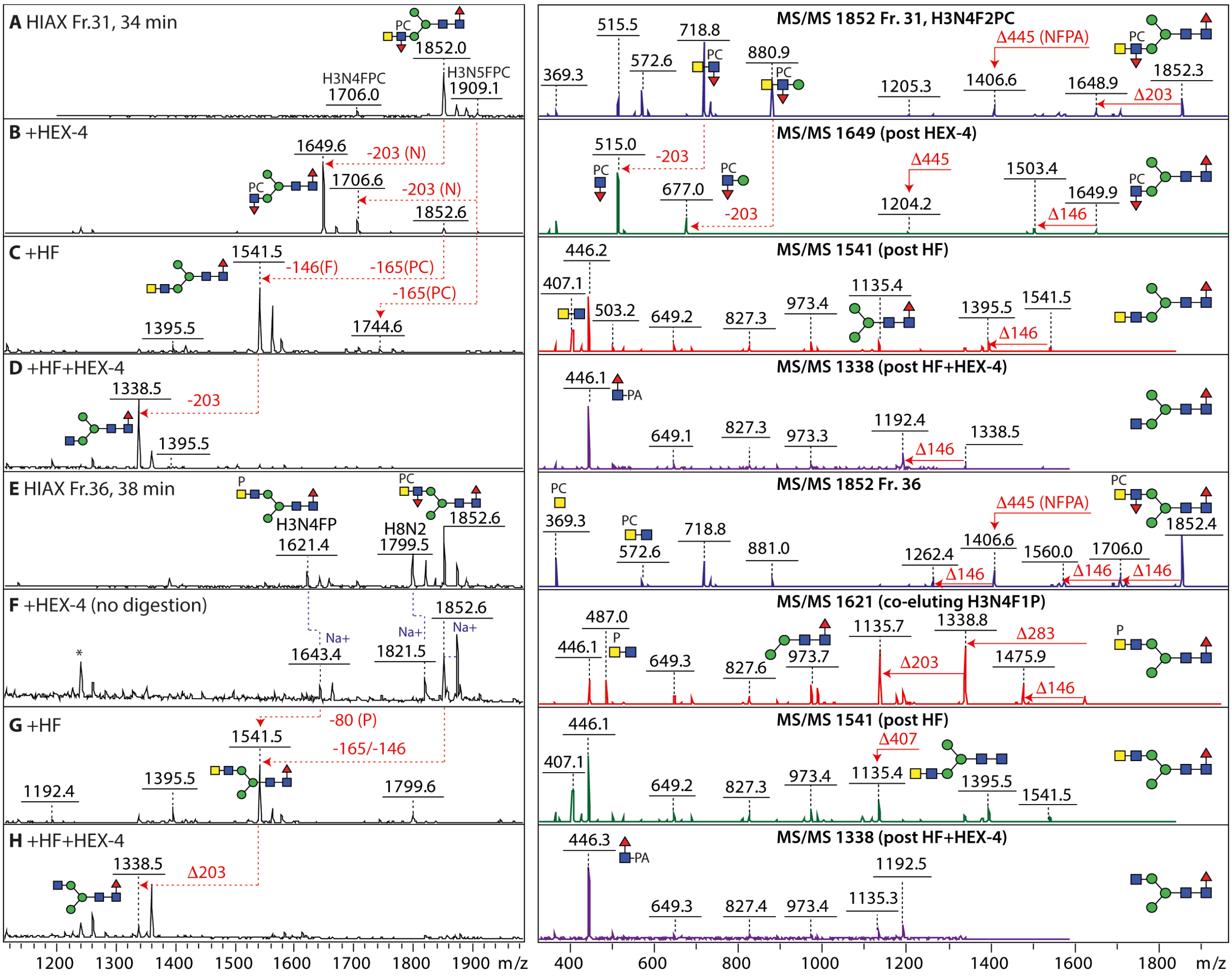

**Supplementary Figure 11. 2D-HPLC and chemical/enzymatic treatment of *T. suis* N-glycans of around 3200-3400 Da.** The HIAX fractions containing glycans of ca. 3200-3400 Da were treated with HF and chitinase and then pooled prior to another round of RP-amide HPLC (see also MS and MS/MS). Subsequent jack bean hexosaminidase treatment resulted in peaks co-eluting with normal forms of $Man_3GlcNAc_2Fuc_{0\text{-}1}$, indicative that the original glycans are tetra-antennary structures. However, the intermediate late-eluting *m/z* 1947 does not correspond to the *m/z* 1947 isomer from *Dirofilaria*, suggesting that the *T. suis* tetra-antennary structure may not be one carrying two β1,2, one β1,4 and one β1,6-antennae (see note also for ***Supplementary Figure 8***), despite a 'normal' $Man_3GlcNAc_2Fuc_{0\text{-}1}$ core region, but rather possess two β1,2 and two β1,4 antennae. Note that while PC-modified glycans tend to ionize as $[M+H]^+$, the digested glycans were detected as $[M+Na]^+$ and fragmented as $[M+H]^+$.

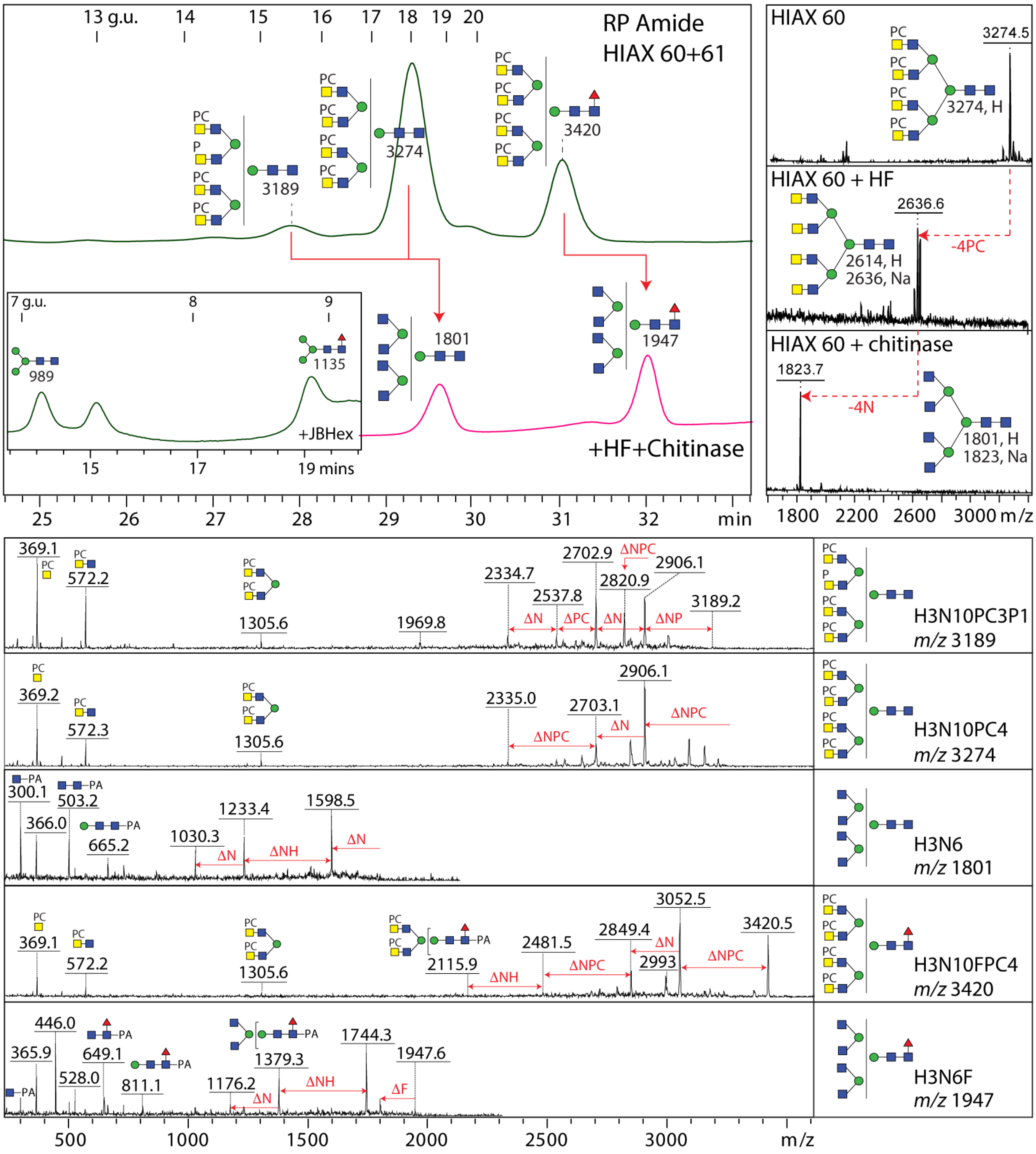

**Supplementary Figure 12. HPLC and chemical treatment of *T. suis* N-glycans of 3900-4700 Da.** Limited MS/MS data are available on larger N-glycans; the example for *m/z* 4499 suggests the occurrence of phosphorylcholine-modified fucosylated LacdiNAc – a tentative structure is shown. The HIAX fractions 70-76 containing glycans of ca. 3900-4700 Da were analysed by linear MALDI-TOF MS, pooled and treated with HF prior to RP-amide HPLC, resulting in glycans of *m/z* 2354, 2557 and 2760 as defined by reflector-mode MALDI-TOF MS; note that the average masses are some 2-3 Da greater than the theoretical monoisotopic masses listed ***in Supplementary Tables 1 and 2***. Based on the observed average masses of the original glycans and the accurate masses and MS/MS of the HF-treated products, it is concluded that these glycans are maximally tetraantennary, carrying three or four antennal fucose residues and up to eight phosphorylcholine residues as compared to the tetra-antennary glycans carrying four phosphorylcholine residues in HIAX fractions 60 and 61 (see ***Supplementary Figure 11***). As the dominant B-fragment of the HF digestion product is *m/z* 407, it is concluded that the antennae are all based on LacdiNAc and not on longer chito-oligomers as found additionally in filarial species or in *C. elegans* (Martini et al., 2019, *Nat. Commun*. 10, 75; Wilson et al., 2023, *Mol. Cell. Proteomics* 22, 100505). The glycans in this region correspond to the largest glycans on the glycan array (Ts-FMAPA fractions 25, 26 and 27; see ***Supplementary Figure 4***).

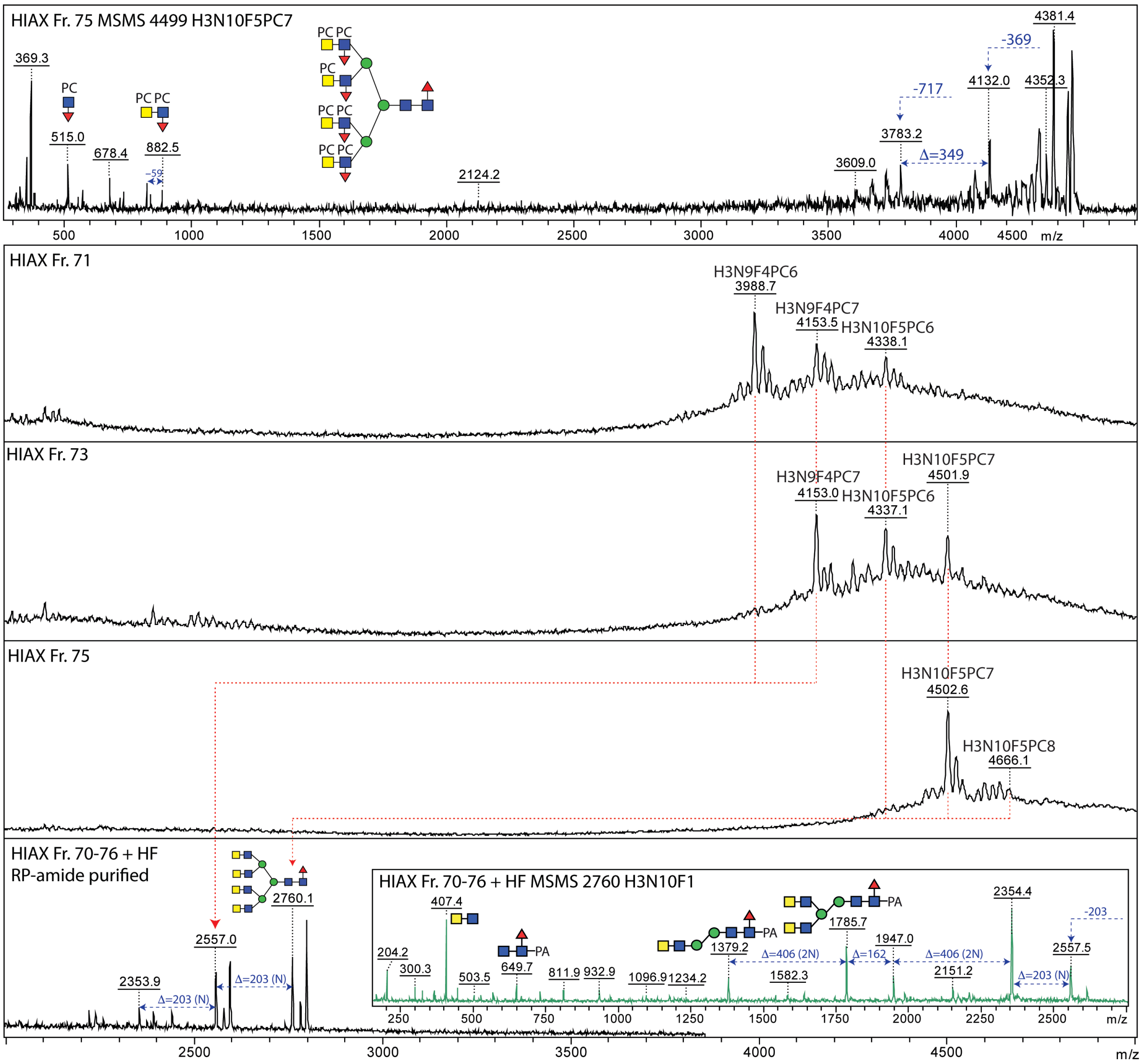


**Supplementary Figure 13. Enzymatic and chemical treatments of two isomeric glycans.** Two different HIAX fractions containing isomeric glycans of *m/z* 2055 were subject to different combinations of HF, fucosidase and hexosaminidase treatments. (A-D) A simple biantennary form of *m/z* 2055 lost (i) a core fucose, but not the antennal fucose, upon bovine fucosidase treatment (loss of 146 Da, but no change in the B-fragments), (ii) the antennal PC-residue and the antennal fucose residue upon HF treatment, resulting in an *m/z* 1744 lacking the characteristic PC-containing B-fragments, but rather an intense B2 LacdiNAc fragment and an intense Y1 core GlcNAc$_1$Fuc$_1$-PA fragment; subsequent treatment with *C. elegans* GalNAc-specific HEX-4 resulted in (iii) loss of the B2 LacdiNAc fragment; note that the PC residue is indicated as modifying the GalNAc residue, due to the lack of an *m/z* 515 (HexNAc$_1$Fuc$_1$PC$_1$). The co-eluting *m/z* 2093 glycan was also sensitive to serial HF and HEX-4 treatment, losing two antennal fucose and then two antennal GalNAc residues. (E-H) A simple hybrid form of m/z 2055 lost (i) a core fucose, but not the antennal fucose, upon bovine fucosidase treatment (loss of 146 Da, but no change in the B-fragments), (ii) was resistant to jack bean hexosaminidase unlike certain co-eluting structures with unsubstituted antennal GlcNAc residues and (ii) lost the antennal PC-residue, the antennal fucose and a HexNAc residue upon HF treatment, resulting in an *m/z* 1541 lacking the characteristic PC-containing B-fragments, but rather an intense B2 LacdiNAc fragment and an intense Y1 core GlcNAc$_1$Fuc$_1$-PA fragment. Note that the *m/z* 921 and 1083 B2 and B3 fragments (Hex$_{0\text{-}1}$HexNAc$_3$Fuc$_1$PC$_1$) lost upon HF treatment indicate the HexNAc-substitution of fucose absent from the co-eluting *m/z* 1852 glycan. Various co-eluting PC-modified and phosphorylated glycans were also sensitive to HF treatment as indicated. Only the [M+H]$^+$ ions are annotated and fragmented, but [M+Na]$^+$ and [M+K]$^+$ were also detected.

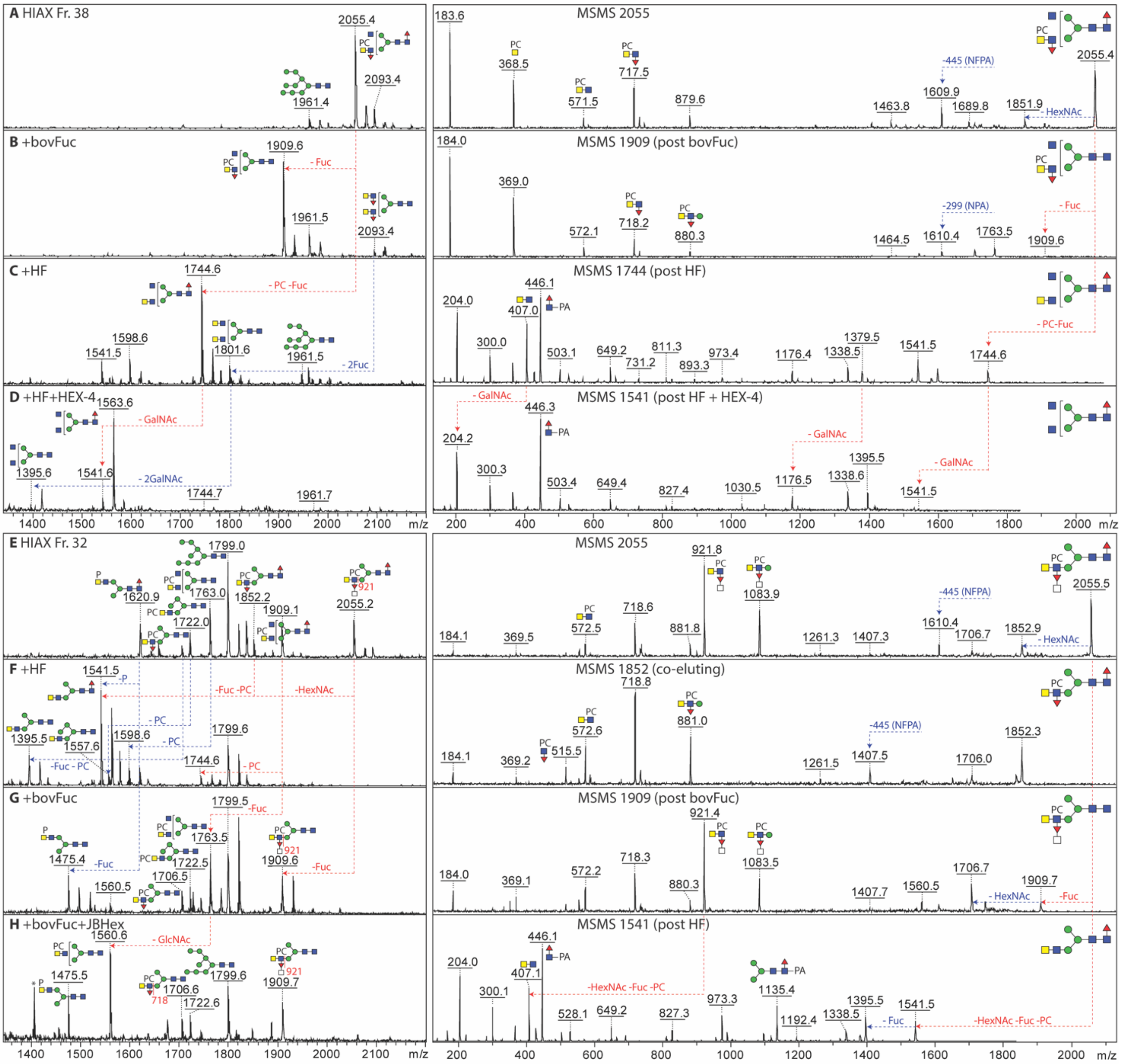

**Supplementary Figure 14. Examples of isomeric and isobaric tri- and tetra-antennary glycans.** (A-C) Three isomers of *m/z* 2684 in different 2D-HPLC fractions distinguished on the basis of the MALDI-TOF MS/MS B3 fragments showing the presence of PC-LacdiNAc units on the same or different core α-mannose residue. (D-F) Three isomers of *m/z* 2773 in different 2D-HPLC fractions distinguished on the basis of the MALDI-TOF MS/MS B3 fragments showing the presence of PC-FucLacdiNAc units, with and without a HexNAc-substitution of fucose, on the same or different core α-mannose residue. (G-J) Three isobaric structures of *m/z* 2995 in different 2D-HPLC fractions distinguished on the basis of the MALDI-TOF MS/MS B and Y fragments showing the presence of either fucosylated LacdiNAc, with and without a HexNAc-substitution of fucose, or phosphorylcholine-modified LacdiNAc; HF treatment of the isobar with the HexNAc-substitution is shown in the inset in panel H and the resulting MS/MS shows the replacement of the *m/z* 553 and 756 B2 fragments ($HexNAc_{2-3}Fuc_1$) by ones solely at *m/z* 407 (LacdiNAc), comparable to data shown for PC-modified glycans in ***Figure 6 D-F*** and ***Supplementary Figure 13 E-H***. The lack of a $HexNAc_3$ (*m/z* 610) fragment after HF treatment is an indication that the third HexNAc on the one antenna in panel G is indeed substituting the HF-sensitive α1,3-fucose modifying the LacdiNAc motif. All glycans fragmented as $[M+H]^+$.

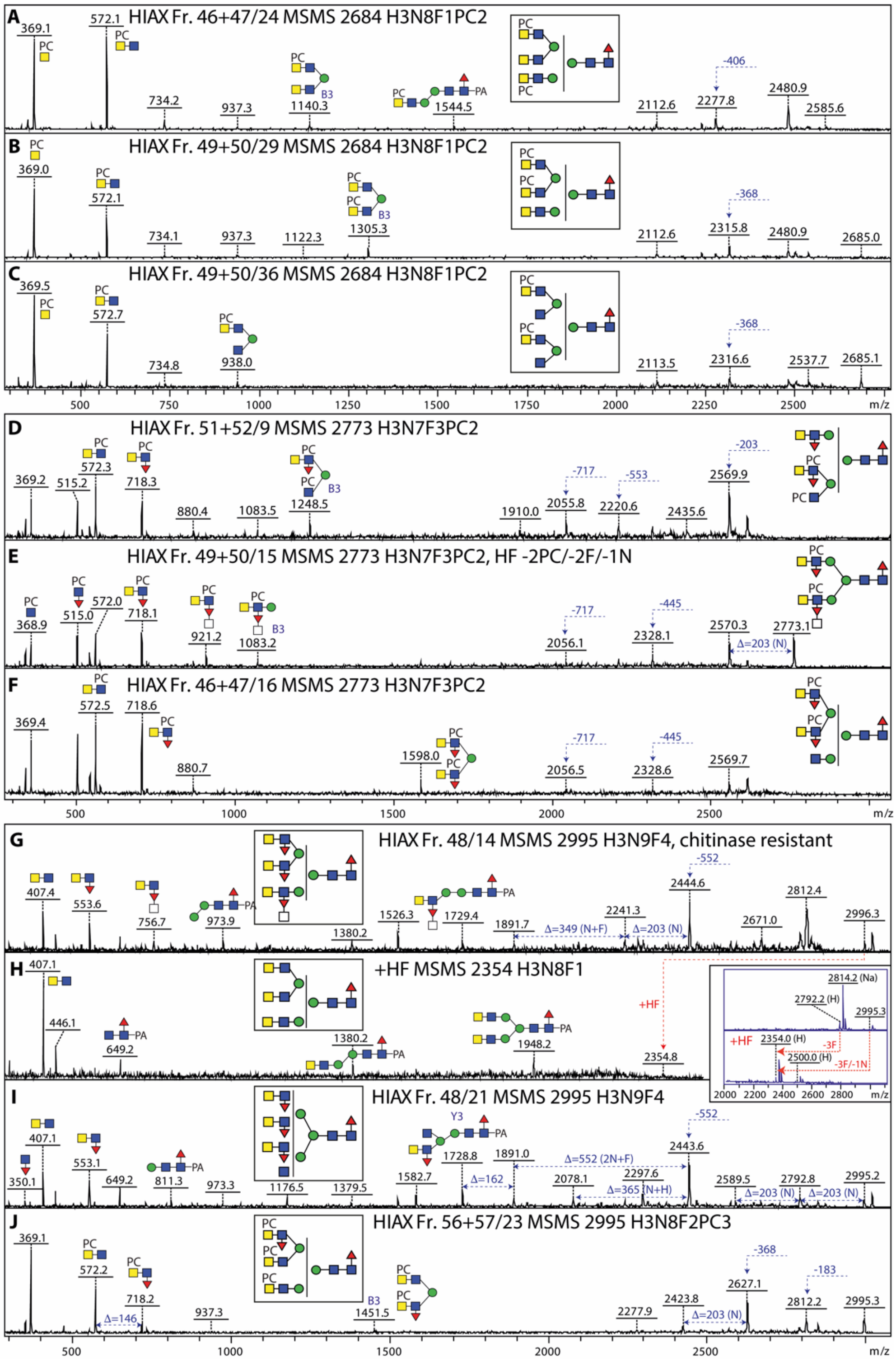

**Supplementary Figure 15. Chemical treatment of biantennary N-glycans with HexNAc-substituted fucose.** (A and B) The 2D-HPLC fraction containing glycans of *m/z* 2773 and 2976 was treated with hydrofluoric acid, resulting in a major product of *m/z* 1947. Both these isomers have either one or two HexNAc-substituted fucose residues in the context of PC-modified $HexNAc_2$ motifs, which are replaced by an *m/z* 407 $HexNAc_2$ B-fragment. As indicated in ***Supplementary Figure 13***, lower molecular weight glycans with the *m/z* 921 fragment were also HF treated, losing a HexNAcFuc motif as well as a PC moiety, replaced by a B fragment at *m/z* 407; as shown in ***Supplementary Figure 9***, the similar motif lacking the phosphorylcholine was also HF sensitive (loss of HexNAcFuc) and a GalNAc residue is lost upon subsequent treatment with HEX-4, indicative that the HexNAcFuc motif is a modification of a LacdiNAc antenna. For other examples of HexNAc-substituted fucose residues, refer to ***Figures 6 and 7*** and ***Supplementary Figures 13, 14 and 16***. The linkage and the nature of the HexNAc residue are unknown, but this modification has not been detected in any nematode to date.

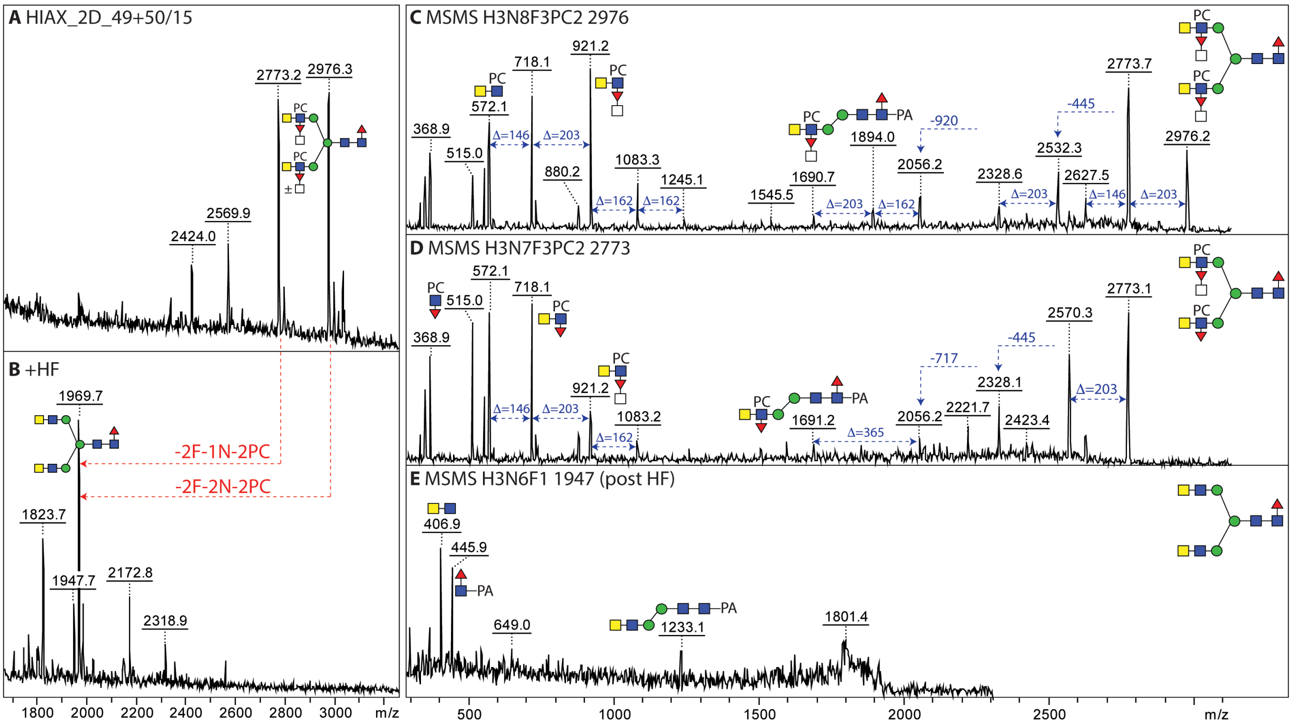

**Supplementary Figure 16. N-glycans with phosphorylcholine and/or HexNAc-substituted fucose.** (A-D) MALDI-TOF MS/MS of hybrid, tri- and tetra-antennary glycans modified with phosphorylcholine; the pattern of B fragments and losses indicate a terminal position for the PC residue, while the B3 ions show disubstitution of a core $\alpha$-mannose. (E) MALDI-TOF MS/MS of a triantennary isobar of *m/z* 3401 (i.e., different composition for an almost identical mass) carrying three HexNAc-substitutions of fucosylated LacdiNAc motifs; losses of 552 and 755 Da and B ions at *m/z* 553 and 756 correspond to fucosylated LacdiNAc and its HexNAc-substituted form; for other examples, including the effect of HF treatment on this motif, see ***Figure 6*** and ***Supplementary Figures 13-15***. (F-I) MALDI-TOF MS/MS of two isomers each of triantennary glycans modified with phosphorylcholine and fucosylated LacdiNAc. The different pattern of smaller B fragments is indicative for a different position of one of the PC residues, i.e., either on GlcNAc and Man residues or juxtaposed HexNAc residues or the HexNAc-substitution.

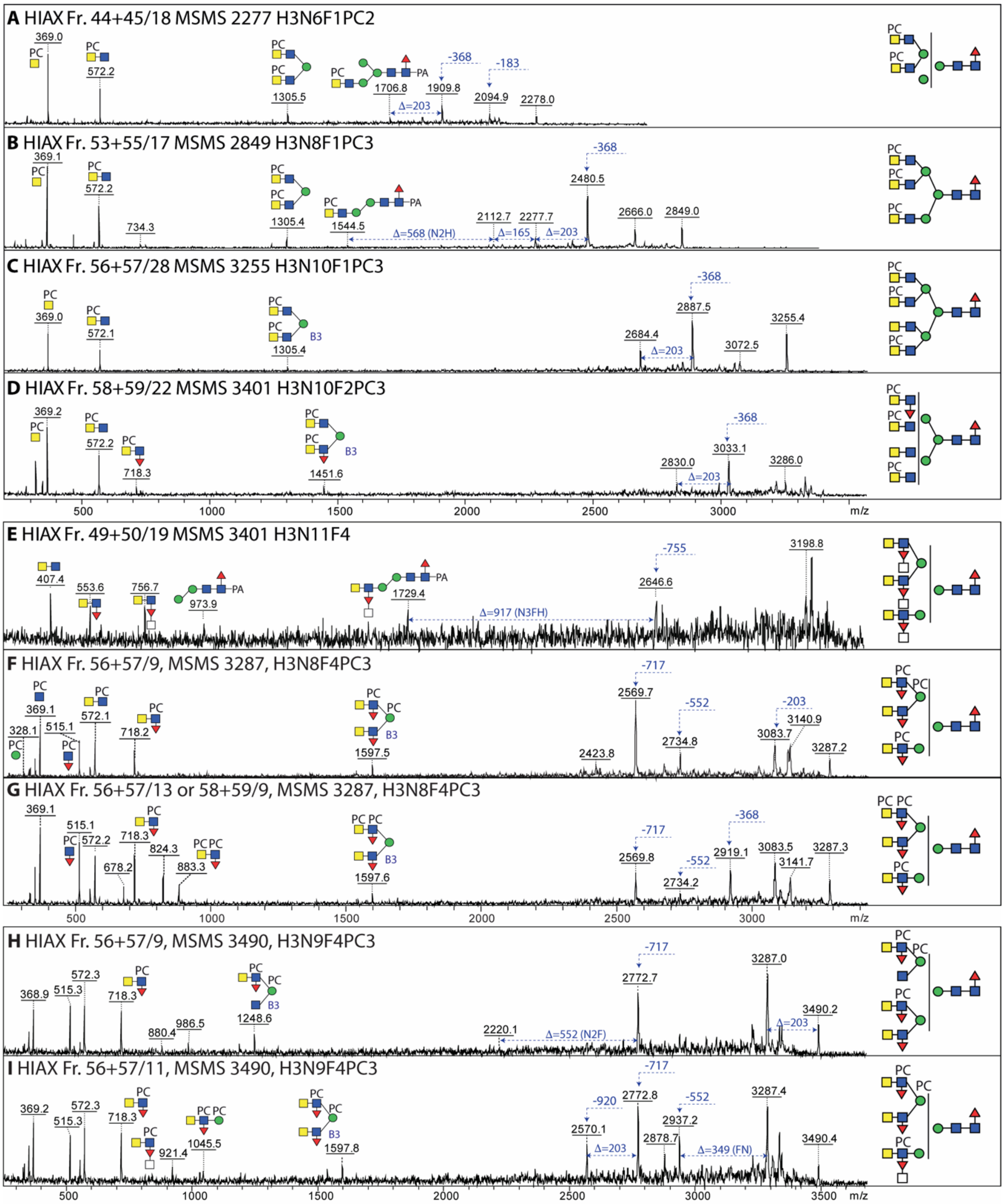

**Supplementary Figure 17. Phosphorylated and phosphorylcholine-modified N-glycans.** (A-J) MALDI-TOF MS/MS of various glycans were detected which were 85 Da smaller than a phosphorylcholine-modified structure. Some of these glycans were also detected in negative ion mode (B) and showed the loss of 283 Da in positive ion mode. It was concluded that these structures are phosphorylated in one position rather than PC-modified. The patterns of B-ion fragments and losses of either 283 (GalNAcP) or 368 Da (GalNAcPC) are indicative of the antennal motifs and branching. All glycans were fragmented as $[M+H]^+$, other than the *m/z* 2190/2192 glycan also fragmented as $[M-H]^-$. B-ion fragments containing only phosphate and not phosphorylcholine are only observed here in negative mode, due to the more efficient ionization of the phosphorylcholine-containing motifs in positive mode.

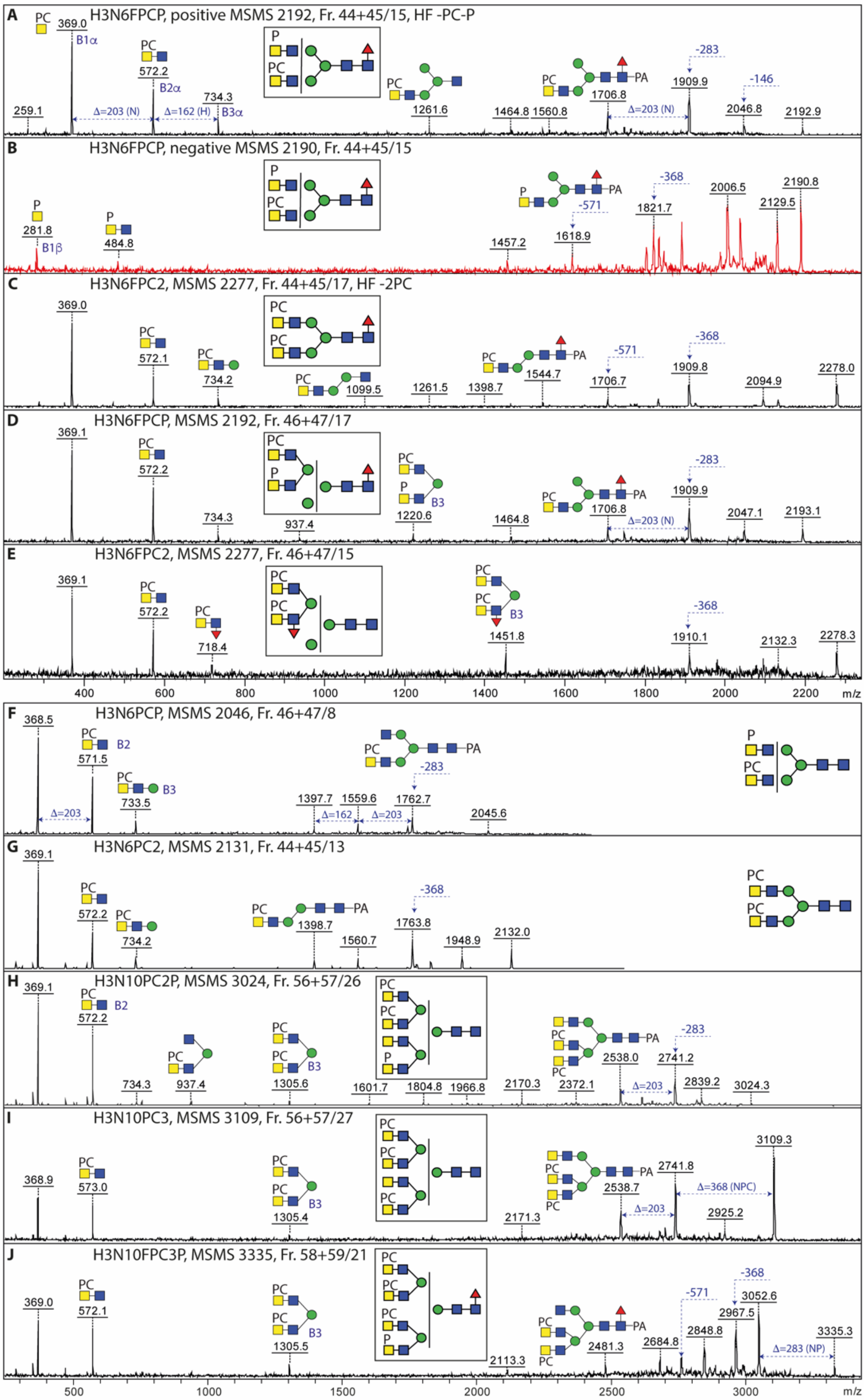


For positive mode MS/MS of a solely phosphorylated HF sensitive structure (*m/z* 1621), showing a $HexNAc_2P_1$ B ion in positive mode, see *Supplementary Figure 10*.

MS/MS of a further *m/z* 2277 isomer is shown in *Supplementary Figure 16A*.

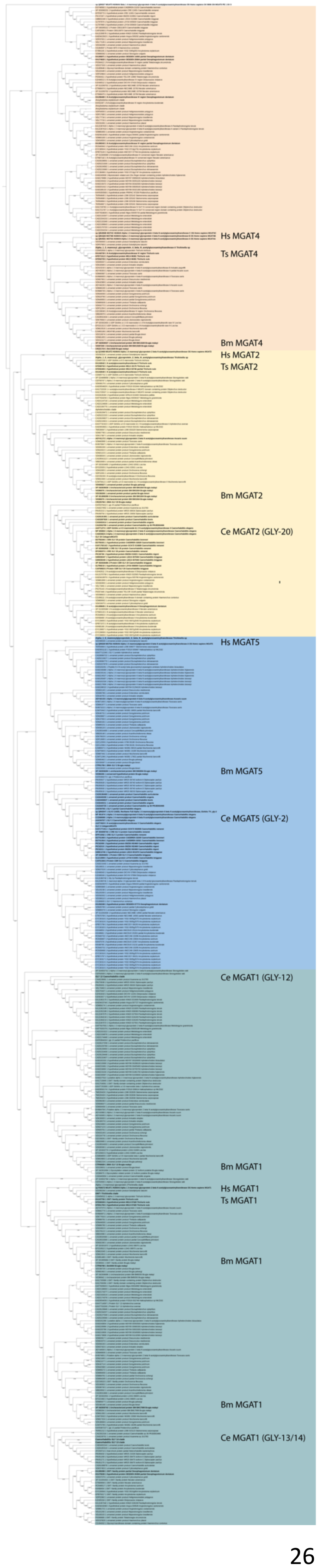


**Supplementary Figure 18. Phylogeny of N-glycan branching β1,2/-4/-6-*N*-acetylglucosaminyltransferases in nematodes.** As described in the relevant methods section, human and *C. elegans* MGAT1, MGAT2, MGAT3 (human only; Hs), MGAT4 (human only) and MGAT5 sequences were used to search for nematode homologues. For some genes from some species, multiple entries were found; some clades of related sequences are collapsed. The resulting phylogenetic search and tree show that *T. suis* possesses MGAT1, MGAT2 and MGAT4 homologues (Ts MGAT1, Ts MGAT2 and Ts MGAT4), but no MGAT5 homologue could be detected, which contrasts with *C. elegans* which has three MGAT1 (GLY-12, -13 and -14; Ce), one MGAT2 (GLY-20) and one MGAT5 homologues (GLY-2), but no MGAT4; like many other nematodes, the filarial species *Brugia malayi* has at least one homologue each of MGAT1, MGAT2, MGAT4 and MGAT5 (annotated as Bm MGAT1, etc.). No bisecting GlcNAc has been detected in nematodes to date (only bisecting galactose in *C. elegans*), which correlates with a lack of obvious MGAT3 homologues. *C. elegans* has only maximally tri-antennary N-glycans, whereas species such as *Dirofilaria immitis* and *Trichinella spiralis* have 'standard' tetra-antennary N-glycans. Due to the unusual elution properties of the *T. suis* tetra-antennary N-glycans (see ***Supplementary Figure 8***), a potential explanation is that its MGAT4 can modify both core α-mannose residues with a β1,4-GlcNAc and not just the 'lower arm' α1,3-mannose and so replaces the activity of the absent β1,6-specific MGAT5; however, the predictions of numbers of homologues and their activities requires experimental verification. For a full-sized version of the figure, refer to the separate pdf file.

**Supplementary Table 1: List of theoretical masses for detected *T. suis* N-glycan structures.** Abbreviated compositions of the form $Hex_{3-5}HexNAc_{2-10}Fuc_{0-5}P_{0-1}PC_{0-9}$ and $Hex_{1-10}HexNAc_2Fuc_{0-1}$. *m/z* as $[M+H]^+$ for N-glycans labelled with 2-aminopyridine (PA) or phosphorylcholine-modified N-glycans labelled with FMAPA, *m/z* as $[M+Na]^+$ for free N-glycans, N-glycans labelled with 2-aminopyridine (PA) or with FMAPA are given. The numbers of detected potential isomers for the PA-labelled N-glycans and the presence or absence of FMAPA-labelled N-glycans in HPLC fractions are also indicated. Isobaric PC/Fuc- or Fuc-containing compositions are highlighted in blue or red.

| Composition | Free [M+Na]+ | PA-[M+H]$^+$ | PA-[M+Na]$^+$ | FMAPA [M+H]$^+$ | FMAPA [M+Na]$^+$ | HPLC PA | HPLC FMAPA |
|---|---|---|---|---|---|---|---|
| H1N2F | 755,27 | 811,34 | 833,32 | | 1063,43 | 1 | nd |
| H2N2 | 771,27 | 827,34 | 849,32 | | 1079,43 | 2 | ✔ |
| H2N2F | 917,32 | 973,39 | 995,37 | | 1225,48 | 2 | ✔ |
| H3N2 | 933,32 | 989,39 | 1011,37 | | 1241,48 | 1 | ✔ |
| H2N3 | 974,34 | 1030,41 | 1052,39 | | 1282,5 | 1 | ✔ |
| H3N2F | 1079,38 | 1135,45 | 1157,43 | | 1387,54 | 1 | ✔ |
| H4N2 | 1095,37 | 1151,44 | 1173,42 | | 1403,53 | 1 | ✔ |
| H3N3 | 1136,4 | 1192,47 | 1214,45 | | 1444,56 | 2 | ✔ |
| H3N2F2 | 1225,43 | 1281,5 | 1303,48 | | 1533,59 | 1 | ✔ |
| H4N2F | 1241,43 | 1297,5 | 1319,48 | | 1549,59 | 1 | ✔ |
| H5N2 | 1257,42 | 1313,49 | 1335,47 | | 1565,58 | 2 | ✔ |
| H3N3F | 1282,46 | 1338,53 | 1360,51 | | 1590,62 | 2 | ✔ |
| H4N3 | 1298,38 | 1354,45 | 1376,43 | | 1606,54 | 1 | nd |
| H3N4 | 1339,48 | 1395,55 | 1417,53 | | 1647,64 | 4 | ✔ |
| H6N2 | 1419,48 | 1475,55 | 1497,53 | | 1727,64 | 3 | ✔ |
| H3N3F2 | 1428,51 | 1484,58 | 1506,56 | | 1736,67 | 1 | ✔ |
| H4N3F | 1444,51 | 1500,58 | 1522,56 | | 1752,67 | 1 | nd |
| H5N3 | 1460,5 | 1516,57 | 1538,55 | | 1768,66 | 1 | nd |
| H3N4F | 1485,53 | 1541,6 | 1563,58 | | 1793,69 | 6 | ✔ |
| H4N4 | 1501,53 | 1557,6 | 1579,58 | | 1809,69 | 1 | nd |
| H3N4PC | 1504,53 | 1560,6 | 1582,58 | 1790,71 | 1812,69 | 2 | ✔ |
| H3N5 | 1542,56 | 1598,63 | 1620,61 | | 1850,72 | 3 | nd |
| H3N4FP | 1565,5 | 1621,57 | 1643,55 | 1851,68 | 1873,66 | 1 | nd |
| H7N2 | 1581,53 | 1637,6 | 1659,58 | | 1889,69 | 3 | ✔ |
| H3N4F2 | 1631,59 | 1687,66 | 1709,64 | | 1939,75 | 5 | ✔ |
| H4N4F | 1647,59 | 1703,66 | 1725,64 | | 1955,75 | 1 | nd |
| H3N4FPC | 1650,59 | 1706,66 | 1728,64 | 1936,77 | 1958,75 | 3 | ✔ |
| H4N4PC | 1666,59 | 1722,66 | 1744,64 | 1952,77 | 1974,75 | 1 | nd |
| H3N5F | 1688,61 | 1744,68 | 1766,66 | | 1996,77 | 4 | ✔ |
| H3N5PC | 1707,61 | 1763,68 | 1785,66 | 1993,79 | 2015,77 | 1 | nd |
| H8N2 | 1743,58 | 1799,65 | 1821,63 | | 2051,74 | 2 | ✔ |
| H3N6 | 1745,64 | 1801,71 | 1823,69 | | 2053,8 | 2 | ✔ |
| H3N4F2PC | 1796,65 | 1852,72 | 1874,7 | 2082,83 | 2104,81 | 1 | ✔ |
| H3N4FPC2 | 1815,65 | 1871,72 | 1893,7 | 2101,83 | 2123,81 | nd | ✔ |
| H3N5F2 | 1834,67 | 1890,74 | 1912,72 | | 2142,83 | 2 | ✔ |
| H3N5FPC | 1853,67 | 1909,74 | 1931,72 | 2139,85 | 2161,83 | 1 | ✔ |
| H3N6F | 1891,69 | 1947,76 | 1969,74 | | 2199,85 | 2 | ✔ |
| H9N2 | 1905,64 | 1961,71 | 1983,69 | | 2213,8 | 1 | ✔ |
| H3N6PC | 1910,69 | 1966,76 | 1988,74 | 2196,87 | 2218,85 | nd | ✔ |

| | | | | | | | |
|---|---|---|---|---|---|---|---|
| H3N7 | 1948,72 | 2004,79 | 2026,77 | | 2256,88 | 2 | ✔ |
| H3N4F2PC2 | 1961,7 | 2017,77 | 2039,75 | 2247,88 | 2269,86 | 1 | ✔ |
| H3N6PC-P | 1990,66 | 2046,73 | 2068,71 | 2276,84 | 2298,82 | 2 | nd |
| H3N5F2PC | 1999,73 | 2055,8 | 2077,78 | 2285,91 | 2307,89 | 2 | ✔ |
| H3N5FPC2 | 2018,73 | 2074,8 | 2096,78 | 2304,91 | 2326,89 | 1 | ✔ |
| H3N6F2 | 2037,75 | 2093,82 | 2115,8 | | 2345,91 | 1 | ✔ |
| H3N6FPC | 2056,75 | 2112,82 | 2134,8 | 2342,93 | 2364,91 | 2 | ✔ |
| H10N2 | 2067,69 | 2123,76 | 2145,74 | | 2375,85 | 1 | ✔ |
| H3N6PC2 | 2075,75 | 2131,82 | 2153,8 | 2361,93 | 2383,91 | 1 | ✔ |
| H3N7F | 2094,77 | 2150,84 | 2172,82 | | 2402,93 | 1 | ✔ |
| H3N7PC | 2113,89 | 2169,96 | 2191,94 | 2400,07 | 2422,05 | nd | ✔ |
| H3N6FPCP | 2136,72 | 2192,79 | 2214,77 | 2422,90 | 2444,88 | 2 | nd |
| H5N4FPC2 | 2139,93 | 2196,00 | 2217,98 | 2426,11 | 2448,09 | 1 | nd |
| H3N8 | 2151,79 | 2207,86 | 2229,84 | | 2459,95 | nd | ✔ |
| H3N5F2PC2 | 2164,78 | 2220,85 | 2242,83 | 2450,96 | 2472,94 | 1 | ✔ |
| H3N6F3 | 2183,81 | 2239,88 | 2261,86 | | 2491,97 | 1 | ✔ |
| H3N6F2PC | 2202,81 | 2258,88 | 2280,86 | 2488,99 | 2510,97 | 2 | ✔ |
| H3N6FPC2 | 2221,8 | 2277,87 | 2299,85 | 2507,98 | 2529,96 | 3 | ✔ |
| H3N7PC2 | 2278,83 | 2334,9 | 2356,88 | 2565,01 | 2586,99 | 1 | ✔ |
| H3N8F | 2297,85 | 2353,92 | 2375,90 | | 2606,01 | nd | ✔ |
| H3N5F3PC2 | 2310,96 | 2367,03 | 2389,01 | 2597,14 | 2619,12 | 1 | nd |
| H3N8PC | 2316,85 | 2372,92 | 2394,9 | 2603,03 | 2625,01 | nd | ✔ |
| H3N6F3PC | 2348,86 | 2404,93 | 2426,91 | 2635,04 | 2657,02 | 1 | ✔ |
| H3N6F2PC2 | 2367,86 | 2423,93 | 2445,91 | 2654,04 | 2676,02 | 6 | ✔ |
| H3N6FPC3 | 2386,86 | 2442,93 | 2464,91 | 2673,04 | 2695,02 | 1 | nd |
| H3N7F3 | 2386,89 | 2442,96 | 2464,94 | | 2695,05 | 1 | nd |
| H3N7FPC2 | 2424,88 | 2480,95 | 2502,93 | 2711,06 | 2733,04 | 2 | nd |
| H3N8FPC | 2462,91 | 2518,98 | 2540,96 | 2749,09 | 2771,07 | 1 | ✔ |
| H3N8PC2 | 2481,91 | 2537,98 | 2559,96 | 2768,09 | 2790,07 | 4 | ✔ |
| H3N6F3PC2 | 2513,92 | 2569,99 | 2591,97 | 2800,10 | 2822,08 | 6 | ✔ |
| H3N9PC | 2520,05 | 2576,12 | 2598,1 | 2806,23 | 2828,21 | 2 | nd |
| H3N6F2PC3 | 2532,92 | 2588,99 | 2610,97 | 2819,10 | 2841,08 | 3 | ✔ |
| H3N7F3PC | 2552,02 | 2608,09 | 2630,07 | 2838,20 | 2860,18 | 1 | nd |
| H3N10 | 2557,95 | 2614,02 | 2636,00 | | 2866,11 | 1 | ✔ |
| H3N7F2PC2 | 2570,94 | 2627,01 | 2648,99 | 2857,12 | 2879,1 | 3 | nd |
| H3N8F3 | 2589,97 | 2646,04 | 2668,02 | | 2898,13 | 1 | nd |
| H3N8F2PC | 2609,05 | 2665,12 | 2687,1 | 2895,23 | 2917,21 | 1 | nd |
| H3N8FPC2 | 2627,96 | 2684,03 | 2706,01 | 2914,14 | 2936,12 | 5 | ✔ |
| H3N8PC3 | 2646,96 | 2703,03 | 2725,01 | 2933,14 | 2955,12 | nd | ✔ |
| H3N9FPC | 2666,15 | 2722,22 | 2744,2 | 2952,33 | 2974,31 | 2 | nd |
| H3N6F3PC3 | 2678,98 | 2735,05 | 2757,03 | 2965,16 | 2987,14 | 3 | ✔ |
| H3N9PC2 | 2684,98 | 2741,05 | 2763,03 | 2971,16 | 2993,14 | 3 | nd |
| H3N10F | 2704,01 | 2760,08 | 2782,06 | | 3012,17 | 1 | nd |
| H3N7F3PC2 | 2717,0 | 2773,07 | 2795,05 | 3003,16 | 3025,16 | 5 | ✔ |
| H3N10PC | 2723,01 | 2779,08 | 2801,06 | 3009,19 | 3031,17 | 1 | nd |
| H3N7F2PC3 | 2736,0 | 2792,08 | 2814,05 | 3022,18 | 3044,16 | 1 | |
| H3N8F4 | 2736,03 | 2792,1 | 2814,08 | | 3044,19 | 3 | nd |
| H3N8F3PC | 2755,02 | 2811,09 | 2833,07 | 3041,20 | 3063,18 | 1 | nd |

| | | | | | | | |
|---|---|---|---|---|---|---|---|
| H3N8FPC3 | 2793,02 | 2849,09 | 2871,07 | 3079,20 | 3101,18 | 1 | ✓ |
| H3N9F3 | 2793,05 | 2849,12 | 2871,1 | | 3101,21 | 3 | nd |
| H3N9FPC2 | 2831,04 | 2887,11 | 2909,09 | 3117,22 | 3139,2 | 1 | nd |
| H3N6F3PC4 | 2844,03 | 2900,1 | 2922,08 | 3130,21 | 3152,19 | 2 | ✓ |
| H3N9PC3 | 2850,04 | 2906,11 | 2928,09 | 3136,22 | 3158,2 | 1 | ✓ |
| H3N10FPC | 2869,07 | 2925,14 | 2947,12 | 3155,25 | 3177,23 | 1 | nd |
| H3N7F3PC3 | 2882,06 | 2938,13 | 2960,11 | 3168,24 | 3190,22 | 1 | ✓ |
| H3N8F5 | 2882,06 | 2938,13 | 2960,11 | | 3190,22 | 1 | nd |
| H3N8F4PC | 2901,08 | 2957,15 | 2979,13 | 3187,26 | 3209,24 | 2 | ✓ |
| H3N8F3PC2 | 2920,08 | 2976,15 | 2998,13 | 3206,26 | 3228,24 | 4 | ✓ |
| H3N8F2PC3 | 2939,08 | 2995,15 | 3017,13 | 3225,26 | 3247,24 | 2 | ✓ |
| H3N9F4 | 2939,11 | 2995,18 | 3017,16 | | 3247,27 | 3 | nd |
| H3N10PC2P | 2968,03 | 3024,1 | 3046,08 | 3254,21 | 3276,19 | 2 | nd |
| H3N8F3PC2P | 3000,05 | 3056,12 | 3078,1 | 3286,23 | 3308,21 | 1 | nd |
| H3N10FPC2 | 3034,12 | 3090,19 | 3112,17 | 3320,30 | 3342,28 | 1 | nd |
| H3N10PC3 | 3053,12 | 3109,19 | 3131,17 | 3339,30 | 3361,28 | 1 | ✓ |
| H3N8F4PC2 | 3066,14 | 3122,21 | 3144,19 | 3352,32 | 3374,3 | 1 | nd |
| H3N8F3PC3 | 3085,13 | 3141,2 | 3163,18 | 3371,31 | 3393,29 | 2 | nd |
| H3N10PC3P | 3133,09 | 3189,16 | 3211,14 | 3419,27 | 3441,25 | 1 | nd |
| H3N10F4 | 3142,18 | 3198,25 | 3220,23 | | 3450,34 | 2 | nd |
| H3N10FPC3 | 3199,18 | 3255,25 | 3277,23 | 3485,36 | 3507,34 | 1 | ✓ |
| H3N10PC4 | 3218,18 | 3274,25 | 3296,23 | 3504,36 | 3526,34 | 1 | ✓ |
| H3N8F4PC3 | 3231,19 | 3287,26 | 3309,24 | 3517,37 | 3539,35 | 2 | nd |
| H3N8F3PC4 | 3250,19 | 3306,26 | 3328,24 | 3536,37 | 3558,35 | 1 | nd |
| H3N10FPC3P | 3279,14 | 3335,21 | 3357,19 | 3565,32 | 3587,3 | 1 | nd |
| H3N10F5 | 3288,24 | 3344,31 | 3366,29 | | 3596,4 | 1 | nd |
| H3N10F4PC | 3307,24 | 3363,31 | 3385,29 | 3593,42 | 3615,4 | 2 | nd |
| H3N10F2PC3 | 3345,23 | 3401,30 | 3423,29 | 3631,39 | 3653,39 | 1 | ✓ |
| H3N11F4 | 3345,26 | 3401,33 | 3423,31 | | 3653,42 | 1 | nd |
| H3N10FPC4 | 3364,23 | 3420,3 | 3442,28 | 3650,41 | 3672,39 | 1 | ✓ |
| H3N9F4PC3 | 3434,27 | 3490,34 | 3512,32 | 3720,45 | 3742,43 | 1 | nd |
| H3N10F5PC | 3453,27 | 3509,34 | 3531,32 | 3739,45 | 3761,43 | 1 | nd |
| H3N10F2PC4 | 3510,28 | 3566,35 | 3588,33 | 3796,44 | 3818,40 | 1 | ✓ |
| H3N8F4PC5 | 3561,3 | 3617,37 | 3639,35 | 3847,48 | 3869,46 | 1 | nd |
| H3N9F4PC4 | 3599,33 | 3655,4 | 3677,38 | 3885,51 | 3907,49 | 1 | ✓ |
| H3N10F4PC3 | 3637,35 | 3693,42 | 3715,40 | 3923,53 | 3945,51 | 1 | nd |
| H3N10F3PC4 | 3656,35 | 3712,42 | 3734,4 | 3942,53 | 3964,51 | 1 | nd |
| H3N9F4PC5 | 3764,38 | 3820,45 | 3842,43 | 4050,56 | 4072,54 | 1 | ✓ |
| H3N9F4PC6 | 3929,44 | 3985,51 | 4007,49 | 4215,62 | 4237,6 | 1 | ✓ |
| H3N10F5PC4 | 3948,46 | 4004,53 | 4026,51 | 4234,64 | 4256,62 | 1 | ✓ |
| H3N9F4PC7 | 4094,49 | 4150,56 | 4172,54 | 4380,67 | 4402,65 | 1 | ✓ |
| H3N10F5PC6 | 4278,58 | 4334,65 | 4356,63 | 4564,76 | 4586,74 | 1 | ✓ |
| H3N10F5PC7 | 4443,63 | 4499,7 | 4521,68 | 4729,81 | 4751,79 | 1 | ✓ |
| H3N10F5PC8 | 4608,69 | 4664,76 | 4686,74 | 4894,87 | 4916,85 | 1 | ✓ |
| H3N10F5PC9 | 4773,74 | 4829,81 | 4851,79 | 5059,92 | 5081,90 | nd | ✓ |

**Supplementary Table 2: List of proposed *T. suis* pyridylaminated N-glycan structures.** Abbreviated compositions of the form $Hex_{3-5}HexNAc_{2-10}Fuc_{0-5}P_{0-1}PC_{0-8}$ and $Hex_{1-10}HexNAc_{2}Fuc_{0-1}$, Glytoucan accessions (in blue; for defined structures up to 2000 Da), *m/z* as $[M+H]^{+}$, SNFG-style structure, HPLC fraction (either HIAX or, in brackets, HIAX/RP-amide fraction for the second HIAX run) and figure in main text or supplement are given. Structures with a bracket indicate that the exact antennal configuration is not fixed; the HexNAc-substitution of antennal fucose is undefined.

| Composition | PA-[M+H]⁺ | Structure | Fraction | Figure |
|---|---|---|---|---|
| H1N2F1<br>G32152BH | 811.34 | 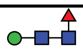 | 6 | |
| H2N2<br>G81315DD | 827.34 | 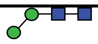 | 8 | |
| H2N2<br>G22573RC | 827.34 | 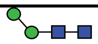 | 9 | |
| H2N2F<br>G42466VF | 973.39 | 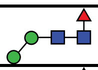 | 8 | |
| H2N2F<br>G00395TQ | 973.39 | 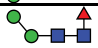 | 10 | |
| H3N2<br>G22768VO | 989.39 | 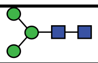 | 11 | |
| H3N2F<br>G45995IV | 1135.45 | 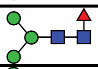 | 12 | |
| H4N2<br>G09724ZC | 1151.44 | 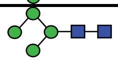 | 15<br>(9-11/11) | |
| H3N3<br>G70073SG | 1192.47 | 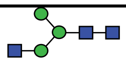 | 14<br>(9-11/09) | |
| H3N3<br>G06920GM | 1192.47 | 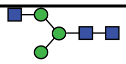 | 16<br>(9-11/15) | |
| H3N2F2<br>G77479VH | 1281.50 | 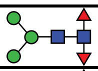 | 13 | |
| H4N2F<br>G48823SZ | 1297.50 | 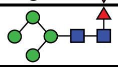 | 16<br>(9-11/15) | |
| H5N2<br>G07617FP | 1313.49 | 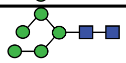 | 19<br>(12+13/6) | |
| H5N2<br>G55220VL | 1313.49 | 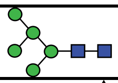 | 21<br>(12+13/07) | |
| H3N3F<br>G14576KZ | 1338.53 | 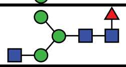 | 15<br>(9-11/14) | |
| H3N3F<br>G69987TD | 1338.53 | 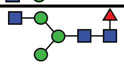 | 16<br>(9-11/19) | |
| H4N3<br>G53168IY | 1354.45 | 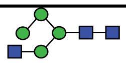 | 18 | |
| H3N4<br>G20956ZK | 1395.55 | 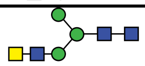 | 19<br>(12+13/05) | |
| H3N4<br>G89096IL | 1395.55 | 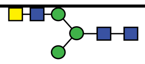 | 20<br>(12+13/11) | |
| H3N4<br>G39213VZ | 1395.55 | 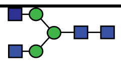 | 18<br>(12+13/08) | |
| H3N4<br>G18119VX | 1395.55 | 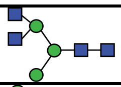 | 17 | |
| H6N2<br>G61846BY | 1475.55 | 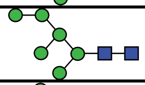 | 24/25<br>(14-16/07) | |
| H6N2<br>G80966KZ | 1475.55 | 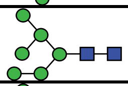 | 24/25<br>(14-16/08) | |
| H6N2<br>G34499SX | 1475.55 | 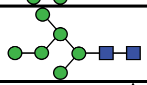 | 24/25<br>(14-16/12) | |
| H3N3F2<br>G41720HN | 1484.58 | 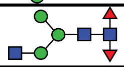 | 16<br>(9-11/8) | |

| | | | | |
|---|---|---|---|---|
| H3N3F2<br>G10425SO | 1484.58 | 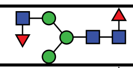 | 18 | |
| H4N3F<br>G58969RU | 1500.58 | 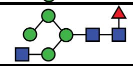 | 19<br>(12+13/09) | |
| H5N3<br>G08520NM | 1516.57 | 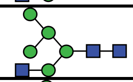 | 23<br>(14-16/09) | |
| H3N4F<br>G80019MD | 1541.60 | 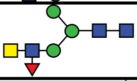 | (14-16/06) | |
| H3N4F<br>G20956ZK | 1541.60 | 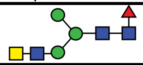 | 19<br>(12+13/10) | |
| H3N4F<br>G63770IR | 1541.60 | 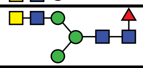 | 21<br>(12+13/15) | |
| H3N4F<br>G80858MF | 1541.60 | 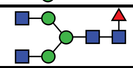 | 20<br>(12+13/11) | |
| H3N4F<br>G92571WK | 1541.60 | 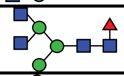 | 19<br>(12+13/17) | |
| H4N4<br>G59231NT | 1557.60 | 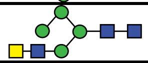 | (14-16/10) | |
| H3N4PC<br>G16427JQ | 1560.60 | 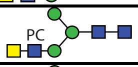 | 27 | |
| H3N4PC<br>G37840NY | 1560.60 | 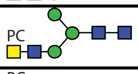 | 29<br>(17-20/10) | |
| H3N4PC<br>G87872NF | 1560.60 | 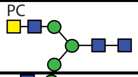 | 29<br>(17-20/17) | |
| H3N5<br>G67045LB | 1598.63 | 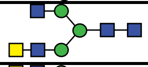 | (14-16/13) | |
| H3N5<br>G19868AX | 1598.63 | 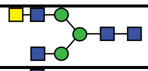 | 23<br>(14-16/14) | |
| H3N5 | 1598.63 | 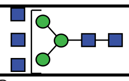 | 22<br>(14-16/15) | |
| H3N4FP | 1621.57 | 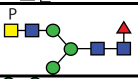 | 35, 36 | Fig. S10 |
| H7N2<br>G83161QT | 1637.60 | 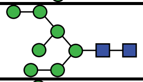 | 28<br>(17-20/06) | |
| H7N2<br>G68668TB | 1637.60 | 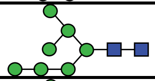 | 29<br>(17-20/08) | |
| H7N2<br>G63337SS | 1637.60 | 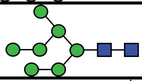 | 28/29<br>(17-20/11) | |
| H3N4F2<br>G84692YW | 1687.66 | 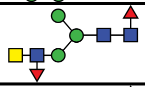 | 25<br>(14-16/11) | Fig. 4B<br>Fig. S9B |
| H3N4F2<br>G47253EY | 1687.66 | 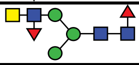 | 26<br>(14-16/16) | Fig. S9A |
| H3N4F2<br>G69912LP | 1687.66 | 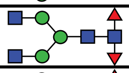 | 20<br>(12+13/07) | Fig. 4A |
| H4N4F<br>G45048TZ | 1703.66 | 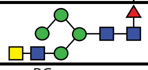 | (14-16/14) | |
| H3N4FPC<br>G04635TH | 1706.66 | 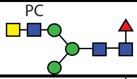 | 27 | Fig. S9C |
| H3N4FPC<br>G16427JQ | 1706.66 | 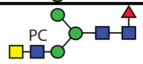 | 28 | |
| H3N4FPC<br>G67053ZX | 1706.66 | 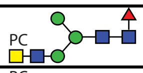 | 29<br>(17-20/14) | |
| H3N4FPC<br>G03090TG | 1706.66 | 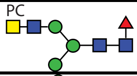 | 30<br>(17-20/21) | Fig. S9C |
| H3N4FPC<br>G90339LD | 1706.66 | 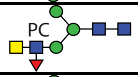 | (17-20/06) | Fig. S9C |
| H4N4PC | 1722.66 | 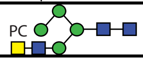 | 32 | |

| | | | | |
|---|---|---|---|---|
| H3N5F<br>G88461FF | 1744.68 | 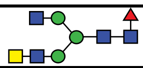 | 23<br>(14-16/17) | |
| H3N5F<br>G82312JY | 1744.68 | 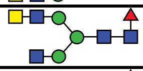 | 24<br>(14-16/18) | |
| H3N5F<br>G61207RZ | 1744.68 | 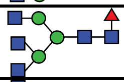 | (14-16/19) | |
| H3N5F<br>G80630NF | 1744.68 | 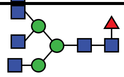 | (14-16/24) | |
| H3N5PC | 1763.68 | 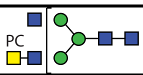 | 32 | |
| H8N2<br>G40702WU | 1799.65 | 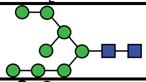 | 32 | |
| H8N2<br>G89864BN | 1799.65 | 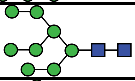 | 33 | |
| H8N2<br>G91704UR | 1799.65 | 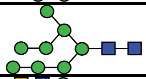 | 34 | |
| H3N6<br>G09482NW | 1801.71 | 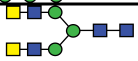 | 28<br>(17-20/16) | Fig. 4C |
| H3N6 | 1801.71 | 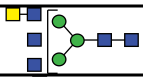 | 27<br>(14-16/22) | |
| H3N6<br>G90223GH | 1801.71 | 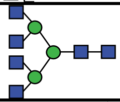 | 25<br>(14-16/23) | Fig. 4D |
| H3N4F2PC<br>G13755PW | 1852.72 | 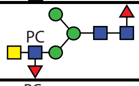 | 31 | Fig. S10A<br>Fig. S13 |
| H3N4F2PC<br>G21650ZZ | 1852.72 | 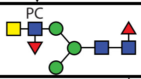 | (17-20/16) | |
| H3N4F2PC<br>G23967QF | 1852.72 | 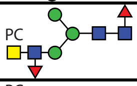 | 35 | |
| H3N4F2PC<br>G64990BS | 1852.72 | 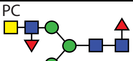 | 36 | Fig. S10E |
| H3N5F2<br>G30205VB | 1890.74 | 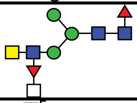 | 26<br>(17-20/12) | Fig. 6A |
| H3N5F2 | 1890.74 | 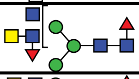 | 27<br>(17-20/17) | Fig. 4H |
| H3N5F2<br>G55860VC | 1890.74 | 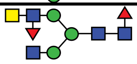 | 29<br>(17-20/15) | Fig. 4G |
| H3N5FPC | 1909.74 | 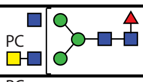 | 32 | |
| H3N5FPC | 1909.74 | 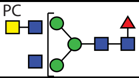 | 34 | |
| H3N6F<br>G85542KD | 1947.76 | 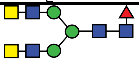 | (17-20/22) | |
| H3N6F | 1947.76 | 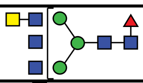 | 27 | |
| H3N6F<br>G59866SG | 1947.76 | 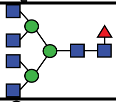 | 26<br>(17-20/26) | |
| H9N2<br>G60230HH | 1961.71 | 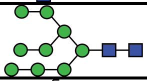 | 37 | |
| H3N7 | 2004.79 | 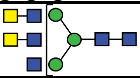 | (17-20/19) | Fig. 4E |
| H3N7 | 2004.79 | 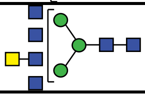 | (17-20/24) | Fig. 4F |

| | | | | |
|---|---|---|---|---|
| H3N4F2PC2 | 2017.77 | 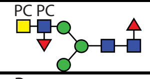  | 40 | |
| H3N6PCP | 2046.73 | 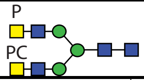  | 44+45<br>46+47 | Fig. S17F |
| H3N5F2PC | 2055.80 | 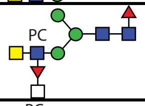  | 31, 32 | Fig. 6B<br>Fig. S13 |
| H3N5F2PC | 2055.80 | 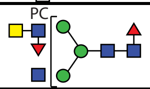  | 34 | |
| H3N5F2PC | 2055.80 | 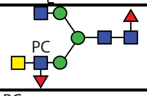  | 36 | |
| H3N5F2PC | 2055.80 | 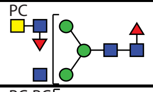  | 38 | Fig. S13 |
| H3N5FPC2 | 2074.80 | 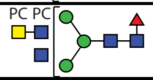  | 39 | |
| H3N6F2 | 2093.82 | 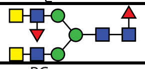 | 39 | |
| H3N6FPC | 2112.82 | 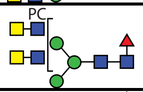  | 34 | |
| H3N6FPC | 2112.82 | 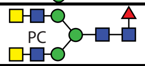  | 35 | |
| H10N2<br>G19958IL | 2123.76 | 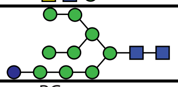 | 40 | |
| H3N6PC2 | 2131.82 | 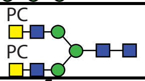  | 44+45 | Fig. S17G |
| H3N7F | 2150.84 | 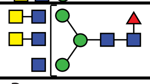 | 31 | |
| H3N6FPCP | 2192.79 | 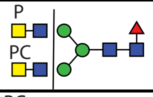  | 44+45 | Fig. S17A |
| H3N6FPCP | 2192.79 | 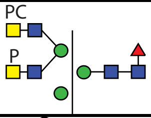  | 46+47 | Fig. S17D |
| H5N4FPC2 | 2196.00 | 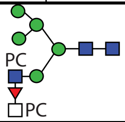  | 46+47 | Fig. 6G |
| H3N5F2PC2 | 2220.85 | 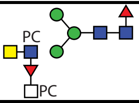  | 40 | Fig. 6C |
| H3N6F3 | 2239.88 | 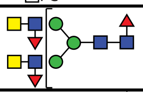 | 39 | Fig. 4I |
| H3N6F2PC | 2258.88 | 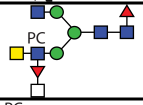  | 36 | |
| H3N6F2PC | 2258.88 | 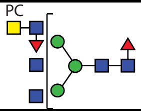  | 38 | |
| H3N6F2PC | 2258.88 | 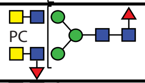  | 42 | |
| H3N6F2PC | 2258.88 | 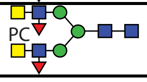  | 46+47 | |

| | | | | |
|---|---|---|---|---|
| H3N6FPC2 | 2277.87 | 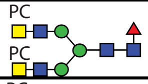 | 44+45 | Fig. S17C |
| H3N6FPC2 | 2277.87 | 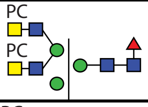 | 44+45 | Fig. S16A |
| H3N6FPC2 | 2277.87 | 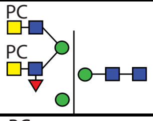 | 46+47 | Fig. S17E |
| H3N7PC2 | 2334.90 | 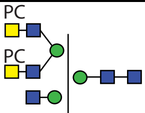 | 46+47 | |
| H3N6F3PC | 2404.93 | 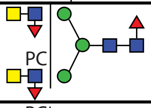 | 43, 44+45 | Fig. 6E |
| H3N6F2PC2 | 2423.93 | 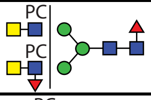 | 44+45 | |
| H3N6F2PC2 | 2423.93 | 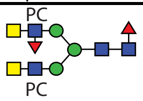 | 44+45 | Fig. 5E |
| H3N6F2PC2 | 2423.93 | 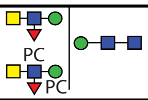 | 46+47 | Fig. 7A |
| H3N6F2PC2 | 2423.93 | 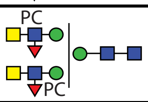 | 46+47 | Fig. 7B |
| H3N6F2PC2 | 2423.93 | 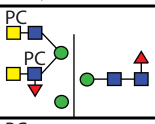 | 46+47 | Fig. 5D |
| H3N6F2PC2 | 2423.93 | 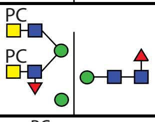 | 46+47 | Fig. 5F |
| H3N6F2PC2 | 2423.93 | 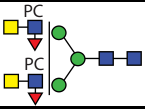 | 48 | Fig. 5B |
| H3N6F2PC2 | 2423.93 | 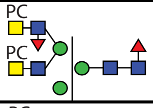 | 48 | |
| H3N6F2PC2 | 2423.93 | 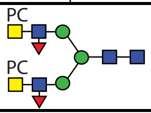 | 49+50 | Fig. 5A |
| H3N6F2PC2 | 2423.93 | 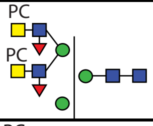 | 49+50 | Fig. 5C |
| H3N6FPC3 | 2442.93 | 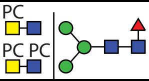 | 48<br>49+50 | Fig. 5G |
| H3N7F3 | 2442.96 | 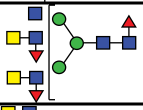 | 39 | |
| H3N7FPC2 | 2480.95 | 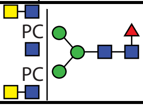 | 46+47 | |

| | | | | |
|---|---|---|---|---|
| H3N7FPC2 | 2480.95 | 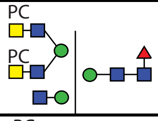 | 46+47 | |
| H3N8FPC | 2518.98 | 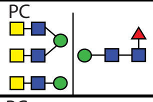 | 44+45 | |
| H3N8PC2 | 2537.98 | 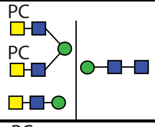 | 48<br>49+50 | |
| H3N8PC2 | 2537.98 | 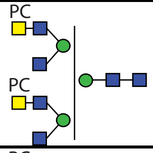 | 48 | |
| H3N8PC2 | 2537.98 | 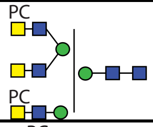 | 49+50 | |
| H3N6F3PC2 | 2569.99 | 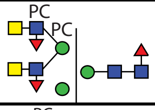 | 44+45<br>46+47 | Fig. 7C |
| H3N6F3PC2 | 2569.99 | 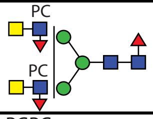 | 48<br>49+50 | Fig. 5H |
| H3N6F3PC2 | 2569.99 | 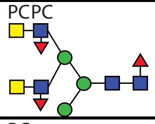 | 49+50 | Fig. 5I |
| H3N6F3PC2 | 2569.99 | 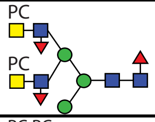 | 49+50 | Fig. 5J |
| H3N6F3PC2 | 2569.99 | 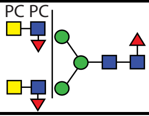 | 51+52 | |
| H3N9PC | 2576.12 | 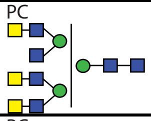 | 44+45 | |
| H3N9PC | 2576.12 | 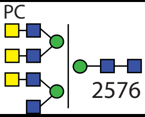 | 44+45 | |
| H3N6F2PC3 | 2588.99 | 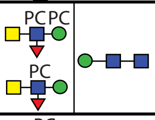 | 49+50 | Fig. 7D |
| H3N6F2PC3 | 2588.99 | 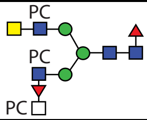 | 51+52 | |
| H3N6F2PC3 | 2588.99 | 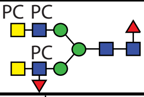 | 51+52 | |
| H3N7F3PC | 2608.09 | 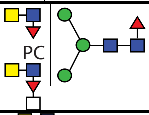 | 44+45 | Fig. 6D |
| H3N10<br>G53059NZ | 2614.02 | 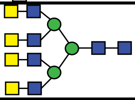 | 41 | |

| | | | | |
|---|---|---|---|---|
| H3N7F2PC2 | 2627.01 | 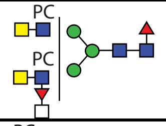 | 44+45 | |
| H3N7F2PC2 | 2627.01 | 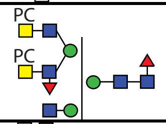 | 49+50 | |
| H3N7F2PC2 | 2627.01 | 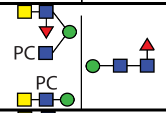 | 48 | |
| H3N8F3 | 2646.04 | 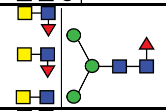 | 44+45 | |
| H3N8F2PC | 2665.12 | 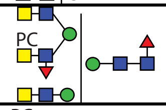 | 46+47 | |
| H3N8FPC2 | 2684.03 | 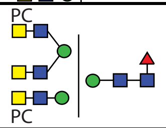 | 46+47 | Fig. S14A |
| H3N8FPC2 | 2684.03 | 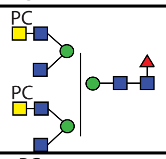 | 49+50 | Fig. S14C |
| H3N8FPC2 | 2684.03 | 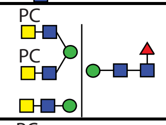 | 48<br>49+50 | Fig. S14B |
| H3N9FPC | 2722.22 | 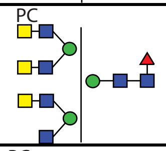 | 46+47 | |
| H3N9FPC | 2722.22 | 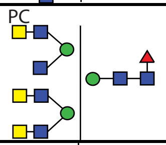 | 46+47 | |
| H3N6F3PC3 | 2735.05 | 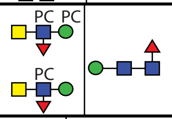 | 48<br>49+50 | Fig. 7E |
| H3N6F3PC3 | 2735.05 | 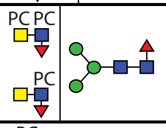 | 56+57 | |
| H3N9PC2 | 2741.05 | 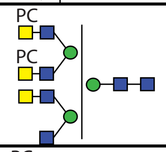 | 49+50<br>51+52 | |
| H3N9PC2 | 2741.05 | 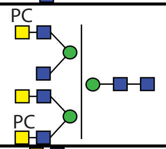 | 49+50<br>51+52 | |
| H3N10F<br>G51374ZY | 2760.08 | 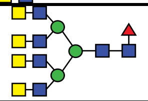 | 43 | |

| | | | | |
|---|---|---|---|---|
| H3N7F3PC2 | 2773.07 | 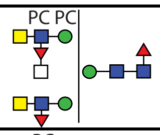 | 46+47<br>51+52 | Fig. 7F |
| H3N7F3PC2 | 2773.07 | 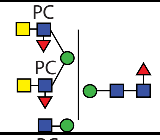 | 46+47 | Fig. S14F |
| H3N7F3PC2 | 2773.07 | 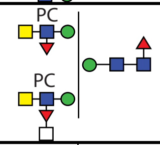 | 48<br>49+50 | Fig. S14E<br>Fig. S15D |
| H3N7F3PC2 | 2773.07 | 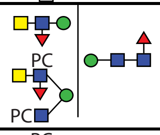 | 51+52 | Fig. S14D |
| H3N7F3PC2 | 2773.07 | 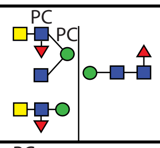 | 51+52 | |
| H3N10PC | 2779.08 | 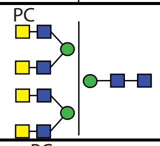 | 48 | |
| H3N7F2PC3 | 2792.08 | 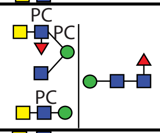 | 51+52 | |
| H3N8F4 | 2792.10 | 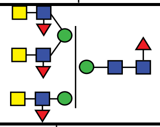 | 48 | Fig. 4J |
| H3N8F4 | 2792.10 | 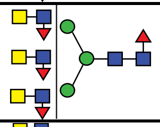 | 49+50 | |
| H3N8F3PC | 2811.09 | 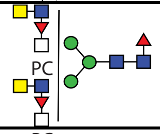 | 44+45 | |
| H3N8FPC3 | 2849.09 | 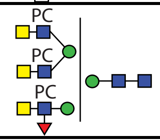 | 51+52 | |
| H3N8FPC3 | 2849.09 | 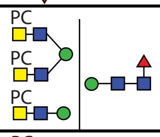 | 53-55 | Fig. S16B |
| H3N8FPC3 | 2849.09 | 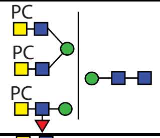 | 56+57 | |
| H3N9F3 | 2849.12 | 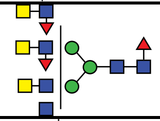 | 44+45 | |
| H3N9F3 | 2849.12 | 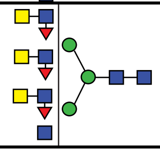 | 48 | |

| | | | | |
|---|---|---|---|---|
| H3N9FPC2 | 2887.11 | 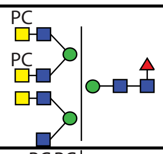 | 51+52 | |
| H3N6F3PC4 | 2900.10 | 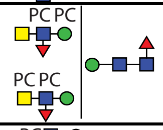 | 53-55 | |
| H3N6F3PC4 | 2900.10 | 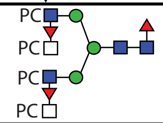 | 58+59 | Fig. 6H |
| H3N9PC3 | 2906.11 | 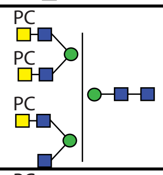 | 53-55 | |
| H3N10FPC | 2925.14 | 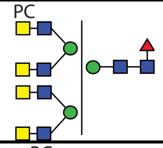 | 48 | |
| H3N7F3PC3 | 2938.13 | 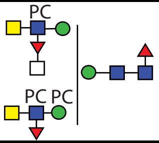 | 48 | Fig. 7G |
| H3N8F5 | 2938.13 | 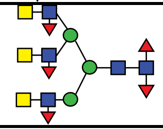 | 48 | Fig. 4K |
| H3N8F4PC | 2957.15 | 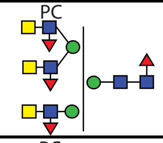 | 51+52 | |
| H3N8F3PC2 | 2976.15 | 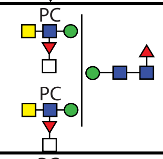 | 49+50 | Fig. S15C |
| H3N8F3PC2 | 2976.15 | 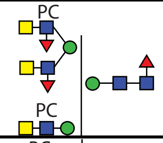 | 51+52 | |
| H3N8F3PC2 | 2976.15 | 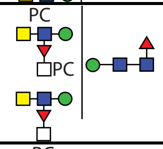 | 51+52 | |
| H3N8F3PC2 | 2976.15 | 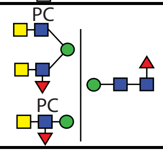 | 51+52 | |
| H3N8F3PC2 | 2976.15 | 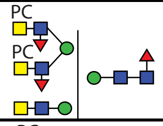 | 53-55 | |
| H3N8F2PC3 | 2995.15 | 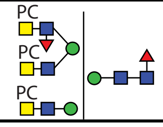 | 56+57 | Fig. S14J |

| | | | | |
|---|---|---|---|---|
| H3N8F2PC3 | 2995.15 | 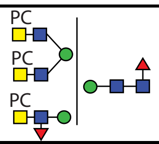 | 56+57 | |
| H3N9F4 | 2995.18 | 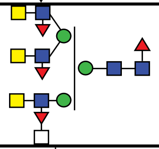 | 48<br>49+50 | Fig. S14G |
| H3N9F4 | 2995.18 | 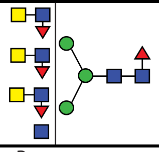 | 48<br>49+50<br>51+52 | Fig. S14I |
| H3N10PC2P | 3024.10 | 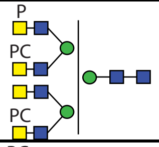 | 56+57 | |
| H3N10PC2P | 3024.10 | 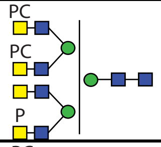 | 56+57 | Fig. S17H |
| H3N8F3PC2P | 3056.12 | 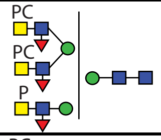 | 56+57 | |
| H3N10FPC2 | 3090.19 | 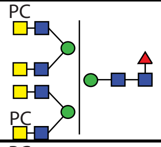 | 51+52 | |
| H3N10PC3 | 3109.19 | 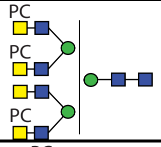 | 56+57 | Fig. S17I |
| H3N8F4PC2 | 3122.21 | 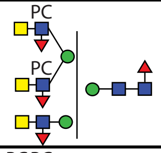 | 53-55 | |
| H3N8F4PC2 | 3122.21 | 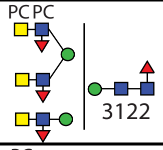 | 56+57 | |
| H3N8F3PC3 | 3141.20 | 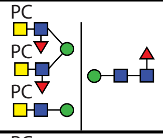 | 58+59 | |
| H3N8F3PC3 | 3141.20 | 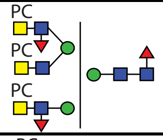 | 58+59 | |
| H3N10PC3P | 3189.16 | 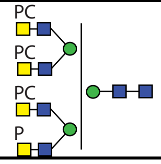 | 58+59<br>60+61 | Fig. S11 |

| | | | | |
|---|---|---|---|---|
| H3N10F4 | 3198.25 | 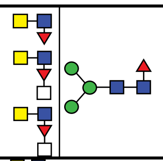 | 49+50 | |
| H3N10F4 | 3198.25 | 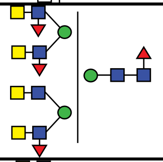 | 51+52 | |
| H3N10F4 | 3198.25 | 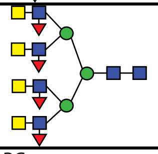 | 53-55 | |
| H3N10FPC3 | 3255.25 | 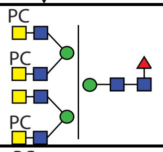 | 56+57 | Fig. S16C |
| H3N10PC4 | 3274.25 | 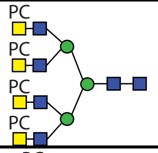 | 60+61 | Fig. S11 |
| H3N8F4PC3 | 3287.26 | 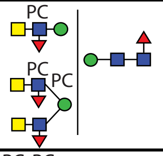 | 56+57 | Fig. S16F |
| H3N8F4PC3 | 3287.26 | 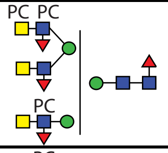 | 56+57<br>58+59 | Fig. S16G |
| H3N8F3PC4 | 3306.26 | 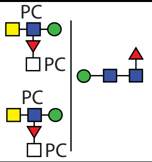 | 56+57 | |
| H3N10FPC3P | 3335.21 | 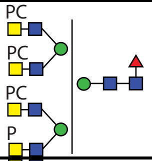 | 58+59 | Fig. S17J |
| H3N10F5<br>G41784WX | 3344.31 | 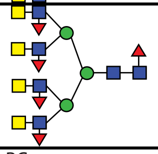 | 53-55 | Fig. 4L |
| H3N10F4PC | 3363.31 | 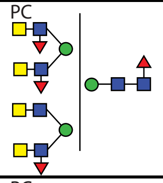 | 58+59 | |
| H3N10F4PC | 3363.31 | 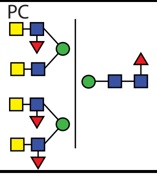 | 58+59 | |

| | | | | |
|---|---|---|---|---|
| H3N10F2PC3 | 3401.30 | 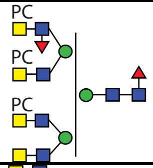 | 58+59 | Fig. S16D |
| H3N11F4 | 3401.33 | 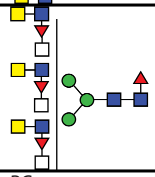 | 49+50 | Fig. S16E |
| H3N10FPC4 | 3420.30 | 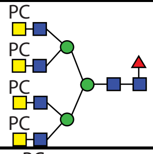 | 60+61 | Fig. S11 |
| H3N9F4PC3 | 3490.34 | 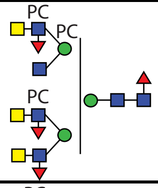 | 56+57 | Fig. S16H |
| H3N9F4PC3 | 3490.34 | 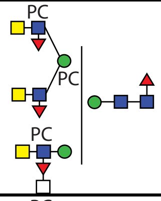 | 56+57 | Fig. S16I |
| H3N10F5PC | 3509.34 | 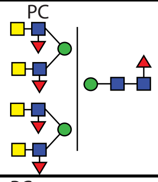 | 56+57 | |
| H3N10F5PC | 3509.34 | 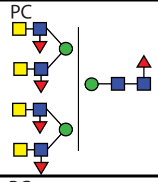 | 58+59 | |
| H3N10F2PC4 | 3566,35 | 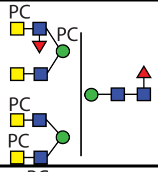 | 61 | |
| H3N8F4PC5 | 3617.37 | 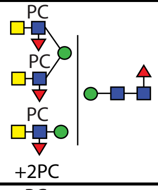 | 66 | |
| H3N9F4PC4 | 3655.40 | 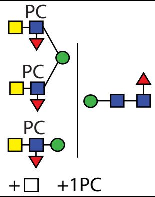 | 62 | |

| | | | | |
|---|---|---|---|---|
| H3N10F4PC3 | 3693.42 | 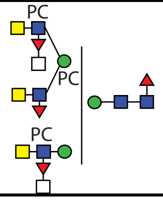 | 56+57 | |
| H3N10F3PC4 | 3712.42 | | 65 | |
| H3N9F4PC5 | 3820.45 | | 66 | |
| H3N9F4PC6 | 3985.51 | 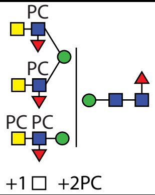 | 71 | |
| H3N10F5PC4 | 4004.53 | | 67 | |
| H3N9F4PC7 | 4150.56 | | 73 | |
| H3N10F5PC6 | 4334.65 | | 73 | |
| H3N10F5PC7 | 4499.70 | 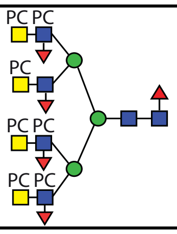 | 75 | Fig. S12 |
| H3N10F5PC8 | 4664.76 | | 75 | |
