## Supplementary figures and images for "Recognition of highly branched N-glycans of the porcine whipworm by the immune system"

### Supplementary Figure 18

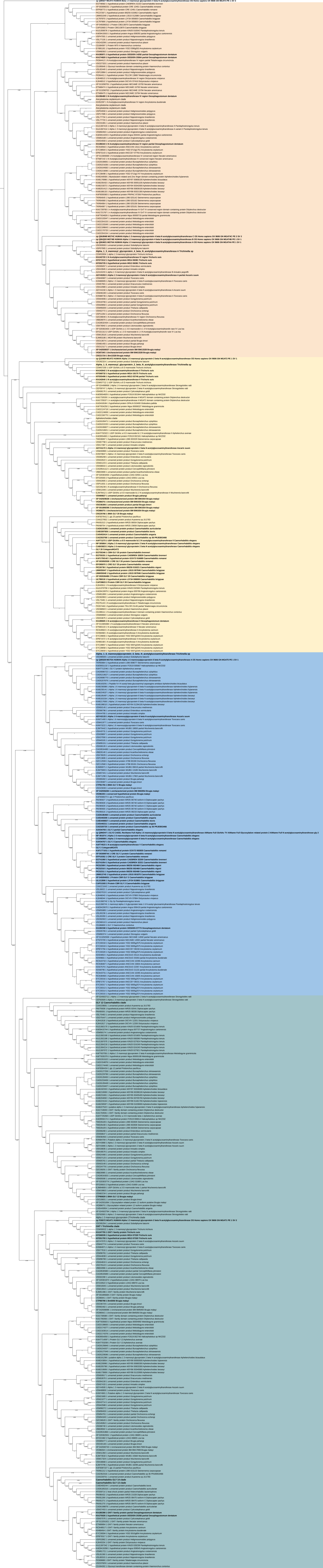
